# A noise-corrected metric for evaluating spatial transcriptomics predicted from histology

**DOI:** 10.64898/2026.11.24.768840

**Authors:** Jiarui Ouyang, Yihang Gao, Cheng Jin, Yihui Wang, Yingxue Xu, Ling Liang, Fengtao Zhou, Zhiyuan Cai, Hao Chen

## Abstract

Deep learning methods that predict spatial transcriptomics from histology offer scalable molecular profiling of clinical cohorts. However, conventional correlation metrics overstate prediction quality across whole sections by crediting models for recognizing broad tissue regions, and understate fine-scale prediction quality because sampling noise dominates sparse RNA counts. To resolve this, we introduce the reproducible-signal correlation (RSC) metric. RSC randomly splits RNA counts into two halves to separate reproducible signal from sampling noise across spatial scales and within tissue regions or cell classes. Across 12 spot-level and three single-cell methods, RSC revealed that most models capture broad tissue patterns but largely miss differences between neighboring spots or cells, which whole-section correlation overlooks. Within malignant cells, RSC distinguished methods that genuinely capture part of the cell-to-cell heterogeneity from those whose scores across all cells were inflated by cell-type composition. Guided by RSC, we designed a pairwise difference loss between neighboring spots that improved fine-scale predictions on average across 11 models. RSC thus provides a foundation for benchmarking predictions, guiding model design and determining which spatial analyses histology can support.

## Introduction

Spatial transcriptomics maps gene expression within tissue architecture, revealing molecular organization across broad tissue compartments, local cellular neighborhoods and individual cells^1–3^. Deep learning methods that infer spatial expression profiles directly from routine hematoxylin and eosin (H&E) histology promise to extend molecular profiling to large clinical cohorts and archival biobanks that lack matched RNA measurements^4,5^. Recent deep learning methods have pushed nominal output resolution from multicellular tissue spots down to single cells^4–7^, aiming to resolve cellular interactions beyond broad anatomical compartments. Yet high nominal resolution is only meaningful if predictions faithfully capture expression differences at the corresponding fine scales, such as variation across cellular neighborhoods, between adjacent spots or among individual cells of the same class. Determining whether a method resolves expression at these fine scales requires evaluation metrics that remain valid across spatial distances, a standard that current evaluation practices fail to meet.

In practice, current evaluations treat the RNA counts measured in spots or cells as the ground truth and correlate predictions with the measured counts across whole tissue sections^5,8–10^. However, assessing prediction quality with conventional whole-section correlation is fundamentally distorted in two opposing ways. First, whole-section correlation is dominated by macroscopic differential expression between broad anatomical compartments such as tumor nests and stroma, which overstates prediction quality for analyses within individual regions or cell classes (Fig. 1b). A model can attain a deceptively high whole-section correlation simply by recognizing coarse tissue architecture, even while its predictions remain flat, uninformative or falsely ordered within an individual region. As a result, a naive class-mean baseline that assigns constant average expression to each region can score above the median of 11 published methods on whole sections without learning any local variation whatsoever. Second, attempting to evaluate predictions at fine spatial scales directly against raw measured counts systematically understates prediction quality because sparse transcript counts are subject to substantial Poisson sampling noise (Fig. 1c). Single-cell studies have established that random molecule capture introduces stochastic sampling fluctuations that intensify as transcript counts become sparse^11,12^. Because transcript counts are sparsest in individual cells and localized spots^13,14^, this measurement noise severely attenuates correlation with measured counts, making genuine model errors indistinguishable from unavoidable assay noise. Consequently, current evaluations overstate predictive success within tissue regions while leaving fine-scale predictive capacity unresolved.

**Figure 1.**
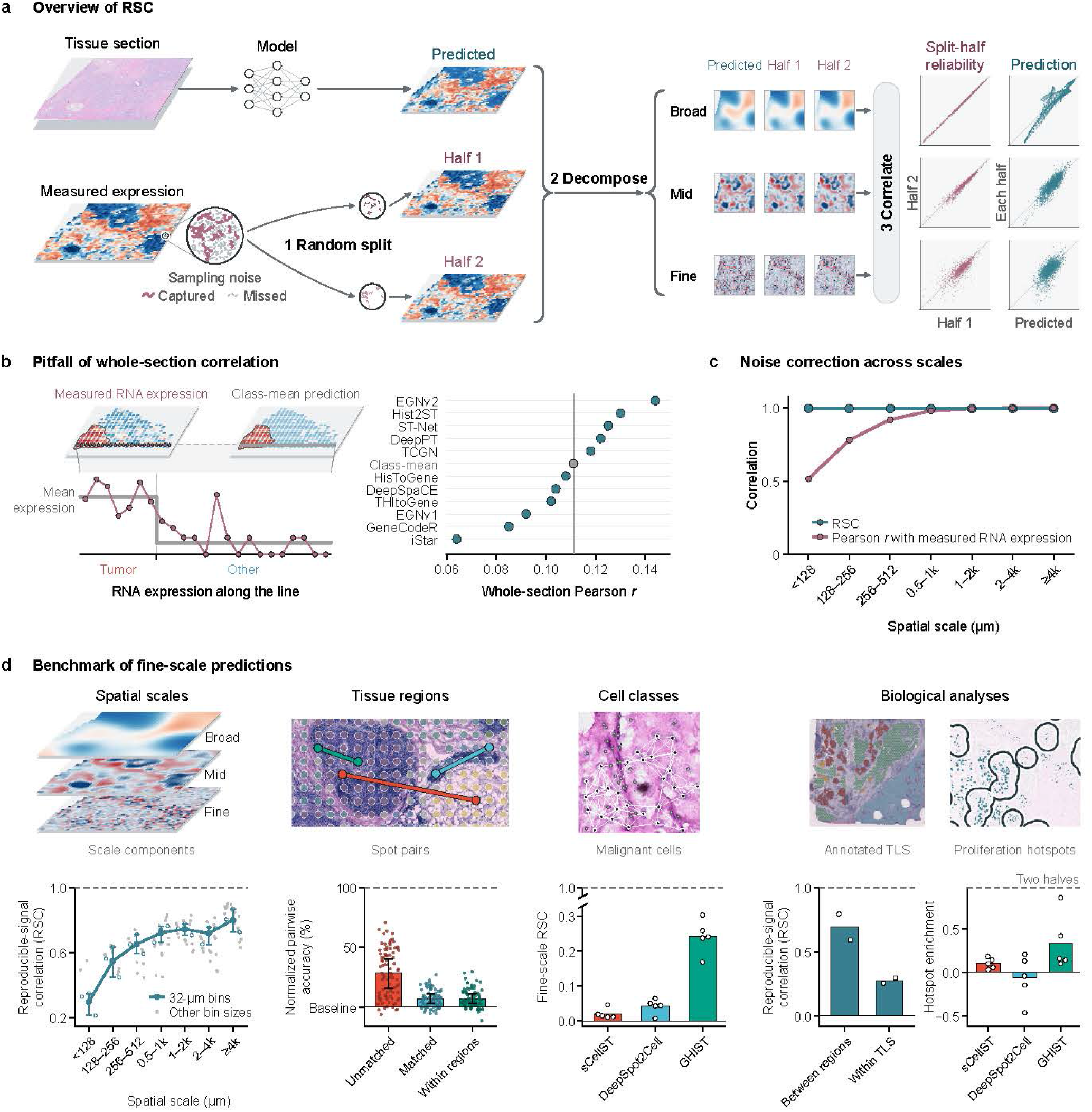
Reproducible-signal correlation (RSC) largely avoids overstating and understating prediction quality. a, RSC framework overview: (1) random split partitions measured transcripts into two halves with independent Poisson noise; (2) spatial multiscale decomposition isolates broad, mid and fine scale components; (3) prediction correlation with each half is divided by the square root of split-half reliability (Methods). **b,** Pitfall of whole-section correlation. Expression transect across tumor and non-tumor regions (left) and whole-section Pearson *r* across 785 genes in HER2ST (*n* = 8 patients; right). A naive class-mean baseline with zero intra-regional variation scores above the median of 11 published methods (*r* = 0.111 versus 0.108), demonstrating that macroscopic differences can dominate conventional evaluation. **c,** Noise correction across scales. Conventional Pearson *r* with measured RNA expression (pink) drops at fine scales due to Poisson noise attenuation, whereas RSC (teal) remains stable near 1.0 across scales (*<* 128 *µ*m to ≥4k *µ*m; ideal prediction matching the underlying expression in three colorectal cancer patients). **d,** Benchmark summary across four axes (Figs. 2–5): spot-level Ridge–Phikon predictions fall to an RSC of 0.298 at the finest scale despite high broad-scale RSC (*spatial scales*; Fig. 3); normalized pairwise accuracy drops toward the majority-order baseline (0%) once spot pairs are matched or evaluated within regions (*tissue regions*; Fig. 2); fine-scale RSC within malignant cells removes cell-type confounding to separate single-cell methods (*cell classes*; Fig. 4); and models capture inter-region differences but largely miss expression variation within TLS, while single-cell models diverge in proliferation hotspot enrichment (*biological analyses*; Fig. 4e and Fig. 5f). Sample sizes and statistics are given in Figs. 2–5. TLS, tertiary lymphoid structures.

To address both problems, an evaluation metric should isolate local spatial variation from differential expression between tissue regions and separate reproducible biological signal from sampling noise in the measured RNA. Here we introduce reproducible-signal correlation (RSC), a metric that corrects noise-induced understatement across spatial scales and avoids overstatement within tissue regions or cell classes (Fig. 1). Drawing on Fourier shell correlation in cryo-electron microscopy and Fourier ring correlation in optical nanoscopy^15–18^, RSC randomly partitions captured RNA molecules into two halves through binomial count splitting^19–21^. Both halves share identical underlying biological expression rates while carrying statistically independent Poisson sampling noise. The correlation between the two halves measures the reproducible biological signal in the measured RNA at each spatial scale, from broad tissue compartments to local neighborhoods and individual spots or cells. Because sampling noise is random, a prediction can correlate with the measured RNA only through this signal, which sets a noise ceiling on the correlation^22^. RSC then evaluates a prediction by dividing its correlation with the RNA halves by this ceiling^23–25^. An RSC of 1.0 therefore indicates that a prediction accounts for all reproducible biological variation accessible at that scale, whereas an RSC of 0 reflects chance agreement. For RNA counts simulated with known expected expression, RSC correctly scored the true expression near 1 across spatial scales, whereas conventional correlation with counts fell toward fine scales (Fig. 1c; Extended Data Fig. 1c). When computed separately within each tissue region or cell class, RSC also distinguishes predictions that capture local spot-to-spot differences and cell-to-cell variability from coarse region-mean predictions such as the class-mean baseline.

Applied to 12 spot-level and three single-cell prediction methods, our evaluations showed that most of these methods capture broad-scale signal such as differential expression between tissue regions or cell classes but little fine-scale signal (Fig. 1d; Extended Data Table 1). At fine scales of individual spots and local neighborhoods, RSC showed that spot-level methods largely missed the reproducible signal that whole-section correlation overlooks. Yet at single-cell resolution, RSC separated genuine within-class resolving power from cell-type composition, identifying GHIST^5^ as the evaluated method that recovers the most cell-to-cell variability among malignant cells from H&E images. In downstream biological analyses, a spot-level model accurately located tertiary lymphoid structures, but RSC showed that it poorly predicted the expression variation inside these structures. Support for predicted proliferation hotspots varied markedly across single-cell methods, and GHIST located these hotspots best among the three methods. Finally, guided by the RSC principle, we designed a training loss that supervises how expression changes between neighboring spots. The loss improved predictions on average across 11 models in both fine-scale RSC and whole-section correlation. RSC therefore shows which analyses histology-based predictions can support and where methods need improvement. Researchers can evaluate their own methods on an interactive website (https://stRSC.org), which also displays the evaluations of the datasets in this study.

## Results

### Whole-section correlation overstates prediction quality within tissue regions

Histology-based spatial transcriptomics prediction models are routinely evaluated by computing the correlation between predicted and measured expression across all spots in an entire tissue section^5,8,9^. However, whole-section correlation pools spots across distinct anatomical compartments and is often dominated by large macroscopic expression differences between regions. As a result, whole-section correlation credits models for recognizing coarse tissue architecture while obscuring whether they resolve local spot-to-spot expression differences within individual regions. To demonstrate how this pooling distorts evaluation, we examined the held-out predictions of 11 published methods^8^ across 785 genes in HER2ST, a dataset of HER2-positive breast cancer sections with pathologist-annotated histological regions (Fig. 1b; Fig. 2; Supplementary Table 1)^26^. We compared these 11 methods against a naive class-mean baseline that distinguishes only tumor from non-tumor regions and assigns identical expression to every spot within the same region. Although this baseline captures no spot-to-spot differences within regions, whole-section Pearson correlation scored it above the median of the 11 published methods (Pearson *r* of 0.111 versus a median of 0.108 and a maximum of 0.144; Fig. 1b; Supplementary Note 1). Whole-section correlation alone therefore failed to distinguish a model that captures genuine local variation from one that merely outputs a constant regional average.

**Figure 2.**
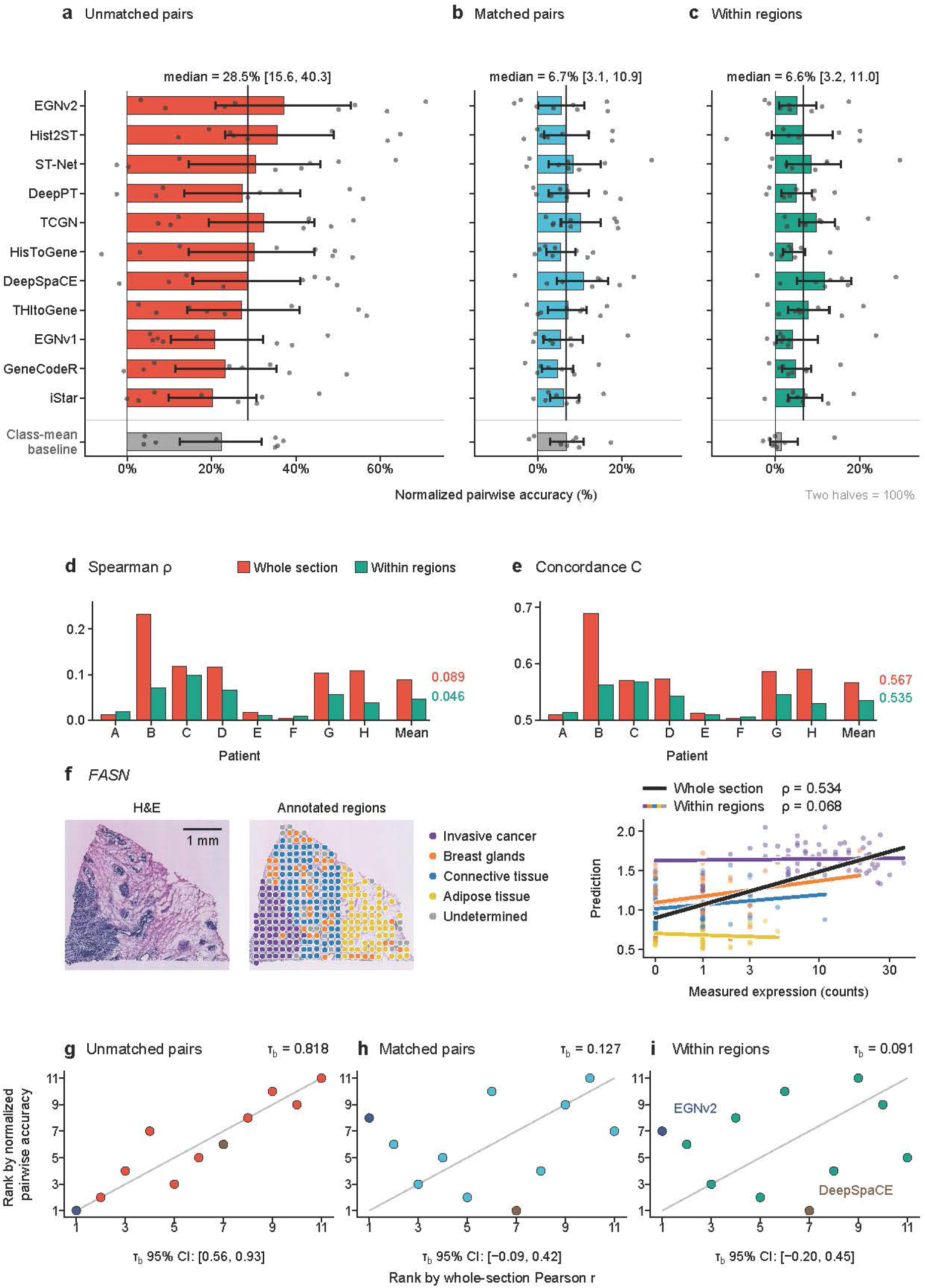
Evaluation within tissue regions reveals whole-section correlation overstates prediction quality. a–c, Normalized pairwise accuracy of 11 published spot-level methods and the class-mean baseline (gray) across 785 genes in HER2ST (*n* = 8 patients). Spot pairs are unmatched (**a**), matched for expression profile and sequencing depth (**b**) or matched strictly within the same histological region (**c**). Normalized accuracy is scaled between a majority-order baseline (0%) and split-half agreement (100%); vertical lines mark medians, bars show method means, and points show patients with 95% bootstrap intervals. Performance collapses from a median of 28.5% (unmatched) to 6.7% (matched) and 6.6% (within regions). **d,e,** Gene-wise Spearman correlation (*ρ*; **d**) and all-pair concordance (*C*; **e**) for patients A–H, showing drops from whole sections to within-region evaluations (mean *ρ*: 0.089 to 0.046; mean *C*: 0.567 to 0.535). **f,** Hist2ST prediction of *FASN* in an example patient section (patient B). Although Hist2ST mirrors coarse tumor morphology (whole-section *ρ* = 0.534), it largely fails to resolve expression variation within regions (*ρ* = 0.068). Scale bar, 1 mm. **g–i,** Method rankings by whole-section Pearson *r* versus normalized pairwise accuracy for unmatched pairs (**g**; Kendall’s *τ_b_* = 0.818), matched pairs (**h**; *τ_b_* = 0.127) and within regions (**i**; *τ_b_* = 0.091), demonstrating that controlling for region-level expression reshuffles method rankings in HER2ST. The 95% confidence intervals are shown.

Consequently, whole-section correlation substantially overstated the prediction quality of the 11 published methods for analyses within tissue regions, where all methods scored much lower than across whole sections (Fig. 2a–f). This overstatement is exemplified by individual marker genes such as *FASN*, for which Hist2ST predictions correlated with measured RNA at Spearman *ρ* = 0.534 across all spots of a section but collapsed to *ρ* = 0.068 among spots within individual regions (Fig. 2f). To extend this comparison across genes, we quantified how often a prediction correctly identifies which of two spots has higher measured expression. Normalized pairwise accuracy scales this ordering fraction from 0% for a majority-order baseline to 100% for the empirical agreement between two random halves of the measured RNA counts. The majority-order baseline assigns all spot pairs of a gene the relative direction observed most frequently across training patients. We compared three sets of spot pairs that progressively remove differential expression between regions. Unmatched pairs were sampled randomly across the entire section, matched pairs had similar total transcript counts and overall expression profiles, and within-region pairs were matched pairs in the same pathologist-annotated region. The median normalized pairwise accuracy of the 11 published methods collapsed from 28.5% on unmatched pairs to 6.7% on matched pairs (Holm-adjusted *P* = 0.023) and stayed at 6.6% on within-region pairs (Fig. 2a–c; Supplementary Table 1).

Unlike whole-section correlation and matched pairs, within-region pairs scored a constant regional average near zero, and matching changed method rankings. Because matched spots could still lie in different regions, the class-mean baseline scored as high as the median published method on matched pairs (6.8% versus 6.7%; Fig. 2b). Within a single annotated region, however, the class-mean baseline assigned both spots the same value and scored 1.5%, lower than each of the 11 published methods (median 6.6%; Fig. 2c; Supplementary Note 1). Method rankings by whole-section correlation agreed closely with rankings on unmatched pairs (Kendall’s *τ_b_*= 0.818) but were nearly unrelated to rankings on within-region pairs (*τ_b_* = 0.091). The top-ranked method changed from EGNv2 to DeepSpaCE (Fig. 2g–i).

In Xenium samples of breast, pancreatic and lung cancer^27^, matched spot pairs retained substantial reproducible spot-to-spot differences, yet 11 spot-level models that we trained largely missed these differences. On matched pairs, the two halves of the measured RNA still agreed 32.9 percentage points above the majority-order baseline, compared with 45.5 percentage points before matching (Supplementary Fig. 1). By contrast, matching lowered the mean gain in pairwise accuracy of the 11 models over this baseline from 20.9 to 3.4 percentage points (*P* = 0.0039; Supplementary Fig. 1). Unlike the HER2ST rankings, the rankings of these 11 models changed little after matching (*τ_b_* = 0.927; Supplementary Table 2). These findings demonstrate that whole-section correlation overstates prediction quality by crediting methods for reproducing differences between tissue regions. However, directly evaluating local predictions against raw measured counts faces an obstacle, because at fine spatial scales molecular counts are sparse and dominated by Poisson sampling noise^13,14^. This measurement noise severely attenuates conventional correlations, making it difficult to determine whether poor local agreement reflects prediction errors or unavoidable assay noise. For spot pairs, the agreement between two RNA halves resolved this ambiguity, and a general metric should apply the same split-half reference to correlation at each spatial scale.

### RSC isolates spatial scales and separates prediction quality from sampling noise

To separate reproducible biological variation from sampling noise across spatial scales, we developed the reproducible-signal correlation (RSC) metric (Fig. 1a; Methods; Supplementary Note 2). In spatial transcriptomics, gene expression varies across multiple spatial scales, ranging from millimeter-scale tissue compartments to adjacent spots and individual cells. RSC isolates distinct spatial scales and evaluates predictions against the reproducible biological variation of the assay in three steps. First, each RNA molecule counted in a held-out test sample is randomly partitioned into two halves through binomial count splitting^19–21^. Under Poisson sampling, both halves share identical expected biological expression rates while carrying statistically independent sampling noise. Their mutual correlation, termed the split-half reliability, empirically quantifies the reproducible biological signal and establishes the theoretical noise ceiling on the agreement any noise-free prediction can reach^22^. Second, the predicted expression map and both RNA halves are decomposed into spatial scales, isolating variation across specific physical distances^28^. Third, at each spatial scale, RSC computes the Pearson correlation of the prediction with the RNA halves but replaces the RNA variance with the split-half covariance, which estimates the reproducible biological signal variance^23–25^. Computing this normalized correlation within individual tissue regions or cell classes isolates local biological variation from regional averages.

Before benchmarking prediction methods with RSC, we validated three essential properties of the metric through simulations. First, RSC specifically resolves individual spatial scales and is sensitive to exact spatial position. When we simulated transcript counts from synthetic expression patterns of known spatial scales in Visium HD sections of human colorectal cancer^2^ and evaluated the underlying patterns themselves as predictions, RSC peaked at the true scale of the simulated pattern in 262 of the 264 evaluable conditions (Extended Data Fig. 1a). Shifting the spatial phase of the predicted patterns by 90^◦^ abolished their spatial alignment with the simulated tissue profiles and reduced the mean absolute RSC to 0.0012 (Extended Data Fig. 1b).

Second, RSC provides a largely unbiased assessment across spatial scales, whereas conventional correlation with measured counts systematically penalizes fine-scale predictions. Because individual cells and small spots contain sparse transcript counts, random molecule capture introduces substantial sampling noise at fine scales. Conventional correlation compares predictions directly against noisy counts and therefore understates prediction quality whenever sampling noise is high. To test whether RSC corrects this attenuation, we simulated Poisson transcript counts using the measured counts as underlying expression rates in colorectal cancer sections, Xenium samples of breast, pancreatic and lung cancer^27^, and single breast cancer cells profiled by Xenium^27,29^. When the true underlying expression rates were evaluated as predictions, RSC remained between 0.956 and 1.000 from the broadest to the finest spatial scales (Extended Data Fig. 1c). By contrast, conventional correlation with the simulated counts dropped sharply at fine scales by an amount determined by the split-half reliability, which is the correlation between two random halves of the captured molecules at each scale^22^. Across all simulations, this understatement closely matched the theoretical relationship in equation (2) of Methods (Extended Data Fig. 1c). In the measured colorectal cancer data, the observed split-half reliability implies that even an ideal prediction matching the underlying expression would achieve a conventional correlation of 1.00 at the broadest scale but only 0.44 to 0.57 at the finest scale (Fig. 1c).

Third, RSC remains largely stable across changes in transcript capture depth, whereas conventional whole-section correlation mistakes sampling noise for prediction error. To increase sampling noise, we randomly downsampled captured RNA molecules in the Xenium samples of breast, pancreatic and lung cancer until one-eighth of the molecules remained. We then evaluated identical predictions from Ridge–Phikon-v2, a ridge regression model trained on features from the pathology foundation model Phikon-v2^30^, before and after downsampling. Reducing molecule counts lowered the fine-scale split-half reliability from 0.416 to 0.247 and caused conventional whole-section correlation to drop in all nine samples, from a mean of 0.509 to 0.408. By normalizing prediction agreement against the split-half reliability at each scale, fine-scale RSC instead changed from 0.337 to 0.386 (Extended Data Fig. 1d) and thereby separated prediction quality from sampling noise at each spatial scale.

### RSC reveals spot-level methods largely overlook fine-scale differences between spots

In real tissues, the expression of a gene changes both across millimeter-scale anatomical zones and between neighboring spots within the same zone. After validating RSC, we used it to test whether current spot-level methods resolve these local spot-to-spot expression differences or mainly reproduce broad regional patterns (Extended Data Table 1). With sampling noise corrected across scales, RSC revealed that spot-level predictions generally capture broad regional expression patterns but miss most fine-scale differences between nearby spots. In the Visium HD colorectal cancer sections, we evaluated ridge-regression predictions of 1,000 genes from Phikon image features^31^ in 32-*µ*m bins^2^. Mean RSC fell from 0.800 at the broadest scale of 4,096 *µ*m and above, which reflects major tissue compartments, to 0.298 at the finest scale below 128 *µ*m, which reflects expression differences among neighboring bins (Fig. 3a; Supplementary Table 3). Summing the 32-*µ*m bins into larger bins reduces sampling noise in raw counts but allows broad differences in tissue composition to dominate the evaluation (Fig. 3b). Across bin sizes from 32 to 512 *µ*m, the decline in RSC from broad to fine scales persisted in 22 of 24 combinations of patient and bin size (Fig. 3c; Extended Data Fig. 1e).

**Figure 3.**
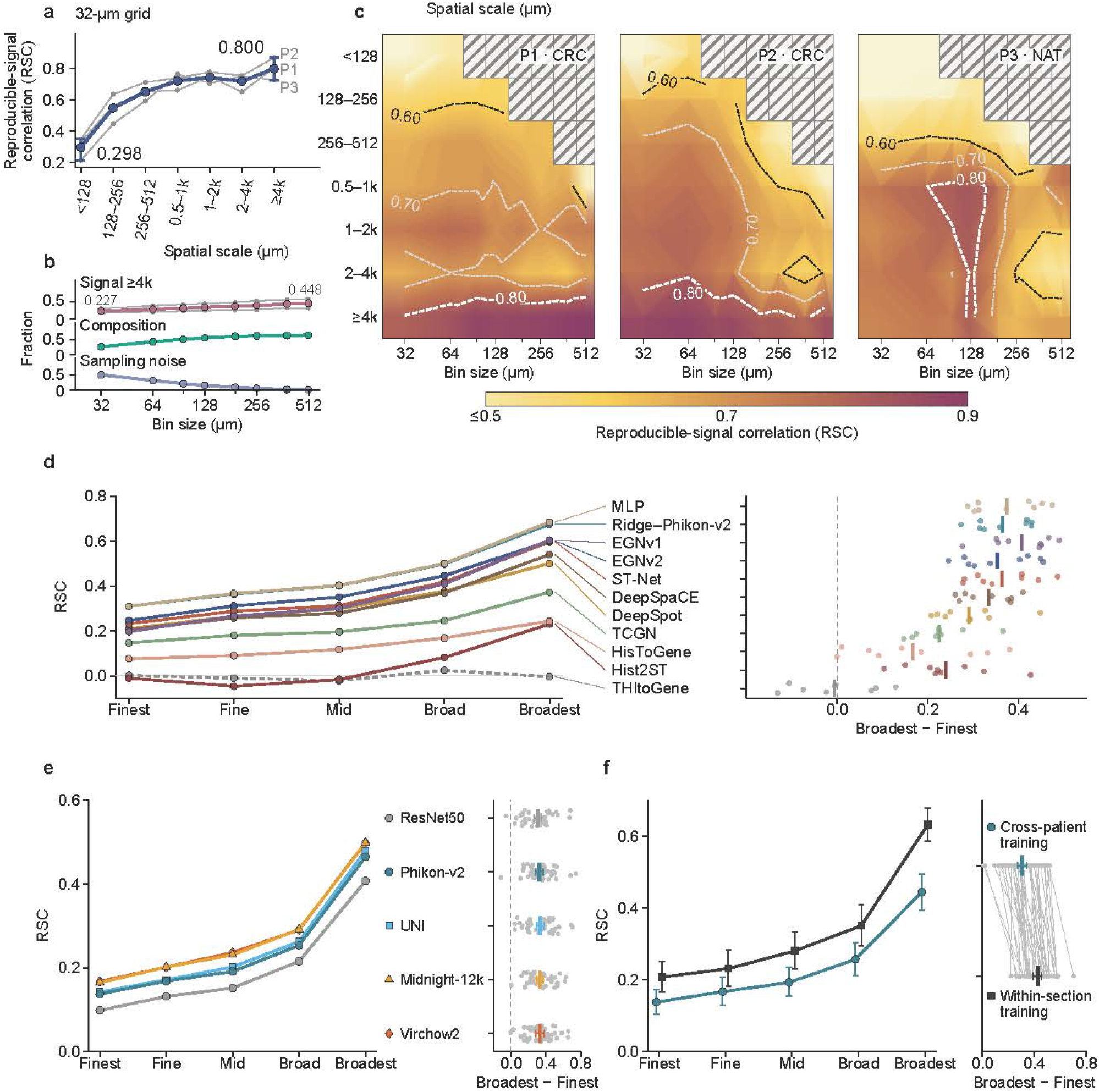
Reproducible-signal correlation (RSC) reveals spot-level methods mainly capture coarse tissue structure. a, RSC across spatial scales (*<* 128 *µ*m to ≥ 4k *µ*m; 1k denotes 1,024 *µ*m) for Ridge–Phikon predictions on a 32-*µ*m grid (*n* = 3 colorectal patients; P1 and P2 with colorectal cancer, CRC; P3 with normal adjacent tissue, NAT). Gray lines show individual patients and error bars show the patient range. **b,** Fraction of variance attributable to tissue composition (which inflates coarse-bin correlations) versus sampling noise (which attenuates fine-bin correlations) across bin sizes (32–512 *µ*m). The top line shows the share of reproducible signal at scales of at least 4,096 *µ*m. **c,** RSC heatmaps across spatial scales and bin sizes in P1–P3, demonstrating that RSC largely decouples spatial scale from bin size (hatching indicates unavailable combinations). **d,** Evaluation across 11 spot-level architectures in nine Xenium cancer samples (breast, pancreatic and lung). Most models achieve high RSC at the broadest scale but drop at the finest scale (Broadest – Finest difference shown right; THItoGene in gray). **e,** Comparison of pathology foundation models (Phikon-v2, UNI, Midnight-12k, Virchow2) against natural-image ResNet50 features (*n* = 45 HEST-Benchmark patients). **f,** Cross-patient versus within-section training in 45 HEST-Benchmark patients, showing that the fine-scale drop in RSC persists even when removing inter-patient generalization gaps. In **d**–**f**, bars show means, and error bars in **e**,**f** represent 95% confidence intervals. Broad-minus-fine tests are in Supplementary Table 3.

This loss of prediction quality at fine spatial scales largely held across the tested model architectures, patient cohorts and image encoders. In the Xenium samples of breast, pancreatic and lung cancer, 10 of the 11 spot-level models that we trained had lower RSC at the finest scale than at the broadest of five spatial scales in all nine samples (Fig. 3d; Supplementary Table 3). The 11 models ranged from linear ridge regression and a multilayer perceptron to published deep-learning architectures such as ST-Net, Hist2ST and EGNv2. Across 70 tissue sections from 45 patients in the nine prediction tasks of HEST-Benchmark^27^, Ridge–Phikon-v2 had lower RSC at the finest scale than at the broadest scale in 44 of 45 patients (sign-flip *P <* 10^−6^; Fig. 3e). Extracting image features from pathology foundation models raised mean RSC across all spatial scales compared with ResNet50 features trained on natural images (Fig. 3e; Extended Data Fig. 2c), yet the drop in RSC from the broadest to the finest scale remained between 0.31 and 0.34 for all five image encoders (Fig. 3e). Although measured spatial transcriptomics data contain substantial reproducible expression differences between nearby spots, the evaluated spot-level methods largely miss these fine-scale differences.

We next asked whether H&E morphology lacks local information beyond visible cell numbers or whether models trained across patients fail to extract it. In HEST-Benchmark, histological features beyond local cell counts accounted for 74% of the fine-scale RSC achieved by Ridge–Phikon-v2 (Extended Data Fig. 3a–c), a share that we measured by removing the prediction component explained by the numbers of segmented nuclei in each cell class from CellViT^32^ (Methods; Supplementary Table 4). Moreover, Ridge–Phikon-v2 models trained on other patients missed part of the fine-scale spot-to-spot differences that could be predicted from the same image features within a tissue section. When we trained Ridge–Phikon-v2 on three spatial blocks of a section to predict the held-out fourth block (Methods), fine-scale RSC rose from 0.152 to 0.219 alongside gains at broader scales (Fig. 3f; Extended Data Fig. 3d,e).

### Within-class RSC identifies single-cell methods that maintain fine-scale prediction quality

Beyond spot-level models, RSC computed within individual cell classes distinguishes single-cell methods that genuinely resolve part of the cell-to-cell expression differences from methods that barely resolve these differences. We trained three single-cell prediction methods, sCellST^29^, DeepSpot2Cell^33^ and GHIST^5^, and evaluated their predictions in breast cancer sections profiled by Xenium^27,29^. To separate expression differences among tumor cells from broad lineage differences between tumor, immune and stromal cells, we restricted the evaluation to malignant cells identified from H&E images by CellViT^32^. Across these malignant cells, RSC revealed a sharp divergence among the three methods (Fig. 4a). GHIST achieved the highest RSC at all five spatial scales (Fig. 4a) and the highest fine-scale RSC in each of the five patients under all four evaluation conditions (Extended Data Fig. 4). Whereas DeepSpot2Cell and sCellST dropped to near-zero RSCs of 0.042 and 0.020 at fine spatial scales, predicting almost none of the expression differences among neighboring malignant cells, GHIST maintained an RSC of 0.242, matching its performance at broader scales (Fig. 4a,b; Supplementary Table 3). RSC therefore demonstrates that routine H&E morphology carries part of the information that distinguishes expression states among individual malignant cells and identifies GHIST as the evaluated method that best recovers these fine-scale differences.

**Figure 4.**
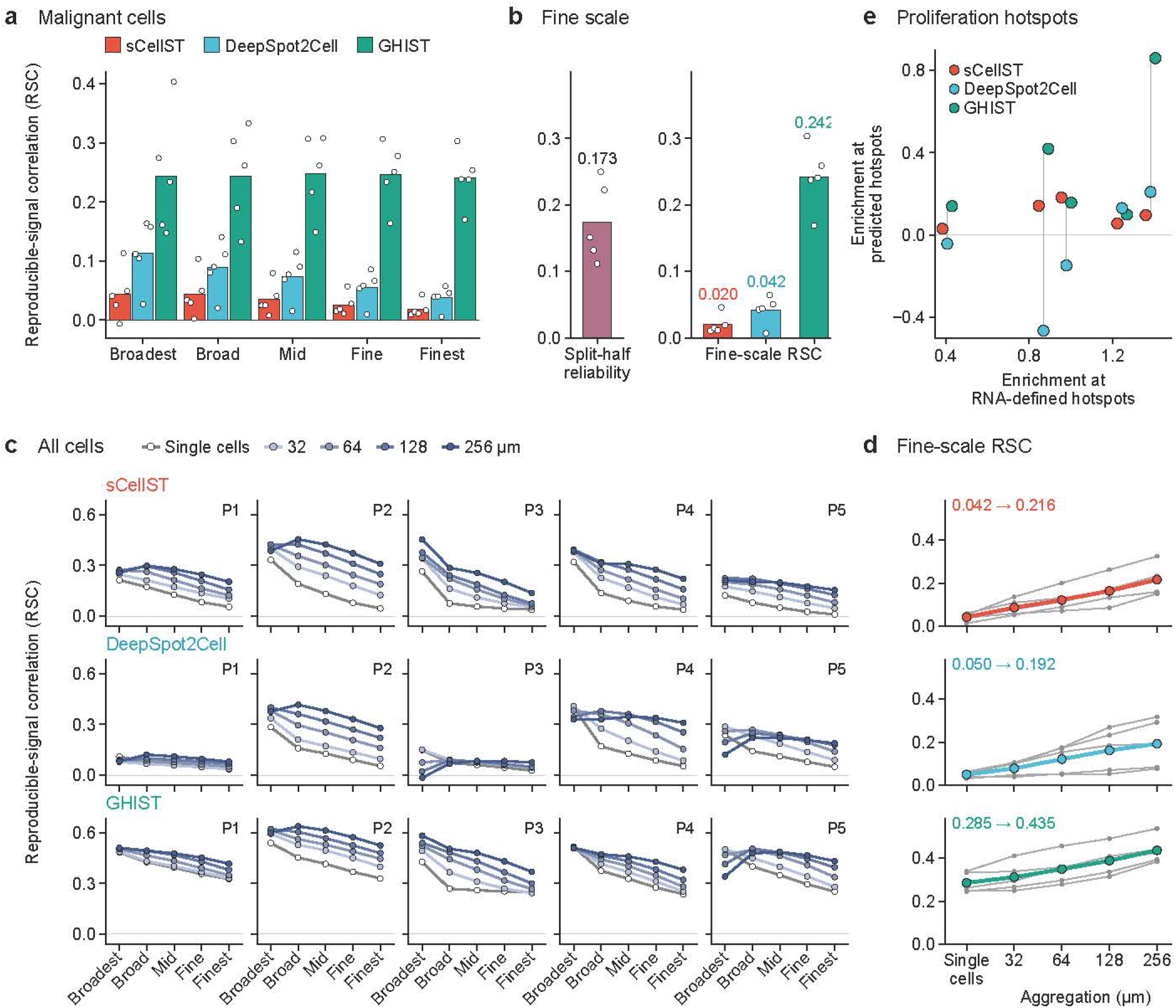
Reproducible-signal correlation (RSC) distinguishes the most robust of three single-cell prediction methods. a, Multiscale RSC evaluated strictly within CellViT-identified malignant cells for three single-cell methods (sCellST, DeepSpot2Cell, GHIST), controlling for broad cell-type composition differences between tumor, stroma and immune compartments. **b,** Fine-scale RSC compared with split-half reliability (0.173) among malignant cells, showing GHIST recovers part of the reproducible cell-to-cell variability (0.242), whereas sCellST (0.020) and DeepSpot2Cell (0.042) capture little fine-scale signal. **c,d,** Effect of spatial aggregation across all cells: multiscale RSC profiles (**c**) and fine-scale RSC (**d**) as single-cell predictions are averaged into bins (32–256 *µ*m). Aggregation inflates fine-scale scores (e.g., GHIST from 0.285 to 0.435) because local cell-type proportions confound the evaluation. **e,** Predicted versus RNA-defined proliferation hotspot enrichment among malignant cells. RNA-defined hotspots are identified in one split half and tested in the other; GHIST identifies proliferation hotspots best among the three methods (0.335), whereas sCellST and DeepSpot2Cell exhibit lower (0.101) or negative (–0.064) enrichment. Bars show means; points and gray lines show individual patients (*n* = 5 patients, nine sections; shared cells and genes). Broad-minus-fine tests are in Supplementary Table 3.

Without the within-class, scale-resolved evaluation of RSC, scores were inflated by cell-type composition. When evaluated across all cells without stratifying by cell class, scores mainly reflected average expression differences between cell lineages, allowing a naive class-mean baseline that assigns constant expression to malignant and non-malignant cells to outscore sCellST (fine-scale RSC of 0.081 versus 0.036 on the cells and genes of sCellST; Supplementary Note 1). Similarly, aggregating single-cell predictions into multicellular spatial bins allows local cell-type proportions to inflate evaluation scores^34^. For all three methods, averaging the exact same single-cell predictions into 256-*µ*m bins raised fine-scale RSC across all cells by 0.14 to 0.17 (Fig. 4c,d), masking the low RSC of sCellST and DeepSpot2Cell at single-cell resolution. Evaluating unbinned individual cells within a specific cell class via RSC is therefore important to separate genuine single-cell resolution from coarse cell-type averaging.

Crucially, the method identified by within-class RSC was also the best of the three at locating proliferative tumor hotspots in downstream biological analysis. We defined predicted hotspots as the 10% of 64-*µ*m malignant-cell neighborhoods with the highest predicted expression of a four-gene proliferation signature (Extended Data Fig. 5a) and quantified enrichment as the mean measured proliferation score in predicted hotspots minus that in the remaining malignant neighborhoods. In the measured RNA, defining hotspots in one split half and scoring them in the other half yielded a mean enrichment of 0.976 (Fig. 4e). Hotspots predicted by GHIST achieved the highest measured enrichment at 0.335, whereas sCellST and DeepSpot2Cell dropped to 0.101 and 0.064 (Fig. 4e; Extended Data Fig. 5b). The RSC of the proliferation score ranked the three methods in the same order as hotspot enrichment (Extended Data Fig. 5a). Computing RSC within a cell class thus identifies which of the three single-cell methods retain the fine-scale resolving power required for intratumoral discovery.

### RSC reveals models locate tertiary lymphoid structures yet poorly predict internal expression

Tertiary lymphoid structures (TLS) are intratumoral immune aggregates that have been linked to clinical response to immunother-apy^35^. We used RSC to test whether models that accurately locate TLS across a tissue section also resolve spot-to-spot expression differences inside these structures. Ridge–Phikon-v2 located expert-annotated TLS with high accuracy but achieved much lower RSC among spots within TLS than across distinct tissue regions (Fig. 5). We evaluated eight TLS-associated genes in kidney and lung cancer Visium sections^36,37^ whose TLS annotations were enriched for lymphocyte transcripts and spatially aligned with measured RNA (Methods; Supplementary Table 5). Each section was predicted by Ridge–Phikon-v2 trained on the remaining sections. Averaged over the eight genes, these predictions separated annotated TLS spots from immune-infiltrated, tumor and normal spots with an area under the receiver operating characteristic curve (AUROC) of 0.93 to 0.99 across sections, exceeding the AUROCs of 0.61 to 0.71 obtained from single-spot measured RNA. In lung cancer section LC3, predicted *LTB* expression followed the annotated TLS boundaries more tightly than measured *LTB* counts did (Fig. 5a–d).

**Figure 5.**
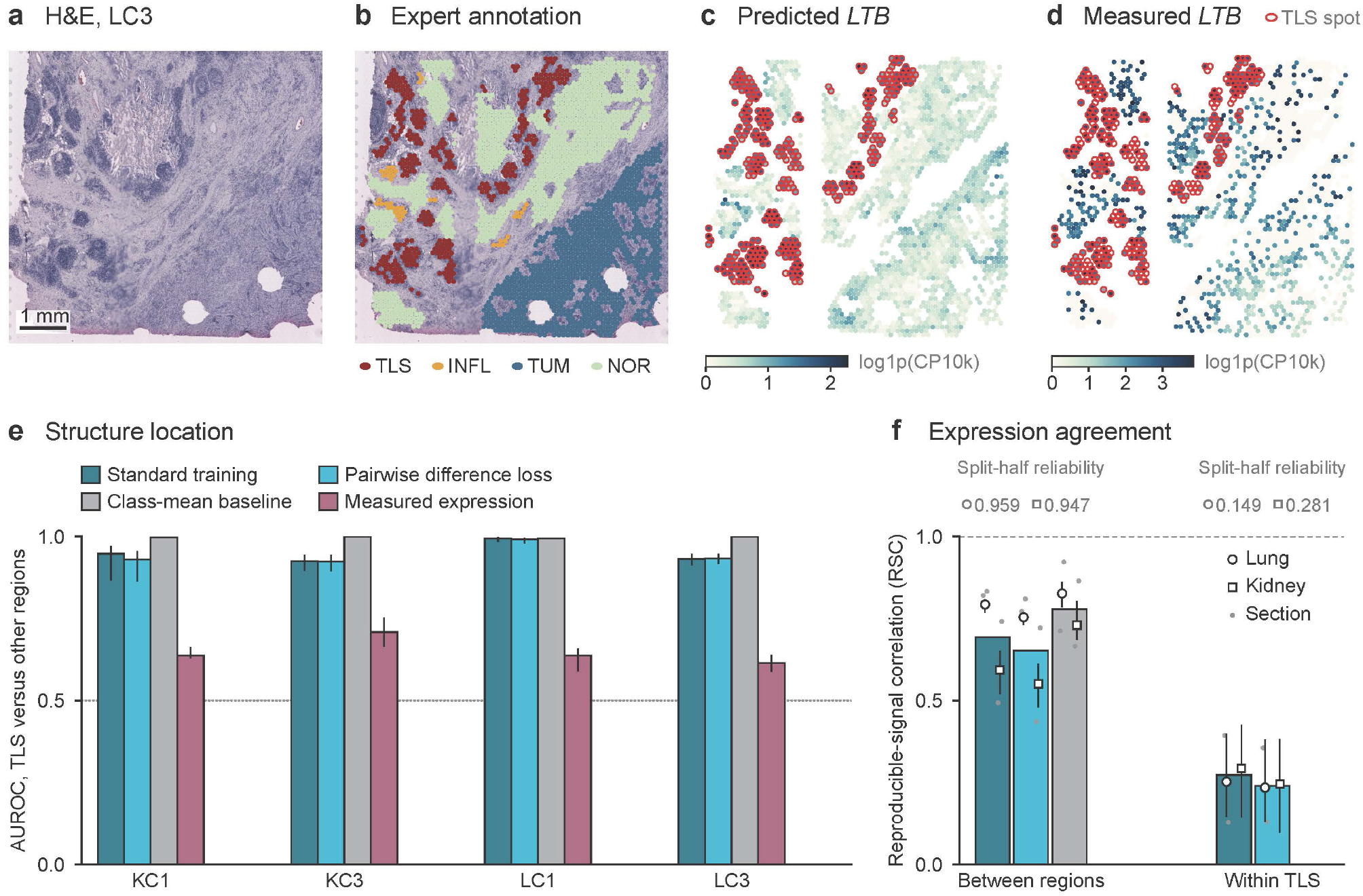
Reproducible-signal correlation (RSC) reveals models locate tertiary lymphoid structures (TLS) yet poorly predict internal expression. a–d, Spatial profiling of the TLS marker gene *LTB* in a representative Visium lung cancer section (LC3). Panels show H&E histology (a), expert histological annotations (b; TLS, immune-infiltrated, tumor and normal regions; red circles indicate 28 annotated TLS), Ridge–Phikon-v2 predicted *LTB* expression (c) and measured *LTB* counts (d; log1p(CP10k)). Scale bar, 1 mm. e, Conventional structure localization by AUROC can overstate prediction quality. Ridge–Phikon-v2 (standard training and pairwise difference loss) and a naive class-mean baseline all attain high AUROC (*>* 0.85, four cancer sections in kidney and lung averaged over eight TLS genes) for separating TLS from other tissue spots, demonstrating that coarse regional contrast suffices for high classification scores. Dotted line marks chance (0.5). f, RSC separates contrast between regions from expression differences within TLS. Between histological regions, models achieve high RSC (≈0.6–0.8, close to the class-mean baseline). Within annotated TLS, however, RSC shows that models largely fail to predict fine-scale, spot-to-spot expression differences beyond constant region labels. Predictions are from Ridge–Phikon-v2; dashed line in f marks RSC = 1. Error bars show 95% resampling intervals.

However, coarse region-level differences alone were sufficient to locate TLS. A naive class-mean baseline that assigns a constant expression value within each annotated region achieved AUROCs of 0.99 to 1.00 for separating TLS from other spots (Fig. 5e; Supplementary Note 1). Agreement across regions primarily reflected this same region-level contrast, as the class-mean baseline achieved between-region RSCs of 0.73 to 0.83 compared with 0.59 to 0.79 for Ridge–Phikon-v2 (Fig. 5f). When evaluated specifically among spots within annotated TLS, Ridge–Phikon-v2 achieved RSCs of only 0.25 to 0.29, far below its between-region performance (Fig. 5f; Supplementary Note 3). A high AUROC for locating anatomical structures can therefore overstate how well a method predicts internal gene expression, whereas computing RSC within TLS measures the spot-to-spot expression differences captured beyond region labels.

### An RSC-guided pairwise difference loss improves fine-scale predictions on average across models

Because RSC showed that resolving fine spatial scales requires capturing expression differences between nearby spots rather than only regional averages, we translated this principle into a training objective for spot-level models. Our single-cell evaluation with RSC demonstrated that H&E images contain information on expression differences within a cell class (fine-scale RSC of 0.242 for GHIST within malignant cells; Fig. 4b), yet conventional training losses for spot-level methods evaluate each spot independently without supervising how expression changes between neighboring spots. Building on relative-expression and pairwise learning strategies^38–40^, we formulated a pairwise difference loss guided by RSC that directly penalizes errors in the predicted expression difference between adjacent spots. We added this pairwise term to the original training objective of each architecture and compared the resulting models against standard training with the original objective alone (Methods).

Supervising expression differences between neighboring spots raised fine-scale RSC on average across models. Across the 11 spot-level models evaluated on the nine Xenium cancer samples, the pairwise difference loss increased mean fine-scale RSC from 0.194 to 0.205, with the multi-model mean rising in all nine samples (Fig. 6a; Supplementary Note 4). The loss also increased broad-scale RSC and conventional whole-section correlation by comparable amounts (Fig. 6b; Extended Data Fig. 6b,c). For Ridge–Phikon-v2 in HEST-Benchmark, adding the pairwise difference loss raised fine-scale RSC by 0.015, recovering 22% of the 0.067 gain obtained from within-section training (Extended Data Fig. 3f). Directly supervising local differences between adjacent spots therefore improves fine-scale predictions on average across diverse spot-level architectures.

**Figure 6.**
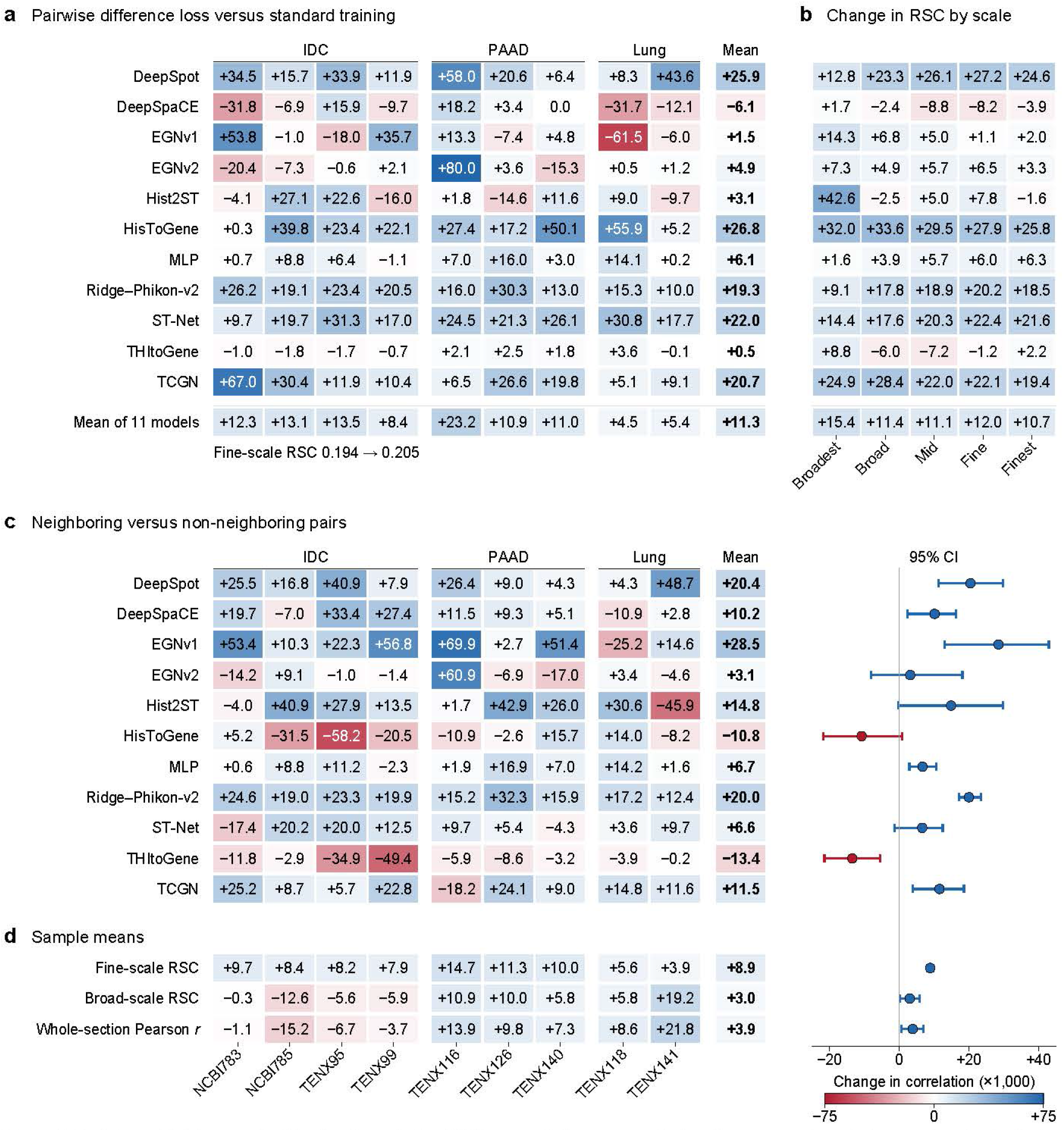
Training guided by reproducible-signal correlation (RSC) generally improves method performance. a, Impact of augmenting standard training with a pairwise difference loss that supervises expression gradients between adjacent spots. Heatmap shows changes in fine-scale RSC (×1,000) across 11 spot-level models in nine samples of invasive ductal carcinoma (IDC), pancreatic adenocarcinoma (PAAD) and lung cancer; mean fine-scale RSC across all models improves from 0.194 to 0.205 (+11.3). **b,** Mean change in RSC partitioned across spatial scales (Broadest to Finest), showing improvements at all five scales. **c,** Performance gain of neighboring pairs over degree-matched non-neighboring spot pairs, showing that fine-scale benefits arise mainly from spatial adjacency rather than generic pairwise regularization. **d,** Per-sample gains of neighboring over non-neighboring pairs, averaged across models, for fine-scale RSC (+8.9), broad-scale RSC (+3.0) and whole-section Pearson *r* (+3.9), demonstrating that local gradient supervision mainly benefits fine-scale RSC. Values and color scales indicate changes multiplied by 1,000; error bars show 95% bootstrap intervals across *n* = 9 patient samples.

Comparing adjacent spots contributed to this fine-scale improvement because standard point-wise objectives do not explicitly penalize models for smoothing over neighboring spots. Applying the pairwise difference loss to neighboring spot pairs increased fine-scale RSC by 0.009 more than applying the same loss to an equal number of randomly paired non-neighboring spots (exact sign-flip *P* = 0.0039; Fig. 6c; Methods), with a positive mean gain in 9 of the 11 models (Extended Data Fig. 6a). By contrast, broad-scale RSC and whole-section correlation differed much less between neighboring and non-neighboring supervision (Fig. 6d). At the same time, RSC showed that the effect of the pairwise difference loss depends on the downstream analysis and model type. In the Visium kidney and lung cancer sections, adding the loss lowered the RSC of Ridge–Phikon-v2 both between tissue regions and within annotated TLS in both cancer types (Fig. 5f; Supplementary Note 3). The loss also produced little change in fine-scale RSC for sCellST and DeepSpot2Cell, the two single-cell methods trained with spot-level expression (Extended Data Fig. 7; Supplementary Table 6). Evaluating predictions with RSC at each spatial scale and tissue compartment therefore both informs the design of fine-scale training objectives and reveals where training modifications improve or degrade predictions.

## Discussion

In this work, we developed and validated reproducible-signal correlation (RSC) and showed that correlation with the measured counts understates the quality of histology-based predictions at fine scales and overstates it for analyses within tissue regions or cell classes. Our scale-resolved evaluations demonstrate that measured RNA retains substantial reproducible signal that the evaluated methods largely miss at fine scales and within tissue regions or cell classes. Nevertheless, routine H&E images carry morphological features that reflect cell-to-cell variability in expression, because the evaluated single-cell method with the highest RSC predicted part of this variability within malignant cells. Predicting the signal that the evaluated methods miss would extend histology-based profiling from broad anatomical mapping to local cellular neighborhoods and individual cell states.

Methodologically, RSC evaluates prediction methods against the reproducible signal in the measured RNA and converges to the correlation with the measured counts as sampling noise approaches zero (Supplementary Note 2). RSC largely corrects both the understatement of correlation with the measured counts at fine scales and the overstatement for analyses within tissue regions or cell classes. The expected expression of simulated counts received an RSC near 1 across spatial scales, and a low RSC therefore means that a prediction misses reproducible signal. The class-mean baseline has no RSC within tissue regions or cell classes, and a positive RSC within a region or class means that a prediction captures signal beyond region or class labels. Evaluating prediction methods against reproducible signal reflects measurement principles shared with Fourier shell correlation in cryo-electron microscopy and Fourier ring correlation in optical nanoscopy^17,18^, metric calibration for molecular perturbation models^41^, and response evaluation in computational neuroscience^22,25,42–45^.

In practice, RSC can guide both the choice and the development of histology-based methods. When screening candidate methods on independent test sections with matched RNA, researchers can assess RSC at the spatial scales, tissue regions, cell classes and genes relevant to their clinical or biological questions. For analyses yielding low RSC, predictions warrant experimental validation or statistical frameworks that model predictive uncertainty^46,47^ and correct estimation bias using assayed subsets^48^. In method development, RSC locates the scales, tissue regions and cell classes where a method misses reproducible signal and measures whether a modification to training recovers this signal. The training loss that we designed to supervise how expression changes between neighboring spots improved fine-scale RSC, broad-scale RSC and whole-section correlation on average across 11 models. Yet the same loss lowered the RSC of Ridge–Phikon-v2 within tertiary lymphoid structures (TLS). A modification to training can therefore help one analysis and harm another, and each analysis needs its own evaluation with RSC. Because RSC is computed from the counts of a single assay and needs no replicate experiment, published methods can be reassessed on existing datasets. Beyond histology-based inference, RSC can also be applied to spatial gene expression imputation and super-resolution methods, which are likewise evaluated on measured counts that carry sampling noise^49^.

Several limitations of this study warrant future work. First, split-half reliability measures the reproducibility of a single assay at half its depth and reflects platform-dependent factors such as lateral molecular diffusion and probe capture limits^13,14,50,51^. As a result, RSC counts technical variation shared by the two halves as reproducible signal. Second, because count splitting assumes Poisson sampling, overdispersed counts can lower the RSC that the expected expression receives at fine scales (Extended Data Fig. 1f). Generalized count splitting for overdispersed counts would refine RSC at these scales. Third, the TLS analysis drew on four sections from two cancer types, and larger annotated clinical cohorts will show how far these findings generalize. Fourth, at single-cell resolution, method evaluation depends on the quality of nuclear segmentation and cell classification^32^. Each of the three single-cell methods also has its own supervision, network architecture and training data. Retraining these methods on common datasets will show which of these design choices raises RSC within cell classes. As histology-based predictions increasingly scale to single cells and large archival cohorts, evaluation with RSC at the scale, tissue region or cell class of each application will help determine which downstream analyses these predictions can support.

## Methods

### Reproducible-signal correlation

The reproducible-signal correlation (RSC) compares a prediction with two halves of the measured counts at each spatial scale. Count splitting, the split-half reliability and RSC are defined in the same way for all datasets. The scale operators, count transforms, gene weighting and numbers of count splits were set for each dataset (see ‘Scales and count transforms’).

#### Measurement model

Each count is one captured molecule. Given the expression of a gene in a bin, spot or cell, molecules are captured independently, and the count follows a Poisson distribution. The expected expression is the expectation of each half-count after the count transform of each dataset and contains no sampling noise. RSC estimates the correlation between a prediction and the expected expression.

#### Count splitting

We used binomial count splitting^21^. For the count *N* of each gene in each bin, spot or cell, we drew *N_A_* Binomial(*N,* 0.5) and set *N_B_* = *N–N_A_*, which gives two non-overlapping count matrices that sum to the original matrix. Given the expression, *N_A_* and *N_B_* are then independent Poisson counts with equal expectations^19,21^. The two halves therefore share the expected expression and carry independent sampling noise. *N*_half_ denotes the count in either half. Both halves were normalized and scored in the same way. Unless stated otherwise, directions and hotspots were defined on one half and evaluated on the other, following the principle of defining and evaluating signals on disjoint data^21^. In the HER2ST analysis of normalized pairwise accuracy, the roles of the halves were then exchanged, and the results were averaged over both assignments.

#### Scale decomposition

Each matrix was decomposed into spatial scales with linear operators that sum to the identity apart from the constant component, which centering removes^28^. The scale components of a centered matrix sum to the matrix. Each scale holds the spatial variation over one range of distances. ⟨, ⟩*_b_*denotes the inner product of the scale-*b* components of two matrices with the same genes and the same bins, spots or cells. Each inner product sums over all genes and all bins, spots or cells of a section or sample, and every ratio below is formed from these sums. The operators were built from a two-dimensional Fourier basis for the colorectal cancer bins, from the eigenvectors of a neighbor-graph Laplacian for the HEST spots and from diffusion operators on a cell graph for the breast cancer cells. These operators follow the layout of each dataset. A Fourier basis decomposes the regular grid of bins, the eigenvectors of a graph Laplacian extend this decomposition to spots on a neighbor graph, and diffusion operators separate scales on graphs of hundreds of thousands of cells without an eigendecomposition.

#### Split-half reliability

The split-half reliability of the measurement is the agreement between the two halves at a given spatial scale^22^. With *A* and *B* the two halves after the count transform of each dataset, *VA=*⟨*A,A*⟩*_b_*, *V_B_=*⟨*B,B*⟩*_b_*and *V_RNA_=*⟨*A,B*⟩*_b_*, the split-half reliability is 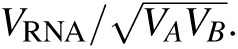 This ratio is the correlation between the two halves. Used as a prediction, the expected expression correlates with one half at about the square root of the split-half reliability^22^. For scores other than RSC, the score of the two halves is obtained by using one half as the prediction of the other half.

#### Reproducible-signal correlation

For a prediction *Ŷ* at spatial scale *b*, the mean covariance of the prediction with the two halves is *C* = (⟨*Ŷ,A*⟩*_b_* + ⟨*Ŷ,B*⟩*_b_*)*/*2, and the variance of the prediction is *V*_model_ = ⟨*Ŷ*,*Ŷ*⟩*_b_*. The RSC is a correlation corrected for attenuation, in which the split-half covariance *V*_RNA_ estimates the variance of the reproducible signal^23,25^:

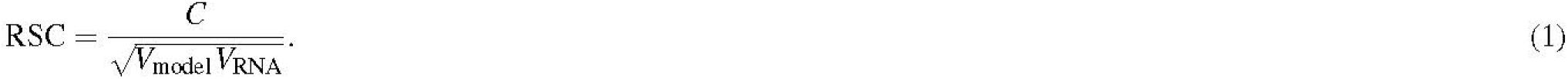

RSC was estimable when *V*_model_ and *V*_RNA_ were both positive. Negative values of RSC were kept.

#### Computation

RSC was computed in six steps. (1) The counts of the test section were split binomially into *N_A_* and *N_B_*. (2) The two halves were transformed and centered as described for each dataset, and predictions were centered and scaled by gene. (3) Predictions and both halves were projected onto the scales with the operators of each dataset. (4) At each scale, *C*, *V*_model_ and *V*_RNA_ were summed over all genes and all bins, spots or cells. (5) RSC was computed at each scale from these sums. (6) Steps 1 to 5 were repeated over independent count splits, and the scores were averaged.

#### Relation to the correlation with measured counts

For halves of equal variance, RSC equals the mean correlation of the prediction with the two halves divided by the square root of the split-half reliability *ρ_b_*. The correlation of the prediction with the mean of the two halves then satisfies (Supplementary Note 2)

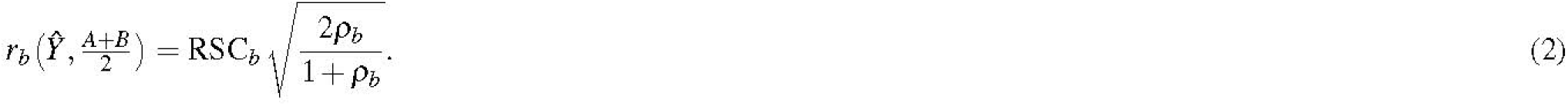

The same relation links the Fourier shell correlation between two cryo-electron microscopy half-maps to the correlation of the full map with the structure without noise^52^. As sampling noise approaches zero, *ρ_b_* approaches 1 and RSC approaches the correlation with the measured counts. Supplementary Note 2 also compares RSC with the correlation without correction for attenuation.

#### Repeated splits

Each analysis used 5 to 32 independent count splits. The number of splits affects the precision of the averaged scores and leaves their expected value unchanged.

#### Assumptions

The derivation assumes that each count is a captured molecule, that the counts are split before processing and that the prediction does not use the counts it is evaluated against. The counts analyzed here are molecule counts, as the HER2ST, Visium HD and Visium counts are deduplicated by unique molecular identifiers and each Xenium count is a decoded transcript. All splits were made before normalization, transformation and scale decomposition. The three colorectal models built partly from cell-type composition are an exception to the third assumption, as their inputs included cell-type proportions and sequencing depth estimated from the RNA of the evaluated section (see ‘Composition models’). Overdispersion of the counts can raise the split-half reliability and lower RSC (Supplementary Note 2). Technical variation shared by the two halves, such as lateral diffusion, is counted as reproducible signal.

### Datasets and evaluated methods

This study reanalyzed public human tissue datasets. Ethical approvals are reported by the original studies. Section, sample and patient identifiers are listed in Supplementary Table 7. All predictions were made for patients, samples, sections or blocks of a section held out from training, and the models were fitted on the training data only. Each prediction was scored on the genes and the spots or cells shared with the measured expression. Extended Data Table 1 lists the predictions used in each analysis.

#### HER2ST

We used eight sections from eight patients with HER2-positive breast cancer^8,26^. The analysis covered 3,481 spots and 785 genes shared by the predictions and the measured expression. Each section has pathologist annotations of tissue regions from the original study^26^. We evaluated the predictions of 11 published methods, DeepPT^53^, DeepSpaCE^54^, EGNv1^55^, EGNv2^56^, GeneCodeR^57^, HisToGene^58^, Hist2ST^59^, ST-Net^4^, TCGN^60^, THItoGene^61^ and iStar^49^. DeepPT and GeneCodeR were run as implemented in a published benchmark^8^. For each patient, we used the held-out predictions released by the benchmark^8^. The benchmark generated these predictions by four-fold cross-validation, in which each fold held out two patients and trained on five of the other six, with the sixth used for validation.

#### Colorectal cancer

We used five Visium HD sections from four patients with colorectal cancer^2^. Three patients formed the discovery set, and each was predicted by a model trained on the other two. The fourth patient was predicted by a model trained on all three discovery patients and served as an independent test of the spatial scales, bin sizes and scoring rules set on the discovery set. The two sections of this patient were scored separately, and the two scores were averaged. Predictions were generated by a ridge regression on Phikon image features^31^ (Ridge–Phikon). At 32 *µ*m, each bin was represented by the features of a 64-*µ*m patch centered on the bin and of a 256-*µ*m context patch, reduced to 64 whitened principal components. The analysis used the 1,000 protein-coding genes with the largest total counts in the discovery sections.

#### HEST-Benchmark

HEST-Benchmark^27^ comprises nine prediction tasks built from HEST-1k, a collection of public spatial transcriptomics samples with matched H&E images. The nine tasks contain 70 samples from 45 patients, and each task has its own 50-gene panel and patient folds. Patients were identified from the HEST metadata, and a sample without a patient identifier was treated as a separate patient.

The HEST samples are the nine Xenium samples of three tasks, four of breast invasive ductal carcinoma (IDC), three of pancreatic adenocarcinoma (PAAD) and two of lung cancer. Seven of these samples were released by 10x Genomics and two by Janesick et al.^62^ (Supplementary Table 7). These nine samples contain 48,273 spots, and each spot pools the transcripts within a 100-*µ*m square. These three tasks use a leave-one-sample-out split. We trained 11 spot-level models on the HEST samples. The models were a ridge regression on Phikon-v2 features^30^ (Ridge–Phikon-v2), a two-layer multilayer perceptron (MLP), ST-Net^4^, HisToGene^58^, Hist2ST^59^, THItoGene^61^, DeepSpot^36^, TCGN^60^, DeepSpaCE^54^, EGNv1^55^ and EGNv2^56^. Each model was trained with and without the pairwise difference loss (see ‘Training with the pairwise difference loss’).

For the analyses of the 45 HEST-Benchmark patients, we used all 70 samples with the patient folds provided by the benchmark. Ridge regression was fitted as for Ridge–Phikon-v2 to the features of five image encoders, ResNet50^63^, Phikon-v2^30^, UNI^64^, Midnight-12k^65^ and Virchow2^66^. The spots were scored as described for the HEST spots. The clear-cell renal cell carcinoma (ccRCC) task contains the samples INT1–INT24 from 24 patients, one sample per patient^35^. The ccRCC task served as a second cohort for the pairwise difference loss (Supplementary Fig. 2).

#### Breast cancer Xenium sections

We used nine Xenium sections from five patients with breast cancer^27,29^. Six of these sections were released by 10x Genomics and three by Janesick et al.^62^ (Supplementary Table 7). Transcripts with a quality value above 20 that were assigned to a cell were kept, and control probes were excluded. sCellST^29^ and DeepSpot2Cell^33^ were trained with spot-level expression from the separate section TENX39 and tested on all nine sections. GHIST^5^ was trained within the cohort, with one patient left out at a time (see ‘GHIST training’). TENX39 and the sections of patient P2 were obtained from the same supplier and have the same diagnosis. The cells and genes shared with the measured expression comprised 4,064,000 cells and 222 to 293 genes for sCellST and 3,341,649 cells and 261 genes for DeepSpot2Cell. Analyses of all cells used these shared cells of each method irrespective of class, except for the comparison of the three single-cell methods.

#### Kidney and lung cancer sections

We used formalin-fixed, paraffin-embedded Visium sections from University Hospital Zurich^37^, three of kidney cancer and five of lung cancer. The sections carry expert spot annotations of TLS, immune-infiltrated, tumor and normal regions made on the H&E images. The four sections KC1, KC3, LC1 and LC3 passed two RNA-based checks of the annotations and were analyzed (see ‘Evaluation of TLS-associated genes’). The sections KC2, LC2, LC4 and LC5 did not pass these checks. KC2 and LC4 had also been excluded in the original study because their spatial transcriptomics data did not align with the tissue morphology^36^. Predictions were generated by Ridge–Phikon-v2 on 112-*µ*m image patches, with features reduced to 256 principal components. Training details are given in Supplementary Methods. Each section was predicted by a model trained on all spots under tissue with non-zero counts in the other three sections, which included sections of the other cancer type. In a further condition, each model was trained only on the other section of the same cancer type. Values were computed separately for kidney and lung.

### Scales and count transforms

#### Colorectal cancer bins

The regular grid of each colorectal section was built from 8-*µ*m bins and aggregated to bin sizes of 32, 64, 96, 128, 192, 256, 384 and 512 *µ*m. A two-dimensional Fourier basis divided non-constant spatial variation into seven scales, which covered distances of less than 128, 128–256, 256–512, 512–1,024, 1,024–2,048, 2,048–4,096 and at least 4,096 *µ*m. The six boundaries double from 128 to 4,096 *µ*m, and the two open-ended scales assign every non-constant Fourier mode to a scale. Each analysis kept the measured tissue mask, with zeros outside the tissue. In the main analysis, for the fourth patient and in Extended Data Fig. 1a, every scale that contained Fourier modes of the grid was scored, including the partly represented scale below 128 *µ*m at the 32-*µ*m bin size. Each half-count was transformed as 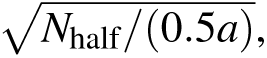 where *a* is the effective bin area, and centered across bins. The denominator corrects for bin area and for the halved depth of each half, and the square root compresses the range of the counts. For each gene *g*, the two transformed halves *A_g_* and *B_g_* were divided by the same factor, {(‖*A_g_*‖^2^ + ‖*B_g_*‖^2^)/2 ^1/2^. Predictions were centered and scaled to unit norm for each gene. Five count splits were used per patient.

#### HEST spots

Spots within 1.25 times the median nearest-neighbor distance were connected, and connected components of at least 20 spots were kept. The eigenvectors of the graph Laplacian were ordered by eigenvalue, the constant mode was removed, and the remaining modes were divided into five consecutive groups of near-equal size. From the smallest to the largest eigenvalues, these groups defined the Broadest, Broad, Mid, Fine and Finest scales. Wavelengths were estimated from the eigenvalues, except at the Finest scale, where the spacing of sign changes was used. Across the nine samples, the median wavelengths of the Broadest, Broad, Mid, Fine and Finest scales were 582, 333, 264, 212 and 168 *µ*m. Half-counts were transformed as log(1 + 2*N*_half_) and centered by gene. Twenty count splits were used. Each gene was weighted by the agreement between its two halves in the other samples of the same cancer type. For each held-out sample and gene *g*, d_g_^2^ and *c_g_* were the mean variance of a half-count and the mean covariance between the two halves in those other samples. The weight of gene *g* was w_g_=max(c_g_/d_g_^2^,0). Before projection onto the scales, both RNA halves of gene *g* were multiplied by 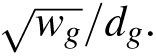 Each centered prediction was scaled to unit norm and multiplied by 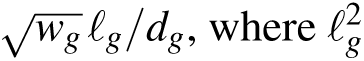 is the inner product of the two halves of gene *g* in the held-out sample, averaged over count splits.

#### Breast cancer cells

Each cell was connected to its eight nearest neighbors within 40 *µ*m. Edges had Gaussian weights with a width equal to the median edge length, and connected components of at least 20 cells were kept. With *K* the weighted adjacency matrix of this graph and *D_K_* its diagonal degree matrix, the lazy symmetric diffusion operator was *S* = (*I* + *D_K_*^−1^*^/^*^2^*KD_K_*^−1^*^/^*^2^)*/*2. The Broadest, Broad, Mid, Fine and Finest scales were *W*_1_ = *S*^16^, *W*_2_ = *S*^8^ – *S*^16^, *W*_3_ = *S*^4^ – *S*^8^, *W*_4_ = *S*^2^ – *S*^4^ and *W*_5_ = *I – S*^2^, respectively. These five operators sum to the identity. Half-counts were transformed as log(1 + 2*N*_half_) and centered by gene within connected components. As for the colorectal cancer bins, the two halves of each gene were divided by a common factor, and centered predictions were scaled to unit norm. Five count splits were used.

#### Fine-scale and broad-scale RSC

For spot-level models in HEST, the fine-scale RSC is the mean RSC at the Fine and Finest scales, and the broad-scale RSC is the mean RSC at the Broadest and Broad scales. In HEST, the fine-scale split-half reliability is the mean split-half reliability at the Fine and Finest scales. For the breast cancer cells, the fine-scale RSC is computed from *C*, *V*_model_ and *V*_RNA_ summed over the Fine and Finest scales. Scales were defined in micrometers for the colorectal cancer bins and relative to the neighbor graph of each HEST sample or breast cancer section. Scale labels are therefore not comparable across datasets.

### Evaluation between and within tissue regions

#### Spot pairs

In HER2ST, three sets of spot pairs were built within each of the eight sections, for the 785 genes and 11 methods. Unmatched pairs were drawn from all pairs of spots with predictions from all methods. Matched pairs had similar expression profiles and total counts. Within-region pairs were matched pairs in the same annotated region. Pairs were selected using only one RNA half, the source half. Scoring used only the other half, the evaluation half. Ten count splits were used, each with both assignments of the source and evaluation halves. For each gene, pairs were kept when their absolute source-half expression difference was at or above the 75th percentile. Matching parameters and gene eligibility are given in Supplementary Methods.

#### Normalized pairwise accuracy

For each pair, the predicted direction of a model was the sign of its predicted difference, and the measured direction was the sign of the count difference in the evaluation half. Pairs tied in the evaluation half were excluded, and a tied prediction was scored as incorrect. For the score of the two halves, the same pairs were assigned the direction observed in the source half. For each gene, the majority-order baseline predicted the direction observed most often in the five training patients of the fold. Pairwise accuracy was averaged within each section over distance quartiles, assignments of the halves and genes, and then over count splits. With Acc_model_, Acc_base_ and Acc_RNA_ the pairwise accuracies of the model, the majority-order baseline and the two halves, the normalized pairwise accuracy was *G* = (Acc_model_ – Acc_base_)*/*(Acc_RNA_ – Acc_base_). The normalized pairwise accuracy was calculated for each patient and averaged with equal patient weights. As the gain of a model over the baseline relative to the gain of the two halves, *G* can exceed 1 or fall below 0. Figures show *G* as a percentage. The denominator was positive in all 240 combinations of patient, pair set and count split (minimum 0.0796; Supplementary Fig. 3a). A sensitivity analysis with measured directions taken from depth-normalized counts is described in Supplementary Methods, and its results are given in Supplementary Note 5 and Supplementary Table 2.

#### Spearman correlation and all-pair concordance

For Fig. 2d–f, Spearman correlation and all-pair concordance with the evaluation half were computed across all annotated spots and within each annotated region, without matching, pair caps or difference thresholds. All-pair concordance is the fraction of spot pairs ordered correctly, with tied predictions given half credit. Inclusion criteria and averaging are given in Supplementary Methods.

#### HEST matched pairs

Unmatched and matched pairs were built and scored with the same procedure for the 11 spot-level models in the nine HEST samples. At most 1,000,000 unmatched pairs per sample were drawn uniformly before scoring. The majority-order baseline was fitted on the other samples of the same cancer type. Results are reported as the gain in pairwise accuracy over the baseline, in percentage points. The change from unmatched to matched pairs was computed over the combinations of gene, direction and distance quartile that could be estimated for both pair sets. For the summary, gains were averaged over the 11 models within each sample, then over the samples of each cancer type and then over the three cancer types with equal weights. The gain of the two halves over the baseline was computed on the same combinations and averaged over samples and cancer types in the same way. Model rankings by gain over the baseline were compared between unmatched and matched pairs with Kendall’s *τ_b_*.

#### Class-mean baseline

The class-mean baseline does not learn from images and predicts a constant expression within each region class or cell class. In HER2ST, the prediction for each gene at each spot of a held-out patient was the mean of log(1 + 10,000 *N/N*_total_) over spots of the same region class in the five training patients of the fold. Here *N* is the count of a gene and *N*_total_ the total count of the spot or cell. The two region classes were tumor and all other regions, and the tumor class comprised invasive cancer and cancer in situ. In the breast cancer Xenium sections, cells were divided into neoplastic and other cells by their CellViT^32^ class. Each cell was assigned the mean of log(1 + 100 *N/N*_total_) over cells of the same class in the sections of the other patients. In these sections, the baseline was evaluated on the cells and genes of sCellST with the five count splits of the main analysis, and sCellST was re-evaluated on the same cells and genes. In the kidney and lung cancer sections, each spot was assigned the mean expression of each TLS-associated gene over spots with the same expert annotation in the three training sections. All other steps followed the procedures used for the other predictions. The class-mean baseline gives tied predictions for every within-region pair. The normalized pairwise accuracy of this baseline is therefore reported with half credit for tied predictions and with tied predictions scored as incorrect, the rule used for the published methods. Supplementary Note 1 gives the results for this baseline, for variants with other class definitions and for a comparison after aggregating HER2ST spots into blocks.

#### Whole-section correlation

In HER2ST, whole-section Pearson *r* of each method and of the class-mean baseline was computed between its predictions and twice the evaluation-half counts, across the spots with predictions from all methods (Fig. 1b). Genes were included when they had at least 20 such spots, detection at 5% or more of spots, at least 20 molecules in the unsplit counts and matched pairs that could be scored. Values were averaged over genes, count splits, assignments of the halves and patients. In the other spot-level analyses, whole-section Pearson *r* was the gene-wise Pearson correlation between predictions and unsplit measured expression across the spots of the held-out section or sample, averaged over genes. For each HER2ST pair set, the ranking of the 11 methods by *G* was compared with the ranking by whole-section Pearson *r* with Kendall’s *τ_b_*.

### Tests of the reproducible-signal correlation

#### Known signals

On the colorectal grid, synthetic signals of known scale were multiplied by the measured tissue mask of each discovery section. Poisson or negative binomial counts were generated from these signals and then split and scored in the same way as the measured counts. The tests covered the three discovery patients, eight bin sizes, seven scales, 64 genes and 12 repeats per condition. Negative binomial dispersions were estimated by the method of moments from the variance of the measured counts across bins for each patient and bin size and truncated to the range 10^−4^ to 2. These estimates count all variation between bins beyond Poisson sampling, including biological variation, as overdispersion. The target correlation of the predictions was 0, 0.3, 0.6 or 1 (Extended Data Fig. 1f), and the spatial phase of the predicted patterns was shifted by 0^◦^, 22.5^◦^, 45^◦^, 67.5^◦^ or 90^◦^. A condition was assigned correctly when the injected scale had the largest RSC. At a target correlation of 1, the estimable conditions assigned correctly were counted (Extended Data Fig. 1a), and the mean absolute RSC was computed after the 90^◦^ phase shift (Extended Data Fig. 1b).

#### Expected expression as a prediction

The measured counts were treated as Poisson rates, and 20 count matrices were drawn for each test. On each drawn matrix, we used the expected expression as a prediction, defined as the expectation of the transformed half-count used by each score. The RSC of the expected expression of genes ranged from 0.956 to 0.998 in the breast cancer cells, 0.991 to 1.000 in the HEST spots and 0.994 to 1.000 in the colorectal cancer bins at the scales whose shortest distance was at least twice the bin size. The colorectal analysis at the 32-*µ*m bin size also scored the partly represented scale below 128 *µ*m, where the RSC of the expected expression was 0.9965 to 0.9973. For the biological analyses, the RSC of the expected expression ranged from 0.999 to 1.003 for the proliferation score of malignant cells and from 0.981 to 1.002 for the TLS-associated genes in the terms that could be tested (see ‘Evaluation of TLS-associated genes’).

#### Expected expression and the mean of the halves

The same procedure was repeated with 20 drawn matrices for the three colorectal discovery patients at the 32-*µ*m bin size, the nine HEST samples and the malignant cells and 218 genes of the comparison of single-cell methods. At each scale, we computed the RSC of the expected expression, its correlation with the mean of the two transformed halves and its correlation with the transformed unsplit counts (Extended Data Fig. 1c). The correlation with the mean of the halves divided by 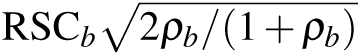 was 1.000 at every scale, and the correlation with the unsplit counts deviated from the correlation with the mean of the halves by at most 0.014. With the measured counts as rates, the drawn counts have a higher split-half reliability than the measured counts. The correlation of the expected expression with the mean of the halves of the measured counts was therefore computed from equation (2) with the split-half reliability of the measured counts, for each colorectal patient and HEST sample and for the malignant cells (Fig. 1c; Extended Data Fig. 1c).

#### Downsampling

The measured counts of the nine HEST samples were downsampled binomially to fractions of 0.5, 0.25 and 0.125 of their depth, with five downsampled matrices per fraction. Each downsampled matrix was split 20 times and scored as the measured counts, with the same gene weights and the predictions of Ridge–Phikon-v2 after standard training (Extended Data Fig. 1d). The factor *ℓ_g_*was recomputed in each downsampled matrix, and in a sensitivity analysis it was fixed at its value at full depth. Whole-section Pearson *r* was the correlation of each gene between the predictions and log(1 + *X*) of the downsampled counts *X* over all spots, averaged over the 50 genes. Scores were averaged over the five downsampled matrices within each sample and then over the nine samples. With *ℓ_g_*fixed at its value at full depth, fine-scale RSC changed from 0.337 to 0.358 at one-eighth of the depth and rose in seven of nine samples.

#### Semi-synthetic test in HEST

For each cancer type, 200 replicates were generated from a beta-binomial count model that used the observed neighbor graphs, total counts and gene marginals. Predictions combined the transformed latent rates with an orthogonal component of equal variance at target correlations of 0, 0.3, 0.6 and 1. A rotation series used angles of 0, *π/*8, *π/*4, 3*π/*8 and *π/*2. At a target correlation of 1, RSC was below 1 at every scale, and the largest deviation of the scale-specific RSC from 1 was 0.056 for IDC, 0.024 for PAAD and 0.045 for lung cancer. In all three cancer types, RSC passed the checks of the response to a target correlation of 0, the increase with the target correlation, the range between scales and the response to rotation. The largest range between scales at target correlations of 0.3 or 0.6 was 0.002.

#### Further tests

A resampling test of the measured HEST counts and a test of scoring on cell graphs are described in Supplementary Methods. In the resampling test, the scores of 324 conditions deviated from their values without sampling noise by at most 0.0109.

### Evaluation across spatial scales

#### Broadest-to-finest decline

For the HEST spots and the breast cancer cells, the broadest-to-finest decline was the RSC at the Broadest scale minus that at the Finest scale. For the colorectal cancer bins, the decline was the RSC at the scale of at least 4,096 *µ*m minus that at the finest scored scale. At the 32-*µ*m bin size, the finest scored scale was the scale below 128 *µ*m. In a sensitivity analysis, each RNA half was divided by its total count in each bin or spot before transformation, and the decline was recomputed for the colorectal discovery patients and the nine HEST samples. With this normalization, the mean decline of Ridge–Phikon in the colorectal discovery patients at the 32-*µ*m bin size changed from 0.502 to 0.468, and the decline of Ridge–Phikon-v2 remained positive in all nine HEST samples (mean, 0.297).

#### Aggregation in colorectal cancer

Bin size was changed while the tissue, genes and underlying 8-*µ*m bins were held fixed. The share of reproducible signal at scale *b* was *V*_RNA_*_,b_/* ∑*_b_*^′^ *V*_RNA_*_,b_*_′_. For Fig. 3c, the decline was computed for each discovery patient at each of the eight bin sizes. For the fourth patient, the decline was averaged over the eight bin sizes (Extended Data Fig. 2b).

#### Composition models

The composition model used cell-type proportions, sequencing depth and tissue fraction. Sequencing depth was log(1 + *N*_total_*/a*), where *N*_total_ is the total UMI count and *a* the effective area of a bin. Cell-type proportions were estimated from the RNA of the same section with 450 marker genes and cell-type profiles from the other two discovery patients. For Extended Data Fig. 2a, the morphology-based predictor was a ridge model that added Phikon image features to these covariates, and the image-only predictor was Ridge–Phikon. The local-context network combined the estimated cell-type composition with Phikon features of a local and a context patch. The fixed scale-specific predictor, labeled ‘Fixed ensemble’, used the local-context network at 32 and 64 *µ*m, the morphology-based predictor at 96, 128, 192 and 256 *µ*m and the composition model at 384 and 512 *µ*m. The composition model, the morphology-based predictor and the Fixed ensemble were the three models built partly from cell-type composition estimated from the RNA of the evaluated section.

#### Decomposition of the mean squared error

We decomposed the error *R*_0_ of a training-mean predictor, which assigns each gene its mean in the training patients, as

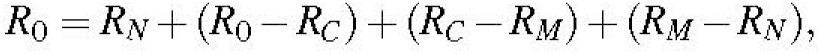

where *R_N_* is the error due to sampling noise, *R_C_* the error of the composition model and *R_M_* the error of the evaluated model. Errors were held-out mean squared errors of the square-root-transformed half-counts, without centering. For gene *g*, the sampling-noise term was 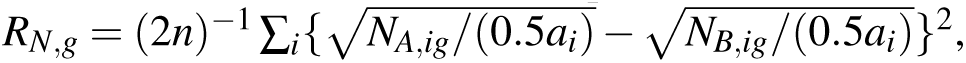 where *N_A_* and *N_B_* are the two half-counts, *a_i_* is the effective area of bin *i* and *n* is the number of bins. The sampling-noise term was averaged over count splits. Within each patient and bin size, gene-wise errors were summed before division by *R*_0_. The four terms then give the fractions of error due to sampling noise, explained by composition, removed by the model and unattributed. Patients were weighted equally, and the decomposition used 17,977 protein-coding genes.

#### Spot-level models in HEST

For Fig. 3d, the standard-training predictions of all 11 HEST models of the training comparisons were scored in the same way as Ridge–Phikon-v2. The broadest-to-finest decline was computed for each model and sample. For Extended Data Fig. 2c, ridge models on Phikon-v2 and on ResNet50 features^63^ were fitted in HEST with the procedure of Ridge–Phikon-v2.

#### Cell-count model

The cell-count model predicted each gene at each spot from the numbers of nuclei in the sequenced area of the spot for the five CellViT classes^32^, neoplastic, inflammatory, connective, dead and epithelial. For each gene, an intercept and non-negative class coefficients were fitted by non-negative least squares on all training spots of each fold, and the prediction was log(1 + *N̂*), where *N̂* is the fitted count. The nuclear-density model used only the total number of nuclei. CellViT detected almost no nuclei in the ccRCC samples INT20, INT22 and INT23, and these samples were kept. To measure the part of the RSC of Ridge–Phikon-v2 outside the prediction of the cell-count model, the ridge prediction *p* at each scale was projected onto the cell-count prediction *c* in each test sample, *β*=⟨*p, c*⟩*/*⟨*c, c*⟩. The RSC of *p* was then divided into the parts carried by *β_c_* and by *p–β_c_*, both normalized by the variance of *p*. With *β* fitted in the test sample, the part outside the cell-count prediction is a conservative estimate. The combined model and the scoring on matched pairs are described in Supplementary Methods.

#### Within-section training

Each HEST-Benchmark section was divided into four blocks at the medians of its spot coordinates. Each block was predicted by Ridge–Phikon-v2 trained on the spots of the other three blocks that lay more than 224 *µ*m (two image-patch widths) from every spot of the block. The colorectal sections used the model of Fig. 3a and a distance of 512 *µ*m, two context-patch widths. Predictions from training on other patients were scored on the same blocks. Each block was scored with its own graph and scales, using the count splits of the whole section and the same gene weights, and block scores were averaged within each section. Five further settings are described in Supplementary Methods.

### Evaluation within cell classes

#### GHIST training

We ran GHIST from its public implementation (commit 917456be305f) with the full framework and no pretrained expression weights. The framework included three components, all ten loss terms, the neighborhood and cell-type branches and the cell-type expression profiles. We trained GHIST in a leave-one-patient-out design over the five patients. Each fold excluded all sections of one patient from training, validation, normalization and model selection. In each fold, GHIST was trained with five seeds, for 25 fits in total. Training targets were the cell-level Xenium counts of the 218 genes shared with the other methods. Nucleus matching and cell-type labels are described in Supplementary Methods. In each training section, the last 20% of image rows formed the validation region. Each fit used the first ten epochs of the published 50-epoch schedule. Among these epochs, we chose the checkpoint with the smallest sum of two ranks on the validation cells of the fold, one by mean gene-wise Pearson correlation and one by macro-F1.

#### Malignant cells

Malignant cells were the cells that CellViT^32^ assigned to the neoplastic class on the H&E image. Labels were transferred to Xenium cells by nearest-centroid matching within 5 *µ*m.

#### Comparison of single-cell methods

The main cell-level outcome was the fine-scale RSC in malignant cells. Comparisons of GHIST, sCellST and DeepSpot2Cell in Fig. 4 and Extended Data Figs. 4 and 5 used the cells and the 218 genes shared by all three methods. The shared cells comprised 2,524,782 of 4,070,765 cells (49–75% per section), including 873,249 malignant cells (83.4% of all malignant cells). A gene was excluded for a method when its predictions did not vary within connected components, which removed 7–17 genes for GHIST and none for the other two methods. Scores therefore used the 201–211 genes eligible for all three methods in each section. For the fine-scale RSC, *C*, *V*_model_ and *V*_RNA_ were averaged over the sections of each patient before the ratio was formed. The RSC of the proliferation score was calculated in each section and then averaged within patients. For GHIST, each seed was evaluated separately, and the five values were averaged within each patient. The three further conditions of Extended Data Fig. 4 are described in Supplementary Methods.

#### Aggregation in breast cancer cells

Predictions and measured expression were held fixed while the same cells were grouped into 32-, 64-, 128-and 256-*µ*m bins. Within each bin, half-counts were summed and divided by the number of cells, and predictions were averaged over the same cells. For bin size *s*, graph nodes were the occupied bins, placed at the mean coordinates of their cells, and edges were kept up to a length of 1.5*s*. Count splitting and scoring were repeated at every bin size.

#### Proliferation hotspots

The signature comprised the four cell-cycle genes *MKI67*, *TOP2A*, *CENPF* and *PCLAF* of the Xenium panel and was evaluated in malignant cells. Each malignant cell defined a neighborhood of radius 64 *µ*m. Neighborhoods with fewer than five malignant cells were excluded. The score of a cell was log(1 + 1,000 *N*_signature_*/* max(*N*_total_, 1)), where *N*_signature_ is the summed count of the four genes and *N*_total_ the count of the cell over all measured genes. For model scores, predictions were clipped at zero and transformed with exp(*x*)–1 before summation, and the denominator summed the genes shared by the prediction and the measured panel. For Extended Data Fig. 5b, the measured score was also computed with *N*_total_ summed over the 218 shared genes. The score of a neighborhood was the mean over its cells. The counts were split 20 times. The 10% of neighborhoods with the highest model score defined predicted hotspots. Hotspot enrichment was the mean score of the second RNA half within these hotspots minus the mean score in the remaining neighborhoods. Hotspots defined from the first RNA half gave the enrichment of the two halves. The RSC between predicted and measured neighborhood scores was computed without spatial decomposition.

### Evaluation of TLS-associated genes

#### Genes and spots

The evaluation used eight TLS-associated genes that the original study^36^ analyzed in the kidney and lung cancer sections. The eight genes were *LTB*, *CXCL13*, *MS4A1*, *CCL19*, *CORO1A*, *CD19*, *SPIB* and *BLK*. Spots annotated as TLS, immune-infiltrated, tumor or normal were evaluated, and spots with unassigned, missing or lymph node labels were excluded. Annotated TLS spots adjacent on the Visium grid formed one TLS structure. Measured expression was log(1 + 10,000 *N/* ∑*_g_ N*), with the sum over all genes. Each RNA half was normalized by its own total, and the counts of each section were split 20 times. *RNA-based checks.* Two RNA-based checks were applied to all eight sections. First, a section passed when a lymphocyte score separated annotated TLS from tumor and normal spots with an area under the receiver operating characteristic curve (AUROC) whose 95% interval lay above 0.5. The lymphocyte score was the mean of log(1 + 10,000 *N/* ∑*_g_ N*) over *PTPRC*, *CD52*, *CD3E*, *CD2* and *CD79A*, five genes outside the eight TLS-associated genes. Intervals were obtained from 2,000 resamples of TLS structures with the other spots fixed. LC2, LC4 and LC5 failed this check. Second, the variance of the first 20 principal components of expression explained by the four annotation classes was compared between the recorded annotation and 21 shifted, rotated or mirrored versions of the annotation (Supplementary Methods). A section passed when no transformation raised the explained variance by more than 0.01. Seven sections passed this check. In KC2, the annotation explained 0.059 of the variance as recorded and 0.474 after rotation by 180^◦^ (Supplementary Table 5).

#### Decomposition by region

Within each section, each prediction and each RNA half was decomposed into the mean of its annotated region and the residual from that mean, and the two parts summed to the expression. Whole-section agreement used the expression, agreement between regions the region means, agreement within regions the residuals and agreement within TLS the residuals of TLS spots. The within-TLS term was further decomposed into a part between TLS structures and a part within them. Each gene was divided by the standard deviation of its unsplit expression over the spots of a term. The quantities *C*, *V*_model_ and *V*_RNA_ were summed over the eight genes before the ratio was formed, and the value for each cancer type was computed from these sums averaged over sections. For TLS location, the AUROC of each gene for annotated TLS versus immune-infiltrated, tumor and normal spots was computed in each section and averaged over the eight genes. The analysis was repeated with LC2 included (Supplementary Note 3).

### Training with the pairwise difference loss

#### Pairwise difference loss

For a model with training targets *Y*, predictions *Ŷ*, residual *E* = *Y–Ŷ* and *D* the incidence operator of a graph over training spots,

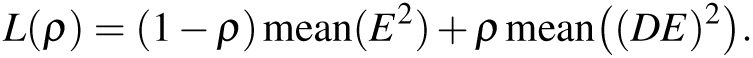

Standard training used *ρ* = 0, and training with the pairwise difference loss used *ρ* = 0.5. For an edge joining spots *i* and *j*, the second term penalizes the error in the predicted change in expression between the two spots^38^, (*Y_i_ – Y_j_*) (*Ŷ_i –_Ŷ_j_*). The graph was built within each training sample by connecting spots no further apart than 1.5 times the median nearest-neighbor distance, which joins directly adjacent spots without bridging gaps in the tissue. Each model kept its own auxiliary losses and their weights.

For ridge models, with *E* = *Y* − *Xβ*, *n* training spots and *m* edges, the objective was

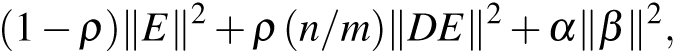

where the factor *n/m* places the difference term on the scale of the term for *n* spots. Features were reduced to 256 principal components fitted on the training samples^67^. In HEST, *α* = 100*/*(256×50) = 0.0078125. The training graph was fixed before optimization, and the edges drawn by each model are given in Supplementary Methods.

#### HEST models

Each of the 11 HEST models entered two training comparisons, the pairwise difference loss compared with standard training and neighboring compared with non-neighboring pairs. The two conditions of each comparison used the same architecture, encoder, inputs, data splits, initialization, optimizer and checkpoint rule. Stochastic models were trained with three seeds, and predictions were averaged over seeds before scoring. For TCGN in sample NCBI783, both conditions of each training comparison used two of the three seeds. TCGN training stopped when non-finite values occurred, and the checkpoint with the lowest validation error before the stop was used. Such stops occurred in 10 of 26 fits with standard training and 11 of 26 with the pairwise difference loss. Results without TCGN are given in Supplementary Note 4. The main outcome was the mean fine-scale RSC across the nine HEST samples. Secondary outcomes were the broad-scale RSC and whole-section Pearson *r*. Pairwise accuracy between neighboring spots (Extended Data Fig. 6d–g) is defined in Supplementary Methods.

#### Comparison with within-section training

For Ridge–Phikon-v2 in the 45 HEST-Benchmark patients, the pairwise difference loss was added to training on other patients, and its fine-scale gain was scored on the blocks of the within-section analysis. This gain was divided by the fine-scale gain from within-section training.

#### Non-neighboring pairs

Each training graph was rewired by double-edge swaps, which keep the number of edges and the degree of every spot and connect only spots further apart than the neighbor radius. All 11 models were trained on these non-neighboring pairs with the same loss weight. The input of Hist2ST depends on the spots in each batch of edges. The comparison model for Hist2ST was therefore a second neighboring-pair model whose batches contained the same spots as the non-neighboring batches. The two Hist2ST conditions used the same inputs and changed only the edges that entered the loss. Fourteen of the 26 TCGN fits on non-neighboring pairs stopped in the same way.

#### Further cohorts

In the ccRCC cohort, an MLP and a ridge model for the proliferation score were trained with and without the pairwise difference loss on 16 training patients of each fold (Supplementary Methods). For the kidney and lung cancer sections, the pairwise difference loss at *ρ* = 0.5 was added to the training of each Ridge–Phikon-v2 model.

#### Single-cell methods

The same loss was added to sCellST and DeepSpot2Cell, where the loss compared predicted with measured changes in expression between training spots of TENX39. The conditions and outcomes are described in Supplementary Methods (Extended Data Fig. 7; Supplementary Table 6). A comparison that replaced the ResNet50 image encoder of sCellST with Phikon-v2 is described in Supplementary Note 4.

### Statistical analysis

Apart from the TLS analysis and the tests of RSC, intervals and tests treated patients, donors or samples as the independent biological units^68^. Spots, cells, genes, spatial scales, count splits, training seeds and sections of the same patient were repeated measurements within these units. Unless stated otherwise, scores were averaged over count splits and then over the sections of each patient, and patients received equal weight. Bootstrap 95% percentile intervals are reported as 95% confidence intervals (CIs). Holm adjustment was applied within each family of contrasts stated below, and the resulting *P* values are labeled Holm-adjusted. The tests on cell graphs used Bonferroni adjustment, and all other *P* values are unadjusted. Significance required a two-sided *P <* 0.05 after any adjustment.

#### Normalized pairwise accuracy

In HER2ST, each summary is the median across the 11 methods of the patient means of each method, and this median was recomputed in every bootstrap draw and permutation. Intervals are 95% percentile intervals from 20,000 bootstrap draws of patients, and each drawn patient kept all 11 methods and three pair sets. Contrasts between pair sets were tested with two-sided exact paired permutation tests over all 2^8^= 256 patient-level label swaps, with Holm adjustment across the three contrasts. For method rankings, the same bootstrap draws resampled whole-section Pearson *r* and the three normalized pairwise accuracies together. Ranks and Kendall’s *τ_b_* were recomputed in each draw, which gave intervals for *τ_b_* and for its changes between pair sets.

#### HEST matched pairs

The paired change in gain over the baseline was averaged over the 11 models within each sample, then over the samples of each cancer type and then over the three cancer types with equal weights. The 95% CI of this mean came from 10,000 bootstrap draws of samples stratified by cancer type, and the exact sign-flip test used all 2^9^ = 512 sign patterns. The same interval and test were computed for each model, with Holm adjustment across the 11 models (Supplementary Fig. 1b).

#### Spatial scales

Intervals for the mean at each scale and for the broadest-to-finest declines came from 20,000 bootstrap draws of the nine HEST samples, stratified by cancer type, and of the five breast cancer patients. For the three colorectal patients, intervals are the range of patient values. These declines were tested with exact two-sided sign-flip tests over all sign patterns of the units, 8 for the colorectal patients, 512 for the HEST samples and 32 for the breast cancer patients. Holm adjustment was applied across the four declines for colorectal cancer, HEST Ridge–Phikon-v2, sCellST and DeepSpot2Cell. The GHIST decline and the declines for the other ten HEST models are reported without adjustment. With five patients, the smallest attainable two-sided exact *P* is 0.0625. Analyses of the 45 HEST-Benchmark patients used 20,000 bootstrap draws of patients and two-sided sign-flip tests with 2,000,000 random sign patterns, and the 24 ccRCC patients were tested with the exact sign-flip test.

#### TLS-associated genes

Intervals are 95% percentile intervals from 2,000 resamples of TLS structures for the terms within TLS and of blocks of 10 × 10 spots for the other terms. Values for each cancer type were recomputed in each resample. Intervals for the AUROC of TLS location are 95% percentile intervals from 2,000 resamples of TLS structures with the other spots fixed. A term was evaluated in a section only when the lower limit of the interval of its split-half reliability was above zero. This rule excluded the terms within regions of KC1.

#### Training comparisons

Changes were paired within samples or patients. Intervals are 95% percentile intervals from 20,000 bootstrap draws of the units, stratified by cancer type for HEST, or from 10,000 draws of patients for ccRCC. For HEST, exact two-sided sign-flip tests over all 2^9^ = 512 sample sign patterns were applied to the changes and to the fine-scale change minus the broad-scale change, and are reported without adjustment. The training comparisons of sCellST and DeepSpot2Cell (Supplementary Table 6) were neighboring minus standard and neighboring minus non-neighboring training for each method, and the Phikon-v2 minus the ResNet50 encoder for sCellST. These five comparisons used 20,000 bootstrap draws of patients and exact sign-flip tests over 2^5^ = 32 patterns, with Holm adjustment over all 20 *P* values.

## Data availability

All data analyzed in this study are publicly available. HER2ST data are available from the original study^26^ at https://github.com/almaan/her2st, and the held-out predictions of the 11 methods are available from the benchmark^8^. The breast cancer atlas used for the GHIST cell-type labels and reference expression is available from the original study^3^. The Visium HD colorectal cancer data are available from 10x Genomics (https://www.10xgenomics.com/products/visium-hd-spatial-gene-expression/dataset-human-crc) and from the Gene Expression Omnibus (accession GSE280318)^2^. All 70 HEST-Benchmark samples, the breast cancer Xenium sections and the TENX39 training section are available through HEST-1k (https://huggingface.co/datasets/MahmoodLab/hest) and its benchmark (https://huggingface.co/datasets/MahmoodLab/hest-bench)^27^. The HEST samples, the breast cancer Xenium sections and TENX39 were originally released by 10x Genomics under a Creative Commons Attribution license or by Janesick et al.^62^, and Supplementary Table 7 lists the source of each. The kidney and lung cancer Visium sections with TLS annotations are available from Zenodo (https://doi.org/10.5281/zenodo.14620362). Result tables and figure inputs generated in this study are available in the stRSC repository (Code availability).

## Code availability

The code for RSC, the evaluations in this study and the pairwise difference loss is available at https://github.com/oyjr/stRSC under the MIT license. The repository also contains the scripts that produce the result tables and figures. An interactive website at https://stRSC.org evaluates users’ own predictions and also displays these evaluations for the datasets of this study.

## Acknowledgments

This work was supported by the National Key R&D Program of China (Project No. 2023YFE0204000), Innovation and Technology Commission (Project No. PRP/086/24FX and ITC-SKLNSD26SC01), Shenzhen Science and Technology Innovation Committee Fund (Project No. KCXFZ20230731094059008) and the Research Grants Council of the Hong Kong Special Administrative Region, China (Project No. T45-401/22-N and JLFS/M-604/24).

## Author contributions

J.O. led the development of the evaluation method, implemented the analyses, performed the computational experiments and prepared the figures. Y.G. contributed to method development and data analysis. C.J. helped shape the study and revised the manuscript. Y.W., Y.X., L.L., F.Z. and Z.C. contributed to the interpretation of the results and revision of the manuscript. H.C. supervised the study. J.O. wrote the initial manuscript, with input from the other authors.

---

**Extended Data Fig. 1.**
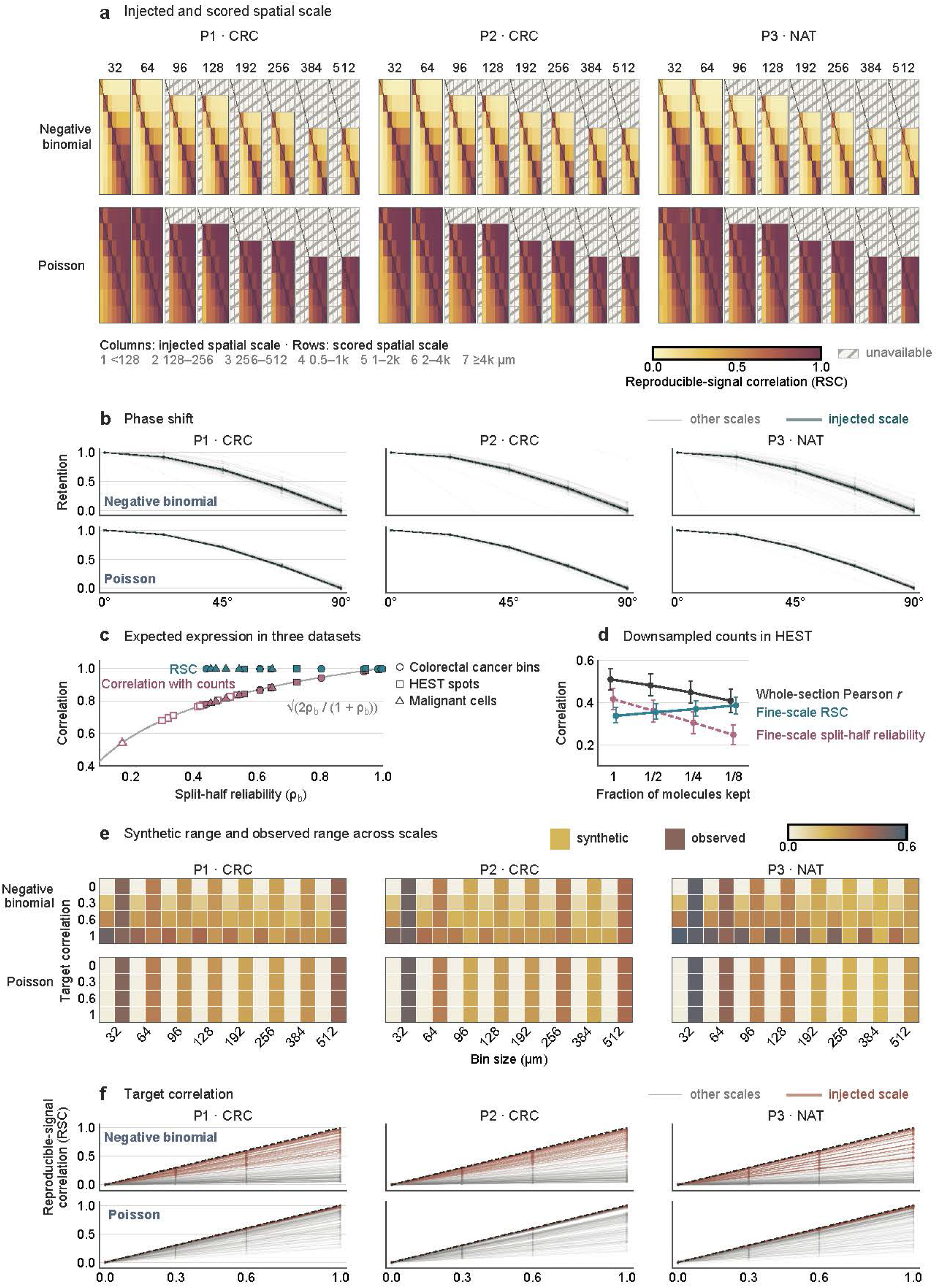
Validation of reproducible-signal correlation with simulated and downsampled counts. a, With the injected patterns as predictions, the reproducible-signal correlation (RSC) is highest at the injected scale in 262 of 264 estimable conditions. Columns show the injected scale, rows show the scored scale, and hatching marks unavailable combinations. **b,** A 90^◦^ shift of the spatial phase at fixed scale and variance lowers the mean absolute RSC to 0.0012. Retention is RSC(*θ*)*/*RSC(0). **c,** The expected expression reaches an RSC near 1 in three datasets, and its correlation with counts follows equation (2). Filled points are means over 20 simulated count matrices, and open points apply equation (2) to the split-half reliability of the measured counts at fine scales. **d,** Downsampling lowers the split-half reliability and whole-section correlation, whereas fine-scale RSC changes from 0.337 to 0.386. Points show means over nine Xenium samples of breast, pancreatic and lung cancer, and error bars show 95% bootstrap intervals. **e,** Range of RSC across scales in the synthetic tests and in colorectal tissue, by target correlation and bin size. **f,** With negative binomial counts, RSC at the injected scale falls below the target at fine scales. Panels **a**, **b**, **e** and **f** use three colorectal patients with Poisson or negative binomial counts at eight bin sizes.

**Extended Data Fig. 2.**
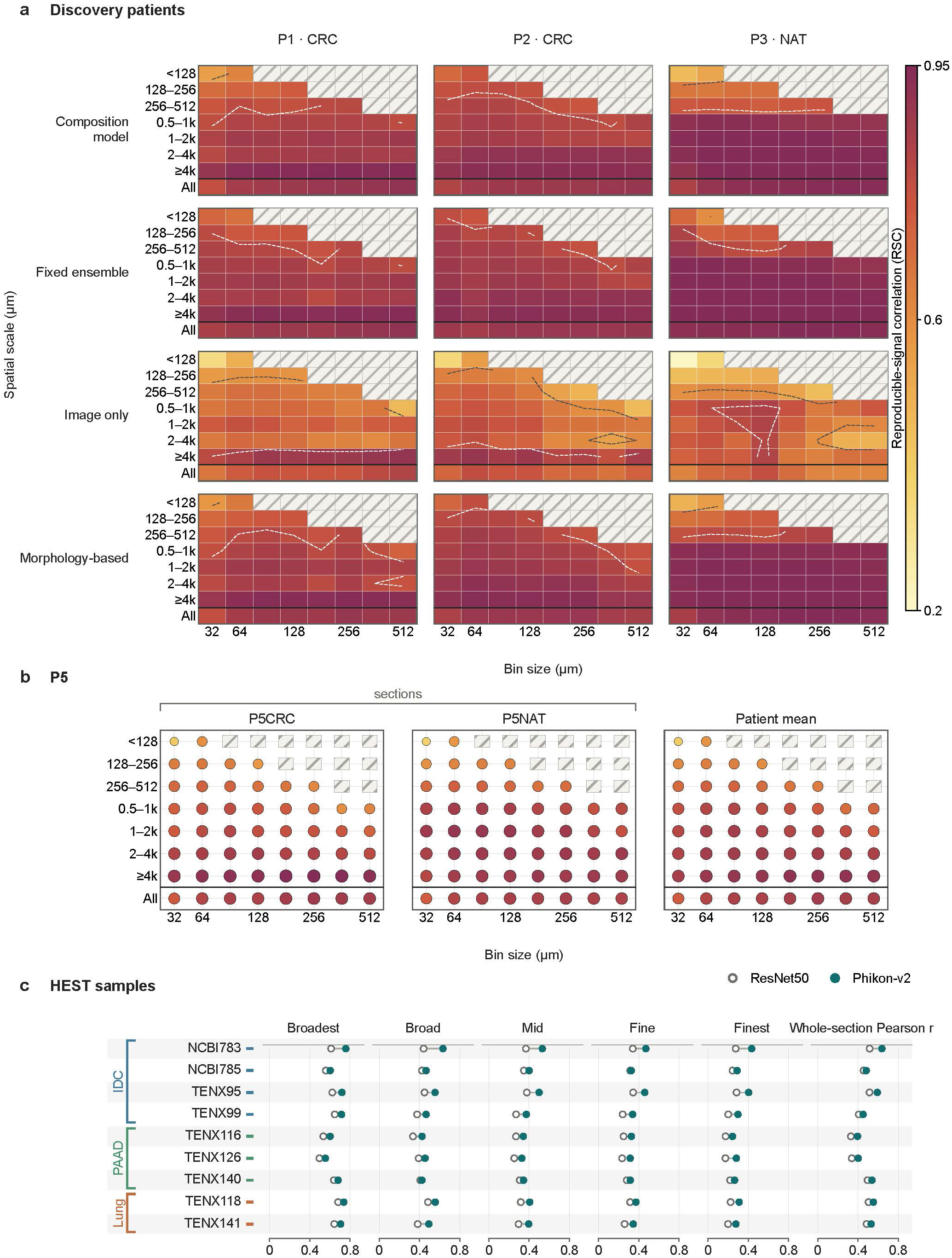
Reproducible-signal correlation across spatial scales in colorectal tissue and in breast, pancreatic and lung cancer. a, Reproducible-signal correlation (RSC) declines toward fine scales for Ridge–Phikon and for three models built partly from RNA-estimated cell-type composition. Scores are shown at seven scales and eight bin sizes in three patients, and hatching marks unavailable combinations. ‘Fixed ensemble’ denotes the fixed scale-specific predictor (Methods). **b,** In a fourth patient, the RSC of Ridge–Phikon declines by a mean of 0.278 from the broadest to the finest scale. **c,** RSC at five scales and whole-section Pearson *r* of ridge regression on ResNet50 or Phikon-v2 features in nine Xenium samples of breast, pancreatic and lung cancer. *n* = 3 patients (**a**), 1 patient (**b**) and 9 samples (**c**).

**Extended Data Fig. 3.**
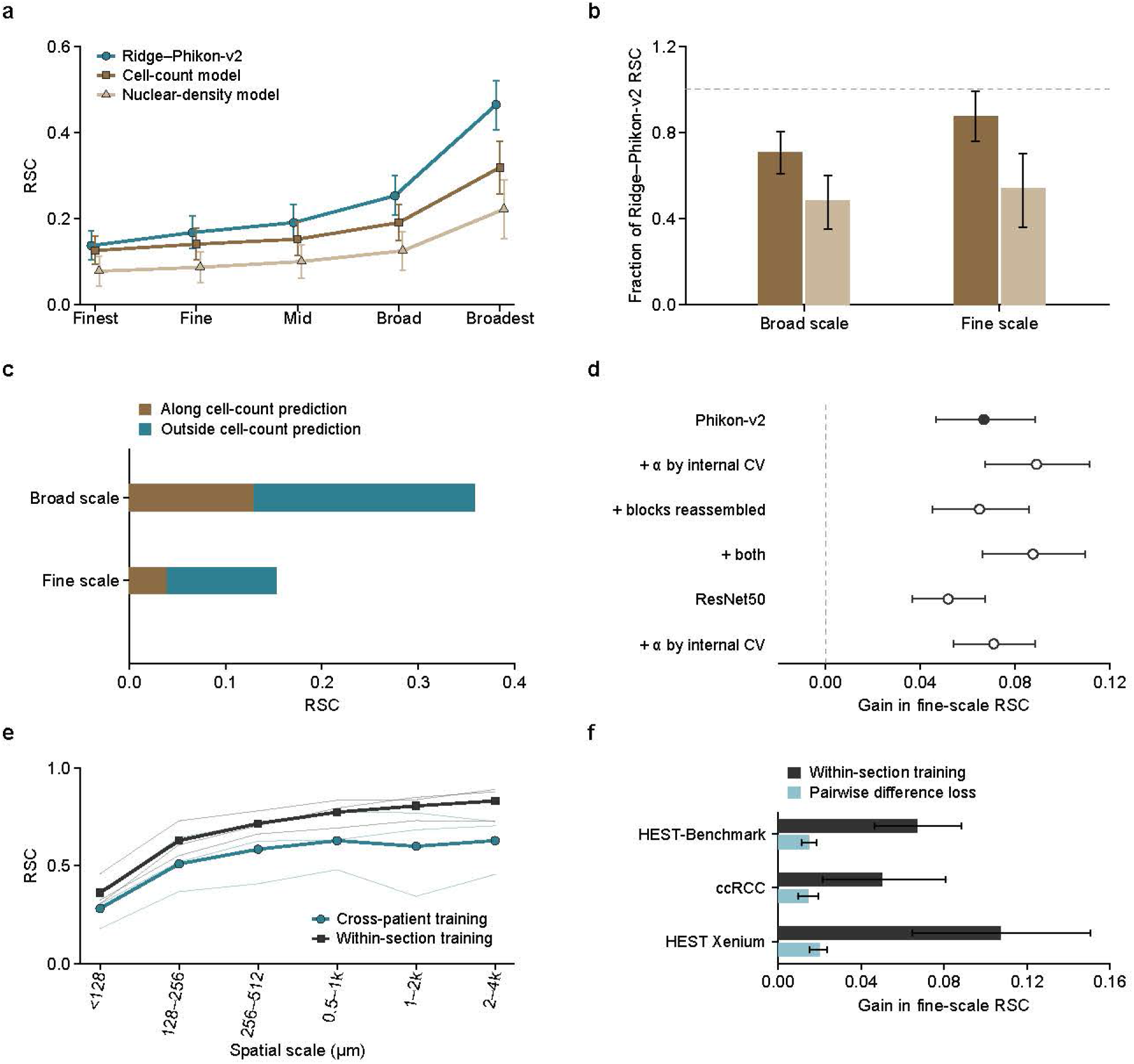
Cell counts, within-section training and the pairwise difference loss in HEST-Benchmark. a, Reproducible-signal correlation (RSC) at five scales for Ridge–Phikon-v2, a cell-count model and a nuclear-density model. The cell-count model uses the number of nuclei of each CellViT class, and the nuclear-density model uses the total number of nuclei. **b,** The cell-count model reaches 0.71 of the broad-scale and 0.87 (95% CI, 0.76 to 0.99) of the fine-scale RSC of Ridge–Phikon-v2. **c,** Image information beyond cell counts accounts for 64% of the broad-scale and 74% (95% CI, 70% to 80%) of the fine-scale RSC of Ridge–Phikon-v2. **d,** Within-section training raises fine-scale RSC by 0.067 (95% CI, 0.047 to 0.088) in the main analysis and by 0.052 to 0.089 in five further settings. **e,** In three colorectal patients, within-section training raises RSC across scales. **f,** The pairwise difference loss recovers 22% (95% CI, 16% to 31%) of the gain from within-section training in HEST-Benchmark, 29% in clear-cell renal cell carcinoma and 19% in Xenium samples of breast, pancreatic and lung cancer. Points and bars show means, thin lines in **e** show patients, and error bars show 95% bootstrap CIs. *n* = 45 patients in **a**–**d**, 3 patients in **e**, and 45 patients, 24 patients and 9 samples in **f**.

**Extended Data Fig. 4.**
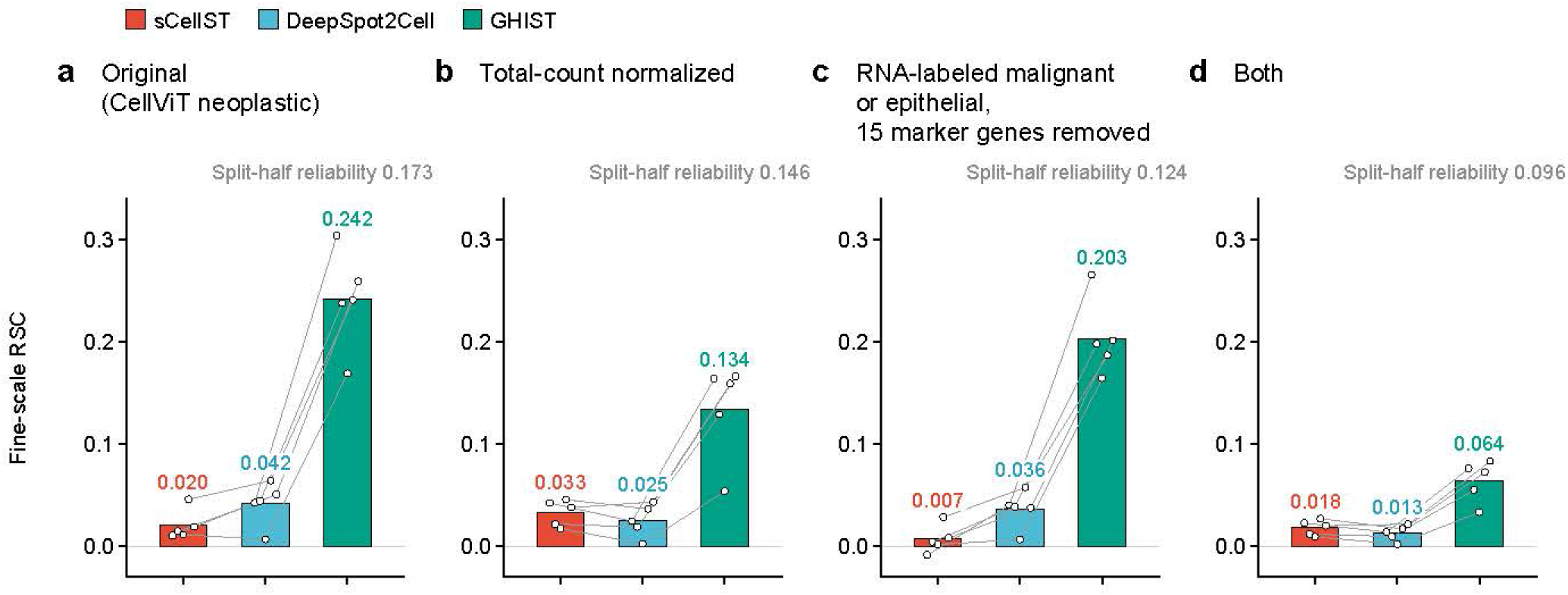
Fine-scale reproducible-signal correlation (RSC) of single-cell methods under four evaluation conditions. a, The original evaluation, in cells assigned to the neoplastic class by CellViT, as in Fig. 4b. **b,** The same cells after normalization by total count. **c,** CellViT neoplastic cells that are also labeled malignant or epithelial from their measured RNA, with 15 epithelial, immune, stromal and endothelial marker genes removed. **d,** The cells and genes of **c** after normalization by total count. Mean fine-scale RSC of GHIST is 0.242, 0.134, 0.203 and 0.064 in **a**–**d**, compared with at most 0.042 for sCellST and DeepSpot2Cell. GHIST is highest in each of the five patients under all four conditions. Numbers above each panel are the split-half reliability. Bars show means of five patients, points show patients, and lines connect the three methods within each patient.

**Extended Data Fig. 5.**
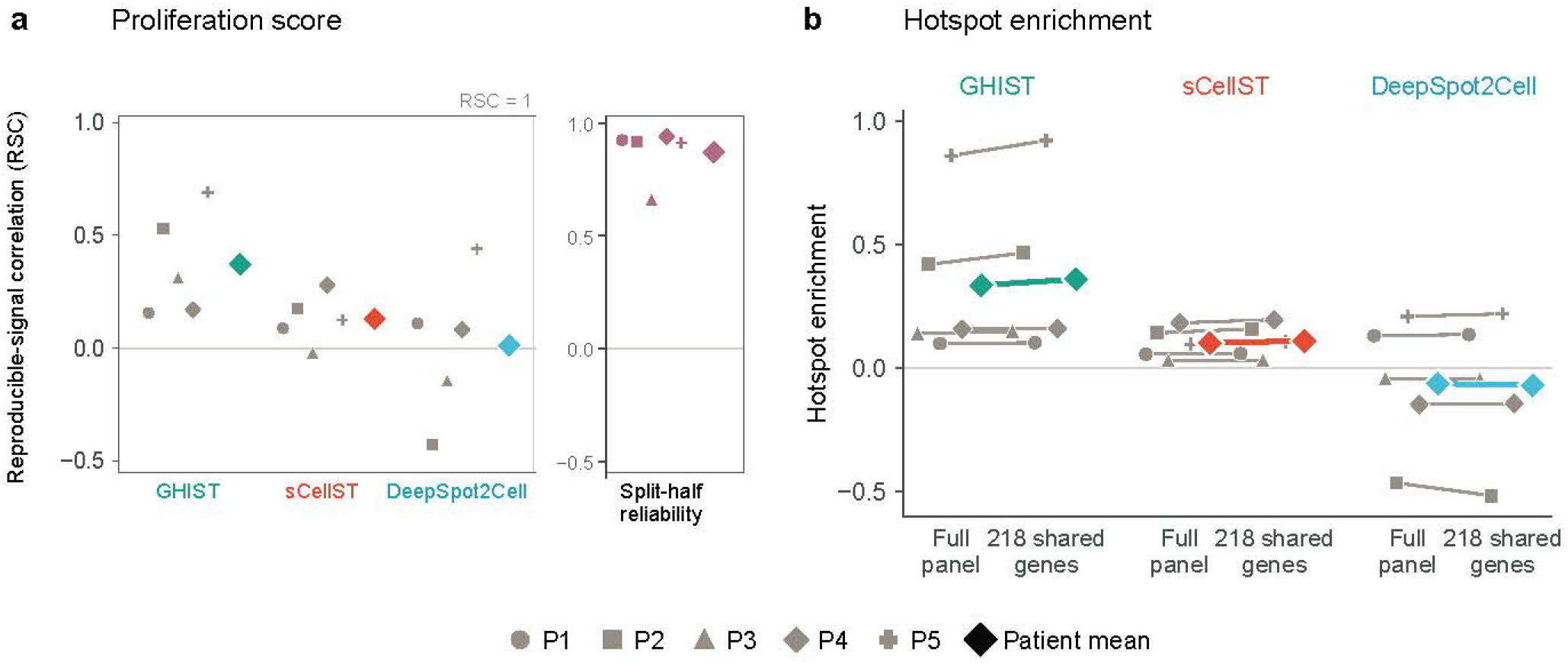
Proliferation scores and hotspots of three single-cell methods on the same cells. a, Reproducible-signal correlation of the proliferation score across malignant-cell neighborhoods is 0.129 for sCellST, 0.012 for DeepSpot2Cell and 0.372 for GHIST, with a split-half reliability of 0.869. **b,** Hotspot enrichment for GHIST is positive in all five patients with the measured score normalized over the full panel as in Fig. 4e or over the 218 shared genes. Lines connect the two normalizations within each patient, and diamonds show means (*n* = 5 patients). Both panels use the cells shared by the three methods, as in Fig. 4.

**Extended Data Fig. 6.**
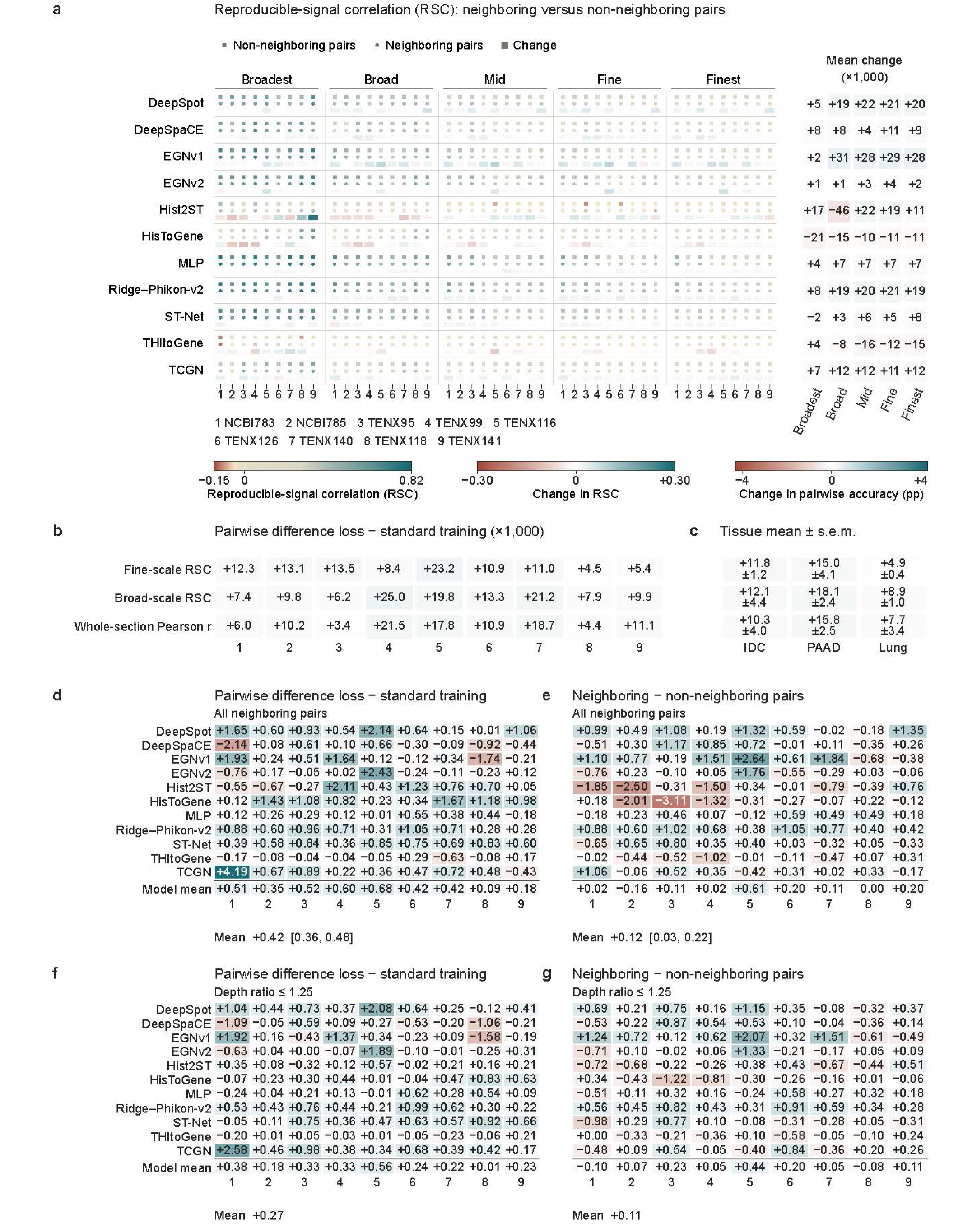
Effects of the pairwise difference loss by scale and on pairwise accuracy between neighboring spots. a, Reproducible-signal correlation (RSC) at five scales with neighboring and non-neighboring pairs for 11 models in nine Xenium samples of breast, pancreatic and lung cancer. Margins show mean changes (×1,000). **b,** Changes from standard training to the pairwise difference loss in fine-scale RSC, broad-scale RSC and whole-section Pearson *r*, averaged over 11 models (×1,000). Mean changes are 0.011, 0.013 and 0.012. **c,** The same changes by tissue (mean and s.e.m.) for invasive ductal carcinoma (IDC), pancreatic adenocarcinoma (PAAD) and lung cancer. **d,e,** Change in pairwise accuracy between neighboring spots relative to standard training (**d**) and to non-neighboring pairs (**e**), in percentage points. **f,g,** The same comparisons for pairs whose total counts are within a ratio of 1.25. Values in brackets are 95% bootstrap intervals. *n* = 9 samples.

**Extended Data Fig. 7.**
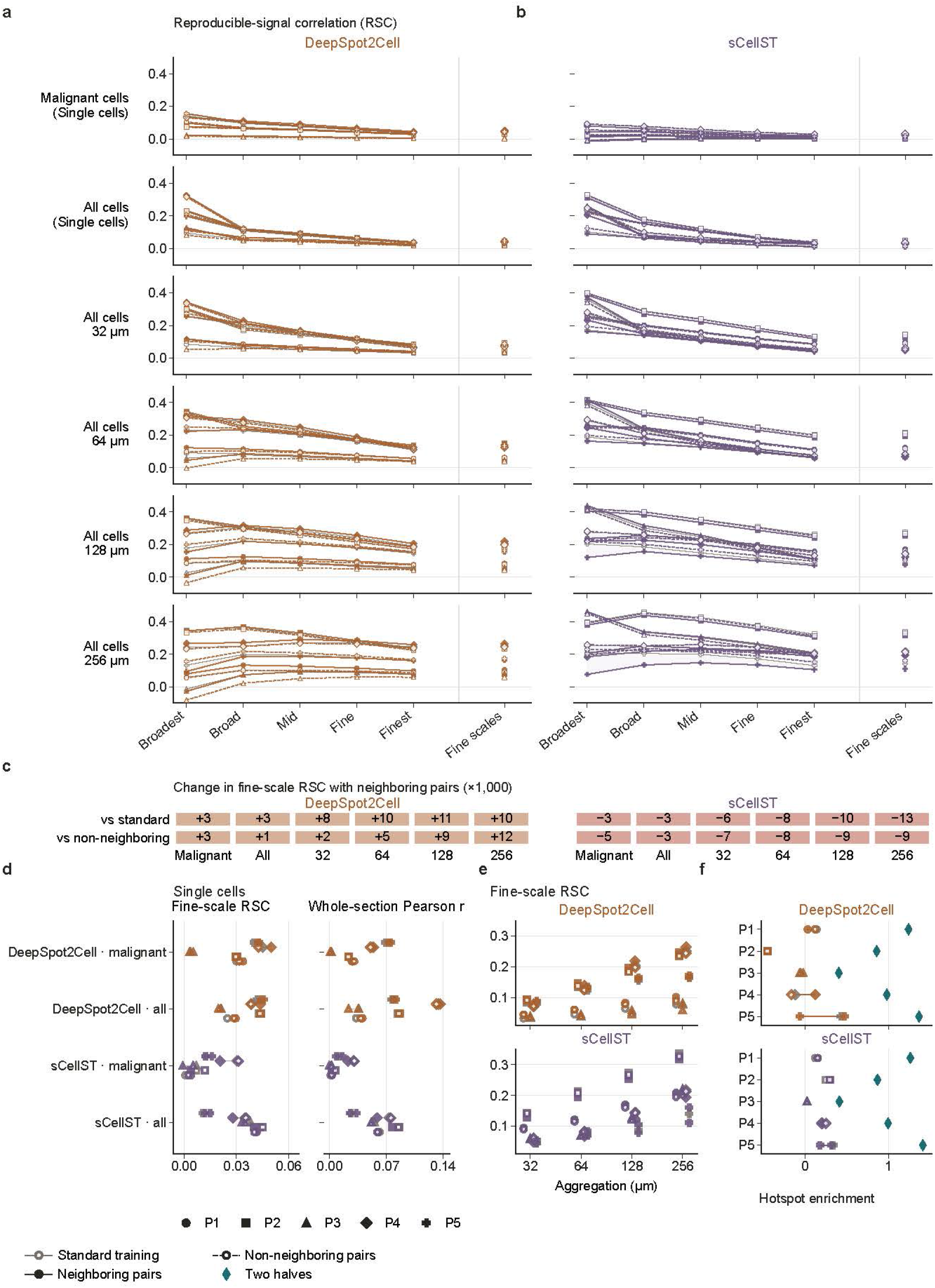
The pairwise difference loss in the two single-cell methods trained with spot-level supervision. a,b, Reproducible-signal correlation (RSC) of DeepSpot2Cell (**a**) and sCellST (**b**) after standard training and with the pairwise difference loss on neighboring or non-neighboring pairs. **c,** Mean change in fine-scale RSC with neighboring pairs relative to standard training and to non-neighboring pairs (×1,000). The changes are small and of opposite sign in the two methods, and no change is significant after Holm adjustment (Supplementary Table 6). **d,** Fine-scale RSC and whole-section Pearson *r* by patient. **e,** Fine-scale RSC after aggregation to 32–256 *µ*m. **f,** Measured enrichment at proliferation hotspots predicted in each training condition, with hotspots of the two RNA halves as reference. All panels use each method’s own cells and genes (*n* = 5 patients, nine sections).

**Extended Data Table 1.** Predictions evaluated in each analysis.

| Analysis | Data | Predictions | Shown in |
|---|---|---|---|
| Pairwise accuracy within regions | HER2ST; 8 patients; 785 genes | 11 published methods; class-mean baseline | Fig. 2; Supplementary Table 1; Supplementary Note 1 |
| Pairwise accuracy after matching | HEST; 9 samples; 50 genes | 11 HEST models | Supplementary Fig. 1; Supplementary Table 2 |
| Spatial scales | CRC; 3 discovery patients and a fourth patient; 1,000 genes (17,977 for the error decomposition in Fig. 3b) | Ridge–Phikon; three models built partly from RNA-estimated composition | Fig. 3a–c; Extended Data Fig. 2a,b; Supplementary Table 3 |
| Spatial scales | HEST; 9 samples; 50 genes | 11 HEST models after standard training; Ridge–ResNet50 | Fig. 3d; Extended Data Fig. 2c; Supplementary Table 3 |
| Spatial scales | HEST-Benchmark; 70 samples from 45 patients; 50 genes per task | Ridge regression on five image encoders; cell-count and nuclear-density models; Ridge–Phikon-v2 with within-section training and with the pairwise difference loss | Fig. 3e,f; Extended Data Fig. 3; Supplementary Table 4 |
| Cell classes | Xenium; 9 sections from 5 patients; 201–211 genes | sCellST, DeepSpot2Cell and GHIST; class-mean baseline on the cells and genes of sCellST | Fig. 4; Extended Data Fig. 4; Supplementary Table 3; Supplementary Note 1 |
| TLS-associated genes | External Visium; 2 kidney and 2 lung sections; 8 genes | Ridge–Phikon-v2 with and without the pairwise difference loss and trained within one cancer type; class-mean baseline | Fig. 5a–f; Supplementary Table 5; Supplementary Notes 1 and 3 |
| Proliferation hotspots | Xenium; 5 patients | sCellST, DeepSpot2Cell and GHIST | Fig. 4e; Extended Data Fig. 5 |
| Pairwise difference loss | HEST; 9 samples | 11 HEST models compared with standard training and with non-neighboring pairs | Fig. 6; Extended Data Fig. 6; Supplementary Note 4 |
| Pairwise difference loss | ccRCC; 24 patients | MLP; ridge model for the proliferation score | Supplementary Fig. 2; Supplementary Note 4 |
| Pairwise difference loss | Xenium; 9 sections from 5 patients | sCellST and DeepSpot2Cell | Extended Data Fig. 7; Supplementary Table 6 |

The 11 published methods are DeepPT, DeepSpaCE, EGNv1, EGNv2, GeneCodeR, HisToGene, Hist2ST, ST-Net, TCGN, THItoGene and iStar, which was evaluated at spot resolution; with DeepSpot they form the 12 spot-level methods counted in the text. The 11 HEST models are Ridge–Phikon-v2, the MLP, ST-Net, HisToGene, Hist2ST, THItoGene, DeepSpot, TCGN, DeepSpaCE, EGNv1 and EGNv2; all 11 were trained with and without the pairwise difference loss. Ridge–Phikon, Ridge–Phikon-v2, Ridge–ResNet50 and the MLP are generic models and are not counted among the 12 methods. Data and methods are described in Methods, and identifiers are listed in Supplementary Table 7.

## References

1. Ståhl, P. L. et al. Visualization and analysis of gene expression in tissue sections by spatial transcriptomics. Science 353, 78–82, DOI: 10.1126/science.aaf2403 (2016).

2. de Oliveira, M. F. et al. High-definition spatial transcriptomic profiling of immune cell populations in colorectal cancer. Nat. Genet. 57, 1512–1523, DOI: 10.1038/s41588-025-02193-3 (2025).

3. Wu, S. Z. et al. A single-cell and spatially resolved atlas of human breast cancers. Nat. Genet. 53, 1334–1347, DOI: 10.1038/s41588-021-00911-1 (2021).

4. He, B. et al. Integrating spatial gene expression and breast tumour morphology via deep learning. *Nat*. Biomed. Eng. 4, 827–834, DOI: 10.1038/s41551-020-0578-x (2020).

5. Fu, X. et al. Spatial gene expression at single-cell resolution from histology using deep learning with GHIST. Nat. Methods 22, 1900–1910, DOI: 10.1038/s41592-025-02795-z (2025).

6. Schroeder, A. et al. Scaling up spatial transcriptomics for large-sized tissues: uncovering cellular-level tissue architecture beyond conventional platforms with iSCALE. Nat. Methods 22, 1911–1922, DOI: 10.1038/s41592-025-02770-8 (2025).

7. Ouyang, J. et al. GenAR: next-scale autoregressive generation for spatial gene expression prediction. Med. Image Anal. 114, 104232, DOI: 10.1016/j.media.2026.104232 (2026).

8. Wang, C. et al. Benchmarking the translational potential of spatial gene expression prediction from histology. Nat. Commun. 16, 1544, DOI: 10.1038/s41467-025-56618-y (2025).

9. Xie, J. et al. Performance of deep learning methods for spatial gene expression prediction from histology images: a comprehensive assessment. Bioinform. Adv. 6, vbag202, DOI: 10.1093/bioadv/vbag202 (2026).

10. Gindra, R. H. et al. A large-scale benchmark of cross-modal learning for histology and gene expression in spatial transcriptomics. In Proceedings of the IEEE/CVF International Conference on Computer Vision Workshops, 1182–1192 (2025). https://openaccess.thecvf.com/content/ICCV2025W/CVAMD/html/Gindra_A_Large-Scale_Benchmark_of_Cross-Modal_Learning_for_Histology_and_Gene_ICCVW_2025_paper.html.

11. Brennecke, P. et al. Accounting for technical noise in single-cell RNA-seq experiments. Nat. Methods 10, 1093–1095, DOI: 10.1038/nmeth.2645 (2013).

12. Grün, D., Kester, L. & van Oudenaarden, A. Validation of noise models for single-cell transcriptomics. Nat. Methods 11, 637–640, DOI: 10.1038/nmeth.2930 (2014).

13. You, Y. et al. Systematic comparison of sequencing-based spatial transcriptomic methods. Nat. Methods 21, 1743–1754, DOI: 10.1038/s41592-024-02325-3 (2024).

14. Vickovic, S. et al. High-definition spatial transcriptomics for in situ tissue profiling. Nat. Methods 16, 987–990, DOI: 10.1038/s41592-019-0548-y (2019).

15. Saxton, W. O. & Baumeister, W. The correlation averaging of a regularly arranged bacterial cell envelope protein. J. Microsc. 127, 127–138, DOI: 10.1111/j.1365-2818.1982.tb00405.x (1982).

16. van Heel, M. Similarity measures between images. Ultramicroscopy 21, 95–100, DOI: 10.1016/0304-3991(87)90010-6 (1987).

17. Scheres, S. H. W. & Chen, S. Prevention of overfitting in cryo-EM structure determination. Nat. Methods 9, 853–854, DOI: 10.1038/nmeth.2115 (2012).

18. Nieuwenhuizen, R. P. J. et al. Measuring image resolution in optical nanoscopy. Nat. Methods 10, 557–562, DOI: 10.1038/nmeth.2448 (2013).

19. Fried, D. L. Noise in photoemission current. Appl. Opt. 4, 79–80, DOI: 10.1364/AO.4.000079 (1965).

20. Batson, J., Royer, L. & Webber, J. Molecular cross-validation for single-cell RNA-seq. bioRxiv 786269, DOI: 10.1101/786269 (2019).

21. Neufeld, A. C. et al. Inference after latent variable estimation for single-cell RNA sequencing data. Biostatistics 25, 270–287, DOI: 10.1093/biostatistics/kxac047 (2024).

22. Lage-Castellanos, A., Valente, G., Formisano, E. & De Martino, F. Methods for computing the maximum performance of computational models of fMRI responses. PLoS Comput. Biol. 15, e1006397, DOI: 10.1371/journal.pcbi.1006397 (2019).

23. Spearman, C. The proof and measurement of association between two things. Am. J. Psychol. 15, 72–101, DOI: 10.2307/1412159 (1904).

24. Gell, M. et al. How measurement noise limits the accuracy of brain–behaviour predictions. Nat. Commun. 15, 10678, DOI: 10.1038/s41467-024-54022-6 (2024).

25. Keshishian, M. et al. Estimating and interpreting nonlinear receptive field of sensory neural responses with deep neural network models. eLife 9, e53445, DOI: 10.7554/eLife.53445 (2020).

26. Andersson, A. et al. Spatial deconvolution of HER2-positive breast cancer delineates tumor-associated cell type interactions. Nat. Commun. 12, 6012, DOI: 10.1038/s41467-021-26271-2 (2021).

27. Jaume, G., et al. HEST-1k: a dataset for spatial transcriptomics and histology image analysis. In Advances in Neural Information Processing Systems, vol. 37, 53798–53833, DOI: 10.52202/079017-1704 (2024).

28. Chang, Y. et al. Graph Fourier transform for spatial omics representation and analyses of complex organs. Nat. Commun. 15, 7467, DOI: 10.1038/s41467-024-51590-5 (2024).

29. Chadoutaud, L. et al. sCellST predicts single-cell gene expression from H&E images. Nat. Commun. 17, 1194, DOI: 10.1038/s41467-025-67965-1 (2026).

30. Filiot, A., et al. Phikon-v2, a large and public feature extractor for biomarker prediction. arXiv DOI: 10.48550/arXiv.2409.09173 (2024).

31. Filiot, A. et al. Scaling self-supervised learning for histopathology with masked image modeling. medRxiv DOI: 10.1101/2023.07.21.23292757 (2023).

32. Hörst, F. et al. CellViT: vision transformers for precise cell segmentation and classification. Med. Image Anal. 94, 103143, DOI: 10.1016/j.media.2024.103143 (2024).

33. Nonchev, K., Manaiev, G., Koelzer, V. H. & Rätsch, G. DeepSpot2Cell: predicting virtual single-cell spatial transcriptomics from H&E images using spot-level supervision. bioRxiv DOI: 10.1101/2025.09.23.678121 (2025).

34. Robinson, W. S. Ecological correlations and the behavior of individuals. Am. Sociol. Rev. 15, 351–357, DOI: 10.2307/2087176 (1950).

35. Meylan, M. et al. Tertiary lymphoid structures generate and propagate anti-tumor antibody-producing plasma cells in renal cell cancer. Immunity 55, 527–541.e5, DOI: 10.1016/j.immuni.2022.02.001 (2022).

36. Nonchev, K. et al. DeepSpot: leveraging spatial context for enhanced spatial transcriptomics prediction from H&E images. medRxiv DOI: 10.1101/2025.02.09.25321567 (2025).

37. Dawo, S., Nonchev, K. & Silina, K. 10x Visium spatial transcriptomics dataset: kidney (3) and lung (5) cancer with tertiary lymphoid structures. Zenodo, DOI: 10.5281/zenodo.14620362 (2025).

38. Zhu, D., Yang, T. & Jerby, L. Gradient aligned regression via pairwise losses. In Proceedings of the 42nd International Conference on Machine Learning, vol. 267 of Proceedings of Machine Learning Research, 80259–80281 (PMLR, 2025). https://proceedings.mlr.press/v267/zhu25x.html.

39. Nishimura, K., Hirose, H., Bise, R., Shiku, K. & Kojima, Y. Learning relative gene expression trends from pathology images in spatial transcriptomics. In Advances in Neural Information Processing Systems, vol. 38, 86684–86705 (2025). https://papers.nips.cc/paper_files/paper/2025/file/7d535a224c8ae54ba75bac0457b6b279-Paper-Conference.pdf.

40. Byeon, K., Park, S., Lim, J. & Kwak, J. T. COAST: context-aware differential learning for gene expression prediction in spatial transcriptomics. arXiv DOI: 10.48550/arXiv.2607.09166 (2026).

41. Miller, H. E. et al. Deep learning perturbation models can outperform baselines on calibrated metrics. Nat. Biotechnol. DOI: 10.1038/s41587-026-03307-w (2026).

42. Hsu, A., Borst, A. & Theunissen, F. E. Quantifying variability in neural responses and its application for the validation of model predictions. Network: Comput. Neural Syst. 15, 91–109 (2004).

43. Schoppe, O., Harper, N. S., Willmore, B. D. B., King, A. J. & Schnupp, J. W. H. Measuring the performance of neural models. Front. Comput. Neurosci. 10, 10, DOI: 10.3389/fncom.2016.00010 (2016).

44. Kazemian, A., Elmoznino, E. & Bonner, M. F. Convolutional architectures are cortex-aligned de novo. *Nat*. Mach. Intell. 7, 1834–1844, DOI: 10.1038/s42256-025-01142-3 (2025).

45. Doerig, A. et al. High-level visual representations in the human brain are aligned with large language models. *Nat*. Mach. Intell. 7, 1220–1234, DOI: 10.1038/s42256-025-01072-0 (2025).

46. Sun, E. D., Ma, R., Navarro Negredo, P., Brunet, A. & Zou, J. TISSUE: uncertainty-calibrated prediction of single-cell spatial transcriptomics improves downstream analyses. Nat. Methods 21, 444–454, DOI: 10.1038/s41592-024-02184-y (2024).

47. Jin, K. et al. Multiscale confidence quantification for virtual spatial transcriptomics with UTOPIA. bioRxiv DOI: 10.64898/2026.03.01.708850 (2026).

48. Angelopoulos, A. N., Bates, S., Fannjiang, C., Jordan, M. I. & Zrnic, T. Prediction-powered inference. Science 382, 669–674, DOI: 10.1126/science.adi6000 (2023).

49. Zhang, D. et al. Inferring super-resolution tissue architecture by integrating spatial transcriptomics with histology. Nat. Biotechnol. 42, 1372–1377, DOI: 10.1038/s41587-023-02019-9 (2024).

50. Moffitt, J. R., Lundberg, E. & Heyn, H. The emerging landscape of spatial profiling technologies. Nat. Rev. Genet. 23, 741–759, DOI: 10.1038/s41576-022-00515-3 (2022).

51. Stickels, R. R. et al. Highly sensitive spatial transcriptomics at near-cellular resolution with Slide-seqV2. Nat. Biotechnol. 39, 313–319, DOI: 10.1038/s41587-020-0739-1 (2021).

52. Rosenthal, P. B. & Henderson, R. Optimal determination of particle orientation, absolute hand, and contrast loss in single-particle electron cryomicroscopy. J. Mol. Biol. 333, 721–745, DOI: 10.1016/j.jmb.2003.07.013 (2003).

53. Hoang, D.-T. et al. A deep-learning framework to predict cancer treatment response from histopathology images through imputed transcriptomics. *Nat*. Cancer 5, 1305–1317, DOI: 10.1038/s43018-024-00793-2 (2024).

54. Monjo, T. et al. Efficient prediction of a spatial transcriptomics profile better characterizes breast cancer tissue sections without costly experimentation. Sci. Rep. 12, 4133, DOI: 10.1038/s41598-022-07685-4 (2022).

55. Yang, Y., Hossain, M. Z., Stone, E. A. & Rahman, S. Exemplar guided deep neural network for spatial transcriptomics analysis of gene expression prediction. In IEEE/CVF Winter Conference on Applications of Computer Vision, 5028–5037, DOI: 10.1109/WACV56688.2023.00501 (2023).

56. Yang, Y., Hossain, M. Z., Stone, E. & Rahman, S. Spatial transcriptomics analysis of gene expression prediction using exemplar guided graph neural network. Pattern Recognit. 145, 109966, DOI: 10.1016/j.patcog.2023.109966 (2024).

57. Banh, D. & Huang, A. Scalable parametric encoding of multiple modalities. bioRxiv DOI: 10.1101/2021.07.09.451779 (2022).

58. Pang, M., Su, K. & Li, M. Leveraging information in spatial transcriptomics to predict super-resolution gene expression from histology images in tumors. bioRxiv DOI: 10.1101/2021.11.28.470212 (2021).

59. Zeng, Y. et al. Spatial transcriptomics prediction from histology jointly through Transformer and graph neural networks. Brief. Bioinform. 23, bbac297, DOI: 10.1093/bib/bbac297 (2022).

60. Xiao, X., Kong, Y., Li, R., Wang, Z. & Lu, H. Transformer with convolution and graph-node co-embedding: an accurate and interpretable vision backbone for predicting gene expressions from local histopathological image. Med. Image Anal. 91, 103040, DOI: 10.1016/j.media.2023.103040 (2024).

61. Jia, Y. et al. THItoGene: a deep learning method for predicting spatial transcriptomics from histological images. Brief. Bioinform. 25, bbad464, DOI: 10.1093/bib/bbad464 (2024).

62. Janesick, A. et al. High resolution mapping of the tumor microenvironment using integrated single-cell, spatial and in situ analysis. Nat. Commun. 14, 8353, DOI: 10.1038/s41467-023-43458-x (2023).

63. He, K., Zhang, X., Ren, S. & Sun, J. Deep residual learning for image recognition. In IEEE Conference on Computer Vision and Pattern Recognition, 770–778, DOI: 10.1109/CVPR.2016.90 (2016).

64. Chen, R. J. et al. Towards a general-purpose foundation model for computational pathology. Nat. Med. 30, 850–862, DOI: 10.1038/s41591-024-02857-3 (2024).

65. Karasikov, M. et al. Training state-of-the-art pathology foundation models with orders of magnitude less data. arXiv DOI: 10.48550/arXiv.2504.05186 (2025).

66. Zimmermann, E., et al. Virchow2: scaling self-supervised mixed magnification models in pathology. arXiv DOI: 10.48550/arXiv.2408.00738 (2024).

67. Pedregosa, F. et al. Scikit-learn: machine learning in Python. J. Mach. Learn. Res. 12, 2825–2830 (2011). https://jmlr.org/papers/v12/pedregosa11a.html.

68. Squair, J. W. et al. Confronting false discoveries in single-cell differential expression. Nat. Commun. 12, 5692, DOI: 10.1038/s41467-021-25960-2 (2021).

69. Graham, S. et al. Hover-Net: simultaneous segmentation and classification of nuclei in multi-tissue histology images. Med. Image Anal. 58, 101563, DOI: 10.1016/j.media.2019.101563 (2019).

70. Lin, Y. et al. scClassify: sample size estimation and multiscale classification of cells using single and multiple reference. Mol. Syst. Biol. 16, e9389, DOI: 10.15252/msb.20199389 (2020).

