## Supplementary Information for "A noise-corrected metric for evaluating spatial transcriptomics predicted from histology"

**Contents**

Supplementary Methods
Supplementary Note 1: Class-mean baseline
Supplementary Note 2: Derivation of the reproducible-signal correlation
Supplementary Note 3: Tertiary lymphoid structures
Supplementary Note 4: Training comparisons
Supplementary Note 5: Sensitivity analyses for pairwise accuracy
Supplementary Figures 1–3
Supplementary Tables 1–7

### Supplementary Methods

These sections complete the Methods of the main text and follow its order.

##### Datasets and evaluated methods

*Kidney and lung cancer sections.* Feature standardization and the principal component analysis were fitted within each training fold. Ridge targets were $\log(1+10{,}000 N_{ig}/\sum_g N_{ig})$, where $N_{ig}$ is the count of gene $g$ in spot $i$ and the sum runs over all genes. The regularization strength was $100/(256\times 748)$.

##### Evaluation between and within tissue regions

*Spot pairs.* To match pairs for a target gene, genes were divided into ten folds, and the fold containing the target gene was excluded. The source-half counts of the remaining genes were normalized to 10,000 per spot and log-transformed. Variable genes entered a principal component analysis with at most 20 components, and each spot was joined to up to eight nearest neighbors in this space. Pairs were kept when their source-half counts, summed over the genes used for matching, were within a ratio of 1.25. The closest quarter of these pairs was used, and pairs were selected from this quarter in order of increasing distance without reusing a spot. A gene was eligible when all methods had predictions for it at 20 or more spots and, in the source half, it had at least 20 molecules and was detected at 5% or more of spots. For each eligible gene, pairs were kept when their absolute source-half expression difference was at or above the 75th percentile. Each gene required at least ten candidate pairs for this step. Unmatched pairs followed the same percentile rule. Distance quartiles were defined from all spot pairs of a section. The number of unmatched pairs was capped at the number of matched pairs within each gene, split, direction and distance quartile.

*Spearman correlation and all-pair concordance.* These analyses used spots with predictions from all 11 methods and a region annotation, regions with at least 20 such spots, and genes with at least 20 molecules in the source half and detection in at least 5% of spots. Scores used twice the evaluation-half counts under both assignments of the halves. For concordance, pairs tied in the evaluation half were excluded. Within regions, scores were averaged with equal weights over regions, then over assignments of the halves, genes, count splits, methods and patients.

*Depth-normalized directions.* In a sensitivity analysis, pairwise accuracy in HER2ST, in HEST and between neighboring spots was recomputed with measured directions taken from depth-normalized counts. Pairs and predictions were unchanged. Each half-count was divided by the total count of that half at the spot. In HER2ST, this total was summed over the 785 shared genes. The score of the two halves and the majority-order baseline used the same normalized directions. For pairs of neighboring spots, the pairs selected in the 32 count splits of the main analysis were kept, and the unsplit count was divided by the unsplit total count of the spot. Results are given in Supplementary Note 5 and Supplementary Table 2.

##### Tests of the reproducible-signal correlation

*Resampling of the measured HEST counts.* Each measured count of the nine HEST samples was treated as a Poisson rate, 64 count matrices were drawn per sample, and each matrix was split once. Synthetic predictions with target correlations of 0, 0.3, 0.316, 0.6, 0.609 and 1 were scored on these matrices, and each score was compared with the same score computed without sampling noise. Across the 324 conditions, the scores deviated from their values without sampling noise by at most 0.0109 (Supplementary Fig. 3b). The same simulated counts were used to rescore the 306 sets of predictions from the training comparisons. The mean gains of these comparisons remained positive (Supplementary Fig. 3c,d). Intervals were computed from 2,000 resamples of the 64 matrices.

*Scoring on cell graphs.* The scoring procedure was tested on 15 cell graphs, the malignant-cell graphs of the nine breast cancer sections and graphs of section NCBI784 under six further settings. When the two halves had identical counts and the prediction equaled the transformed half-counts, all 90 scores deviated from 1 by less than $10^{-5}$. With 16 independent random predictions per graph, 87 of 90 mean scores were consistent with zero after Bonferroni adjustment. The other three mean scores, all on one graph, were about 0.001.

##### Evaluation across spatial scales

*Cell-count model.* Nuclei came from the CellViT segmentations distributed with HEST-1k and were counted within 27.5 $\mu$m of each Visium spot center or within each 100-$\mu$m Xenium square. For the combined model, the five nuclear counts were appended to the 256 principal components of the Phikon-v2 features, with the same regularization strength $\alpha$. On matched pairs, both models were scored on the same pairs, built as described under 'HEST matched pairs' in Methods, with tied predictions given half credit.

*Within-section training.* The five further settings were selection of $\alpha$ by internal cross-validation over 1 to $10^5$ times its value in the main analysis, scoring of the four blocks together on the whole section, both changes together, and ResNet50 features with either choice of $\alpha$.

###### Evaluation within cell classes

*GHIST training.* Nuclei were segmented from the H&E images with HoVer-Net[69] and matched to Xenium cells by maximum overlap. A match required the overlap to cover at least 50% of the nucleus area, and each Xenium cell was assigned the nucleus with the largest overlap. Training nuclei also required an area of at least 10 $\mu m^2$. Cell types of training cells were assigned with scClassify[70] using an independent breast cancer atlas[3] and mapped to the GHIST classes. Of the 100,064 atlas cells, the 92,795 in cell types corresponding to these classes were used, and the cell-type expression profiles were computed from these cells.

*Further conditions.* For Extended Data Fig. 4, the evaluation was repeated under three further conditions. In the first, each half-count was transformed as $\log(1+100N_{\text{half}}/T_{\text{half}})$, where $T_{\text{half}}$ is the total count of the cell in that half over the full panel. Predictions were converted back to counts and normalized in the same way by their total over the shared genes. In the second, the analysis used the CellViT neoplastic cells that the scClassify labels described under 'GHIST training' assigned to malignant or epithelial types, and the cell graph was rebuilt on these cells. This condition also removed 15 epithelial, immune, stromal and endothelial marker genes among the 218 shared genes. The third condition combined both changes.

###### Evaluation of TLS-associated genes

*RNA-based checks.* The 21 transformed versions of the annotation in the second RNA-based check were shifts by one to three spots in each of the six directions of the Visium grid, a rotation by 180° and mirror images about the horizontal and vertical axes.

###### Training with the pairwise difference loss

*Pairwise difference loss.* The training graph was fixed before optimization, and each model drew edges according to its batching scheme. Ridge used all edges in its design matrix, and HisToGene and THItoGene used all training edges in each full-section update. The MLP and Hist2ST iterated over all training edges once per epoch, and ST-Net paired each training item with one edge from a separately shuffled list. DeepSpot used at most 512 shuffled edges per update, and TCGN drew 32 edges per update.

*Pairwise accuracy between neighboring spots.* For Extended Data Fig. 6d–g, spots within 1.5 times the median nearest-neighbor distance were joined in each held-out sample, and the counts were split 32 times. A gene and pair were included when both RNA halves showed the same non-zero direction. A prediction scored 1 for the same direction, 0 for the opposite direction and 0.5 for a tie. Scores were summed over count splits within each gene, and genes were weighted equally within a sample. Changes between training conditions are given in percentage points. For Extended Data Fig. 6f,g and Supplementary Table 2c, the analysis was repeated with the pairs whose total counts were within a ratio of 1.25.

*ccRCC.* These analyses used the six outer folds provided by the benchmark and the 73,813 spots whose image patches were matched to expression barcodes. Each outer fold had four test patients. The models used frozen Phikon-v2 features and were trained on $\log(1+N)$, where $N$ is the count of each gene at each spot. Within each outer fold, four of the 20 non-test patients were held out for validation, and the other 16 formed the training set. Nested training subsets of 1, 2, 4, 8 and 16 patients were drawn from this set in one fixed order. The model was an MLP with a 256-dimensional input, one hidden layer of 512 units and 50 outputs, trained with AdamW for 100 epochs. Standard training (condition S) minimized the mean squared error at each spot. Condition D gave equal weights to this error and to the pairwise difference term, and condition A added a logistic ranking loss on the predicted changes in expression. A control repeated standard training with 1,024 training spots in total. Outcomes were the gene-wise Pearson correlation between predictions and measured expression, the Pearson correlation between predicted and measured changes in expression between neighboring spots and the mean squared error. Checkpoints were chosen by validation mean squared error, and the results are shown in Supplementary Fig. 2. The change reported for the pairwise difference loss was condition D minus condition S with 16 training patients. For the proliferation score, ridge models on Phikon-v2 features were trained on the 16 training patients of each fold, with and without the pairwise difference loss, to predict *MKI67*, *TOP2A*, *CENPF* and *PCLAF*. The score of a spot was the mean of the four $\log(1+N)$ values or of their predictions. Predicted hotspots were the 10% of spots with the highest predicted score in each patient, and hotspot enrichment was the mean measured score in these spots minus that in the remaining spots. Outcomes were the Pearson correlation between predicted and measured scores and the hotspot enrichment. In a sensitivity analysis, both scores were normalized for sequencing depth, and the ridge model was refitted to the normalized score.

*Single-cell methods.* Each method was trained in the standard, neighboring and non-neighboring conditions, and predictions were averaged over seeds. The nine breast cancer Xenium sections were the test data (Extended Data Fig. 7). Within each method, paired comparisons used the same malignant cells and neighborhoods, and the three proliferation outcomes used the same four genes (Supplementary Table 6). The four outcomes were the fine-scale RSC over all genes, the RSC of the proliferation score between neighborhoods, the pairwise accuracy of the score between neighborhood centers no more than 64 $\mu$m apart and hotspot enrichment. For these outcomes, the measured score was normalized over the genes shared by the method and the Xenium panel.

#### Supplementary Note 1: Class-mean baseline

*HER2ST.* The class-mean baseline (Methods) had a whole-section Pearson $r$ of 0.111 and ranked sixth of the 12 sets of predictions, above six of the 11 published methods (median 0.108; highest 0.144). All values were computed with the same gene-selection rule. Recomputing whole-section Pearson $r$ of all sets of predictions with another gene-selection rule changed values by less than 0.004 and left the ranks unchanged. Because all within-region predictions of this baseline are tied, its normalized pairwise accuracy is given with half credit for tied predictions and with tied predictions counted as errors, the rule used for the published methods. For unmatched pairs it was 22.4% and $-128\%$ (median of the published methods, 28.5%). For matched pairs, which can lie in two annotated regions, it was 6.8% and $-288\%$ (median of the published methods, 6.7%). Within annotated regions it was 1.5% and $-399\%$, last of the 12 sets of predictions under both rules (median of the published methods, 6.6%). With half credit, the median of the published methods exceeded the class-mean baseline within regions by 5.1 percentage points (95% CI, 2.8 to 8.3). The published methods had at most 0.012% tied pairs.

For an aggregation comparison, spots were grouped into non-overlapping blocks of $k \times k$ spots ($k = 2, 3, 4$) on the array grid, and predictions and $\log(1 + N)$ of the unsplit counts were averaged within each block. Agreement was the gene-wise Pearson correlation across the blocks of a section, averaged over genes and then over patients. With this definition, the class-mean baseline had a correlation of 0.125 for single spots and 0.245 for blocks of $4 \times 4$ spots, fourth and third of the 12 sets of predictions. The values for EGNv2 were 0.146 and 0.258. A class-mean baseline built from every annotated region, instead of tumor and other tissue, had a whole-section Pearson $r$ of 0.098 and gene-wise correlations of 0.106 for single spots and 0.218 for $4 \times 4$ blocks.

*Xenium.* On the Xenium cells and genes of sCellST, the class-mean baseline had a fine-scale RSC over all cells of 0.081 at single-cell resolution and 0.270 after aggregation into 256-$\mu$m bins, compared with 0.036 and 0.218 for sCellST. Within malignant cells, the baseline is constant and has no RSC. A class-mean baseline built from every CellViT class had a fine-scale RSC over all cells of 0.116 at single-cell resolution and 0.352 after aggregation into 256-$\mu$m bins.

*Kidney and lung cancer sections.* The class-mean baseline separated annotated TLS from immune-infiltrated, tumor and normal spots with a mean AUROC of 0.994 to 1.000 per section, compared with 0.925 to 0.994 for the Ridge–Phikon-v2 predictions and 0.614 to 0.709 for measured expression. Its RSC was 0.556 in lung and 0.523 in kidney over whole sections, compared with 0.493 and 0.345 for Ridge–Phikon-v2, and 0.827 and 0.730 between regions, compared with 0.793 and 0.593. Within regions it is constant and has no RSC.

#### Supplementary Note 2: Derivation of the reproducible-signal correlation

*Model.* For gene $g$ in bin, spot or cell $i$, the count $N_{gi}$ follows a Poisson distribution with rate $\lambda_{gi}$, the expected number of captured molecules. Under binomial splitting with probability 0.5, the two halves $N^A_{gi}$ and $N^B_{gi}$ are independent Poisson counts[19,21] with rate $\lambda_{gi}/2$. With $f$ the count transform of a dataset, the transformed halves are $A_{gi} = f(N^A_{gi})$ and $B_{gi} = f(N^B_{gi})$. Their common expectation, $S_{gi} = \mathrm{E}[f(N^A_{gi}) \mid \lambda_{gi}] = \mathrm{E}[f(N^B_{gi}) \mid \lambda_{gi}]$, is the expected expression. The halves can therefore be written as $A = S + \varepsilon_A$ and $B = S + \varepsilon_B$, where the sampling noise of each half has mean zero and the noise of one half is independent of the noise of the other.

*Expected inner products.* Centering, gene scaling and the scale operators are linear, and $\langle \cdot, \cdot \rangle_b$ is bilinear. We treat the gene scaling factors as fixed and take expectations over the sampling noise, given the rates and a prediction $\hat{Y}$ made without the counts it is evaluated against. Because $\hat{Y}$ is independent of the noise and $\mathrm{E}\langle \varepsilon_A, \varepsilon_B \rangle_b = 0$,

$$\mathrm{E}\langle \hat{Y}, A \rangle_b = \mathrm{E}\langle \hat{Y}, B \rangle_b = \langle \hat{Y}, S \rangle_b, \qquad \mathrm{E}\langle A, B \rangle_b = \langle S, S \rangle_b.$$

In contrast, $\mathrm{E}\langle A, A \rangle_b = \langle S, S \rangle_b + \mathrm{E}\langle \varepsilon_A, \varepsilon_A \rangle_b$ includes the variance of the sampling noise, and the same holds for $B$. The mean covariance $C$ therefore estimates $\langle \hat{Y}, S \rangle_b$, and $V_{\mathrm{RNA}}$ estimates $\langle S, S \rangle_b$, the variance of the expected expression at scale $b$. RSC combines these estimates,

$$\mathrm{RSC}_b = \frac{C}{\sqrt{V_{\mathrm{model}} V_{\mathrm{RNA}}}} \approx \frac{\langle \hat{Y}, S \rangle_b}{\sqrt{\langle \hat{Y}, \hat{Y} \rangle_b \, \langle S, S \rangle_b}} = r_b(\hat{Y}, S),$$

the correlation between the prediction and the expected expression at scale $b$. The approximation arises because RSC is a ratio of estimates, and its deviation is small when $V_{\mathrm{RNA}}$ is large relative to its sampling variation. The tests with the expected expression as a prediction measure this deviation over the full computation of each dataset (Methods). Because $V_{\mathrm{RNA}}$ is an estimate, RSC can exceed 1 when the reproducible signal at a scale is small, and negative values are kept.

*Relation to normalized correlation.* With two halves, RSC equals the normalized correlation coefficient used to evaluate models of neural responses over repeated trials[42,43]. In that coefficient, the covariance between the prediction and the mean response is divided by the square roots of the prediction variance and of the signal power, which is estimated from the covariance between trials. Both are correlations corrected for attenuation[23].

*Correlation with the mean of the halves.* The two halves have the same distribution given the rates, and $V_A$ and $V_B$ have the same expectation. For halves of equal variance, $V_A = V_B = V$, the split-half reliability is $\rho_b = V_{\mathrm{RNA}}/V$, and the mean correlation of the prediction with the two halves is

$$\tfrac{1}{2}\left\{ r_b(\hat{Y}, A) + r_b(\hat{Y}, B) \right\} = \frac{C}{\sqrt{V_{\mathrm{model}} V}} = \mathrm{RSC}_b \sqrt{\rho_b}.$$

For the mean of the halves, $M = (A + B)/2$, the inner product with the prediction is $\langle \hat{Y}, M \rangle_b = C$, and $V_M = \langle M, M \rangle_b = (V_A + V_B + 2V_{\mathrm{RNA}})/4 = V(1 + \rho_b)/2$. This gives equation (2) of the main text,

$$r_b(\hat{Y}, M) = \frac{C}{\sqrt{V_{\mathrm{model}} V_M}} = \mathrm{RSC}_b \sqrt{\frac{2\rho_b}{1 + \rho_b}}.$$

Used as a prediction, the expected expression $\hat{Y} = S$ has an RSC near 1 and a correlation with the mean of the halves of about $\sqrt{2\rho_b/(1+\rho_b)}$. The correlation of any prediction with the mean of the halves is therefore approximately the product of its correlation with the expected expression and the correlation of the expected expression with the mean of the halves. The same relation links the Fourier shell correlation (FSC) between two cryo-electron microscopy half-maps to the correlation of the full map with the structure without noise[52], $C_{\text{ref}} = \sqrt{2\,\text{FSC}/(1+\text{FSC})}$.

*Limit without sampling noise.* As the sampling noise approaches zero, $\rho_b$ and $\sqrt{2\rho_b/(1+\rho_b)}$ approach 1, and RSC approaches the correlation of the prediction with the mean of the halves. At $N_{\text{half}} = N/2$, the half-count transforms of this study equal the corresponding transforms of the full count, as $\log(1+2N_{\text{half}}) = \log(1+N)$ and $\sqrt{N_{\text{half}}/(0.5a)} = \sqrt{N/a}$. In this limit, RSC therefore equals the correlation of the prediction with the transformed measured counts. For the drawn counts of the tests with the expected expression, the correlation with the transformed unsplit counts deviated from the correlation with the mean of the halves by at most 0.014 (Methods).

*Overdispersion.* In negative binomial count models, counts vary more than Poisson counts around the tissue expression. In such models, the rate $\lambda_{gi}$ contains extra-Poisson variation in addition to the tissue expression. This variation arises before the split and is shared by the two halves. For a prediction that is independent of this variation, the variation adds to $V_{\text{RNA}}$ and not to the expectation of $C$, which raises the split-half reliability and lowers RSC. This direction agrees with the semi-synthetic test in HEST, in which predictions equal to the transformed latent rates of a beta-binomial count model had an RSC below 1 at every scale (Methods).

*Correlation without correction for attenuation.* For comparison, we computed the correlation with the mean of the two transformed halves, $r_{\text{mean}} = C/\sqrt{V_{\text{model}}V_M}$. On all malignant cells of each method at fine scales, this correlation was 0.009 for sCellST and 0.017 for DeepSpot2Cell, compared with RSCs of 0.016 and 0.031.

#### Supplementary Note 3: Tertiary lymphoid structures

*Kidney and lung cancer sections.* Within TLS, expression was more reproducible between structures than within them. Between structures, the split-half reliability was 0.509 in lung and 0.610 in kidney (KC3), and the predictions had RSCs of 0.364 (95% interval, 0.147 to 0.573) and 0.331 (0.148 to 0.522). Within structures, the split-half reliability was 0.102 and 0.167, and RSC was 0.183 (0.054 to 0.315) and 0.238 (0.014 to 0.394). With the pairwise difference loss, RSC within TLS was 0.235 in lung and 0.246 in kidney, compared with 0.253 and 0.293 after standard training, and RSC between regions was 0.754 and 0.551, compared with 0.793 and 0.593.

*Training within one cancer type.* With training only on the other section of the same cancer type, mean AUROC for annotated TLS versus other spots was 0.869 and 0.734 in KC1 and KC3 and 0.878 and 0.700 in LC1 and LC3. Over whole sections, RSC was 0.250 (95% interval, 0.153 to 0.402) in kidney and 0.283 (0.169 to 0.388) in lung, compared with 0.345 and 0.493 after training on both cancer types. Between regions, RSC was 0.613 (0.518 to 0.693) in kidney and 0.644 (0.530 to 0.720) in lung, compared with 0.593 and 0.793. Within TLS, RSC was 0.248 (0.167 to 0.321) in kidney (KC3) and 0.275 (0.182 to 0.398) in lung, compared with 0.293 and 0.253. Each of these models was trained on a single section.

*Section selection.* LC2 showed a positive but uncertain lymphocyte enrichment at annotated TLS (AUROC, 0.638; 95% interval, 0.371 to 0.845) and was therefore not part of the main analysis. With LC2 included, the lung predictions reached an RSC of 0.730 (0.693 to 0.759) between regions, compared with 0.844 for the class-mean baseline, and 0.224 (0.096 to 0.324) within TLS. Between TLS structures, RSC fell to 0.083 (−0.044 to 0.227), because the predictions did not order the structures of LC2.

#### Supplementary Note 4: Training comparisons

*Models other than TCGN.* For the ten models other than TCGN, the pairwise difference loss raised the mean fine-scale RSC from 0.197 to 0.207 (change, +0.010; 95% CI, 0.008 to 0.014; exact $P = 0.0039$), with gains in all nine samples and positive model means in 9 of 10 models.

*Neighboring versus non-neighboring pairs at broader scales.* Relative to non-neighboring pairs, neighboring pairs increased the broad-scale RSC by 0.003 (95% CI, 0.0001 to 0.006) and whole-section Pearson $r$ by 0.004 (95% CI, 0.0006 to 0.007), each in five of nine samples. The fine-scale gain of neighboring pairs exceeded the broad-scale gain by 0.006 (95% CI, 0.003 to 0.009; exact sign-flip $P = 0.14$).

*ccRCC cohort.* For an MLP trained on 16 patients, the pairwise difference loss raised the Pearson correlation between predicted and measured changes in expression between neighboring spots by 0.0076 in 24 ccRCC test patients (95% CI, 0.0041 to 0.0118; Supplementary Fig. 2). With standard training, increasing the number of training patients from 1 to 16 raised the mean gene-wise Pearson correlation from 0.021 to 0.158. In the same cohort, a ridge model trained with the loss gave a higher Pearson correlation of the proliferation score (0.197 compared with 0.187; change, +0.0102; 95% CI, 0.0050 to 0.0159) and a higher hotspot enrichment (0.0749 compared with 0.0702; change, +0.0047; 95% CI, 0.0009 to 0.0085). After both scores were normalized for sequencing depth and the ridge model was refitted to the normalized score, the loss no longer raised either outcome (change in Pearson correlation, +0.003, with a 95% CI including zero; change in hotspot enrichment, −0.0001).

*Encoder replacement in sCellST.* Cells, genes, data splits, training procedure and evaluation were held fixed while the image encoder of sCellST was changed from ResNet50 to Phikon-v2, with the four outcomes used for the single-cell training comparisons (Methods). The fine-scale RSC changed by +0.014 (95% CI, −0.002 to 0.031) and hotspot enrichment by −0.173 (95% CI, −0.339 to −0.024). No change was significant after Holm adjustment (Supplementary Table 6).

#### Supplementary Note 5: Sensitivity analyses for pairwise accuracy

*HEST models with higher whole-section correlation.* The decline after matching also occurred for Ridge–Phikon-v2 and the MLP, which had whole-section Pearson correlations of 0.509 and 0.517 with the three cancer types weighted equally. Matching reduced their gains over the baseline by 22.8 and 23.6 percentage points (Supplementary Fig. 1b).

*Depth-normalized measured directions.* With measured directions taken from depth-normalized counts (Methods), 6.4 to 6.8% of HER2ST comparisons and 6.1 to 6.6% of HEST comparisons changed direction. Normalized pairwise accuracy in HER2ST still fell after matching and within annotated regions, the agreement of rankings with whole-section Pearson $r$ still fell from unmatched to within-region pairs, matching still reduced the model gain in HEST ($P = 0.0039$), and the gains in pairwise accuracy between neighboring spots from neighboring pairs remained positive (Supplementary Table 2).

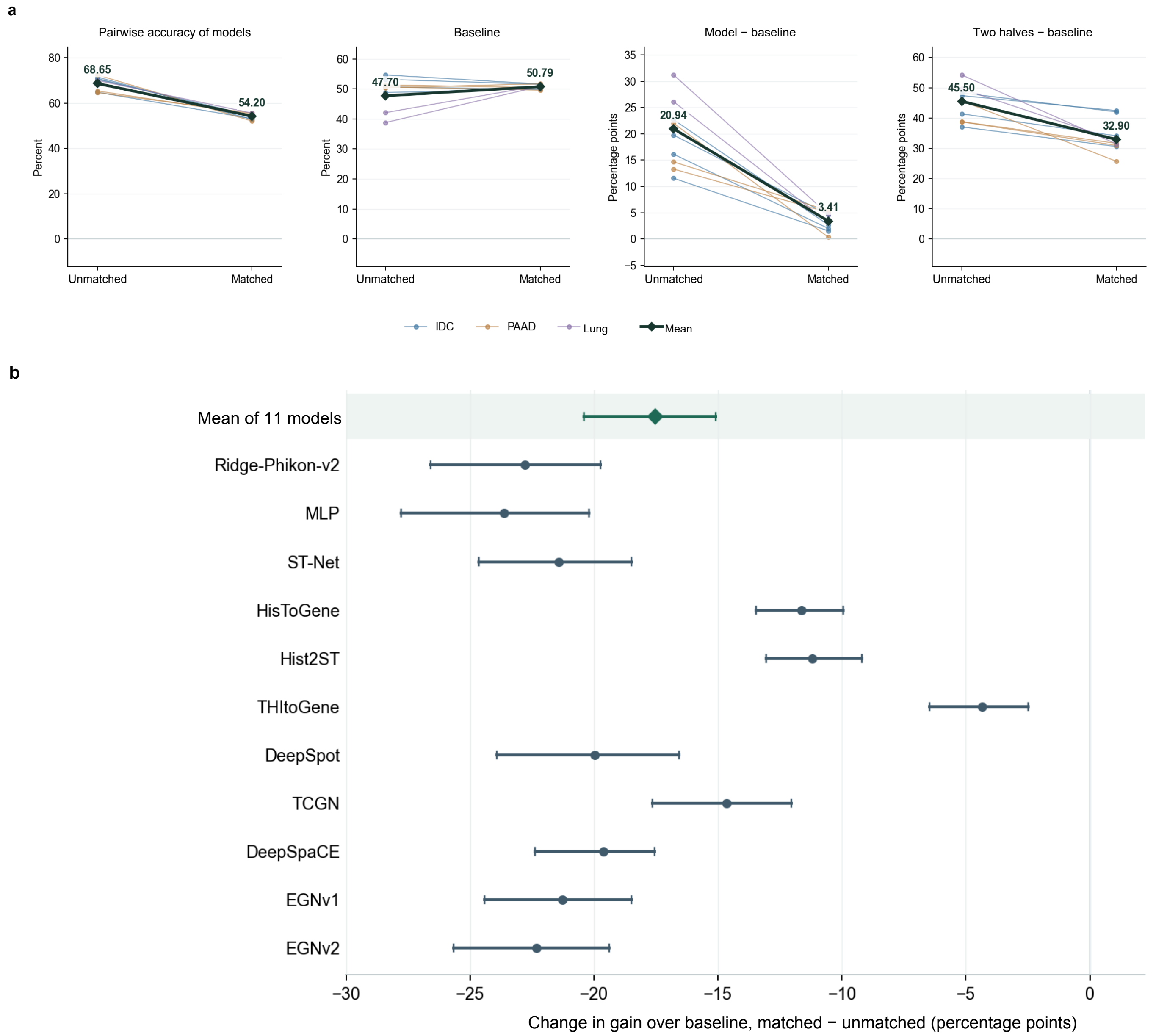


**Supplementary Fig. 1 | Matching for expression profile and total count removes most of the pairwise accuracy above the majority-order baseline in Xenium samples of breast, pancreatic and lung cancer. a,** Pairwise accuracy of the models, accuracy of the majority-order baseline, and the gains of the models and of the two halves over the baseline, for unmatched and matched spot pairs. Means are 68.65% and 54.20% for model accuracy, 20.94 and 3.41 percentage points for the model gain, and 45.50 and 32.90 percentage points for the gain of the two halves. Thin lines, samples colored by cancer type; dark line, mean. **b,** Change in the model gain from unmatched to matched pairs, for the mean of 11 models (−17.53 percentage points; 95% CI −20.39 to −15.08; exact two-sided $P = 0.0039$) and for each model. All 11 model means decrease, and 10 of 11 have Holm-adjusted $P < 0.05$; model-specific $P$ values are from exact sign-flip tests over all 512 sample-level sign patterns, Holm-adjusted across the 11 models. Bars, 95% percentile intervals from 10,000 bootstrap draws of samples within cancer type. Gains are in percentage points of accuracy and are not divided by the gain of the two halves. $n = 9$ samples (four IDC, three PAAD and two lung).

**Supplementary Table 1.** Normalized pairwise accuracy in HER2ST and changes between pair sets.

| Pairs | Median gain | 95% CI |
|---|---|---|
| Unmatched | 28.5 | [15.6, 40.3] |
| Matched | 6.7 | [3.1, 10.9] |
| Within regions | 6.6 | [3.2, 11.0] |

| Contrast | Change | 95% CI | Exact $P$ | Holm $P$ |
|---|---|---|---|---|
| Matched − Unmatched | −21.89 | [−31.33, −11.34] | 0.0078125 | 0.0234375 |
| Within regions − Matched | −0.05 | [−2.40, +1.94] | 0.953125 | 0.953125 |
| Within regions − Unmatched | −21.95 | [−32.22, −10.81] | 0.015625 | 0.03125 |

Each value is the median across 11 methods of their means over eight patients. Intervals are 95% percentile intervals from 20,000 bootstrap draws of patients. Exact two-sided paired tests enumerate all 256 patient-level label swaps, with Holm adjustment over the three contrasts.

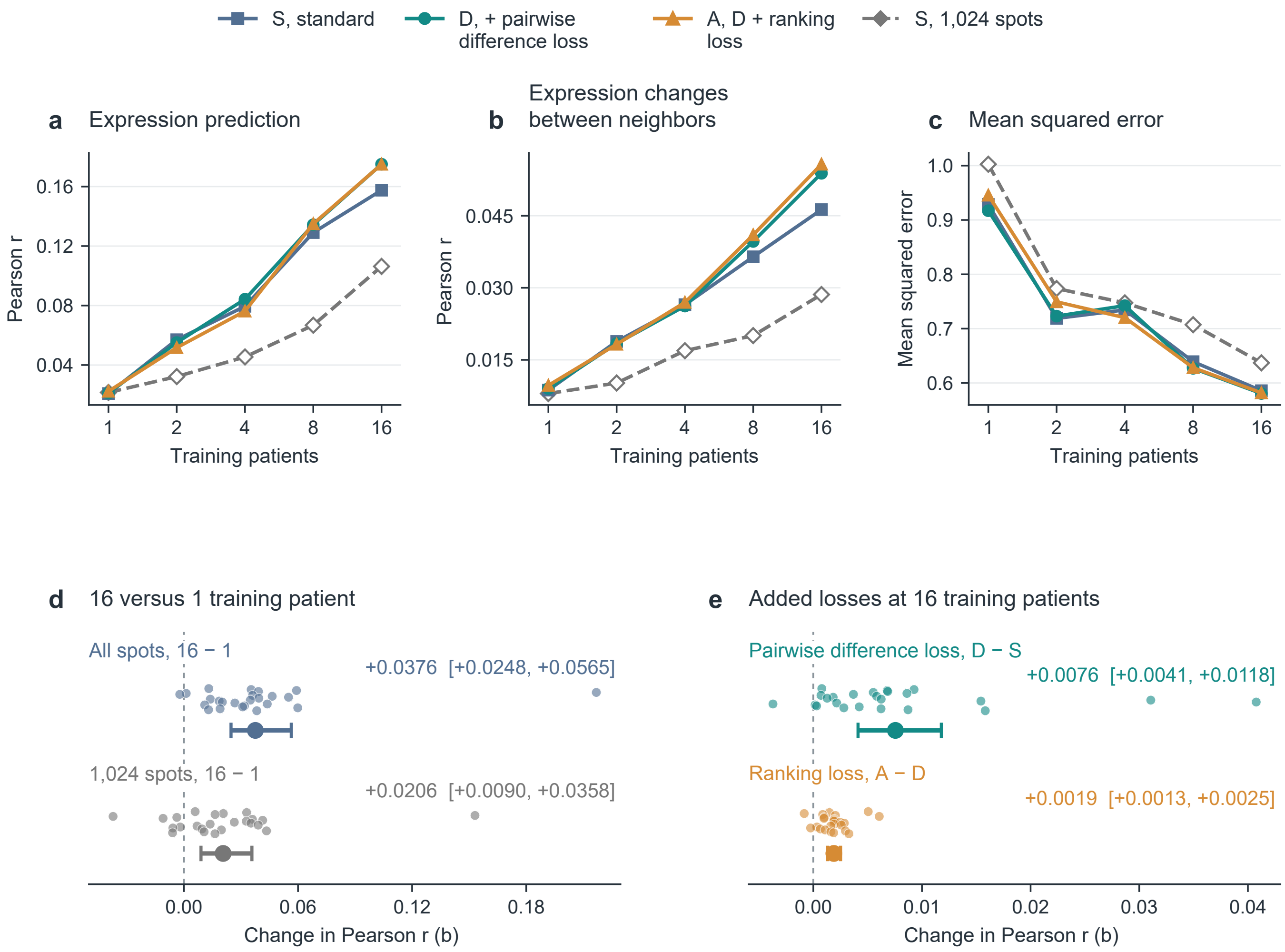


**Supplementary Fig. 2 | Effects of the number of training patients and of added losses in ccRCC. a–c,** Pearson correlation between predictions and measured expression (**a**), Pearson correlation between predicted and measured changes in expression between neighboring spots (**b**) and mean squared error (**c**) in 24 ccRCC test patients, for MLP models trained on 1, 2, 4, 8 or 16 patients. Models were trained with the standard loss (S), with the pairwise difference loss added (D), with a ranking loss added to D (A), or with the standard loss on 1,024 training spots in total and a fixed number of updates. **d,** Change in the correlation of **b** from 1 to 16 training patients, with all spots or with 1,024 spots. **e,** Changes at 16 training patients from adding the pairwise difference loss (D − S) and the ranking loss (A − D). Points, patient-level changes; large points, means with patients weighted equally; bars, 95% bootstrap intervals over patients.

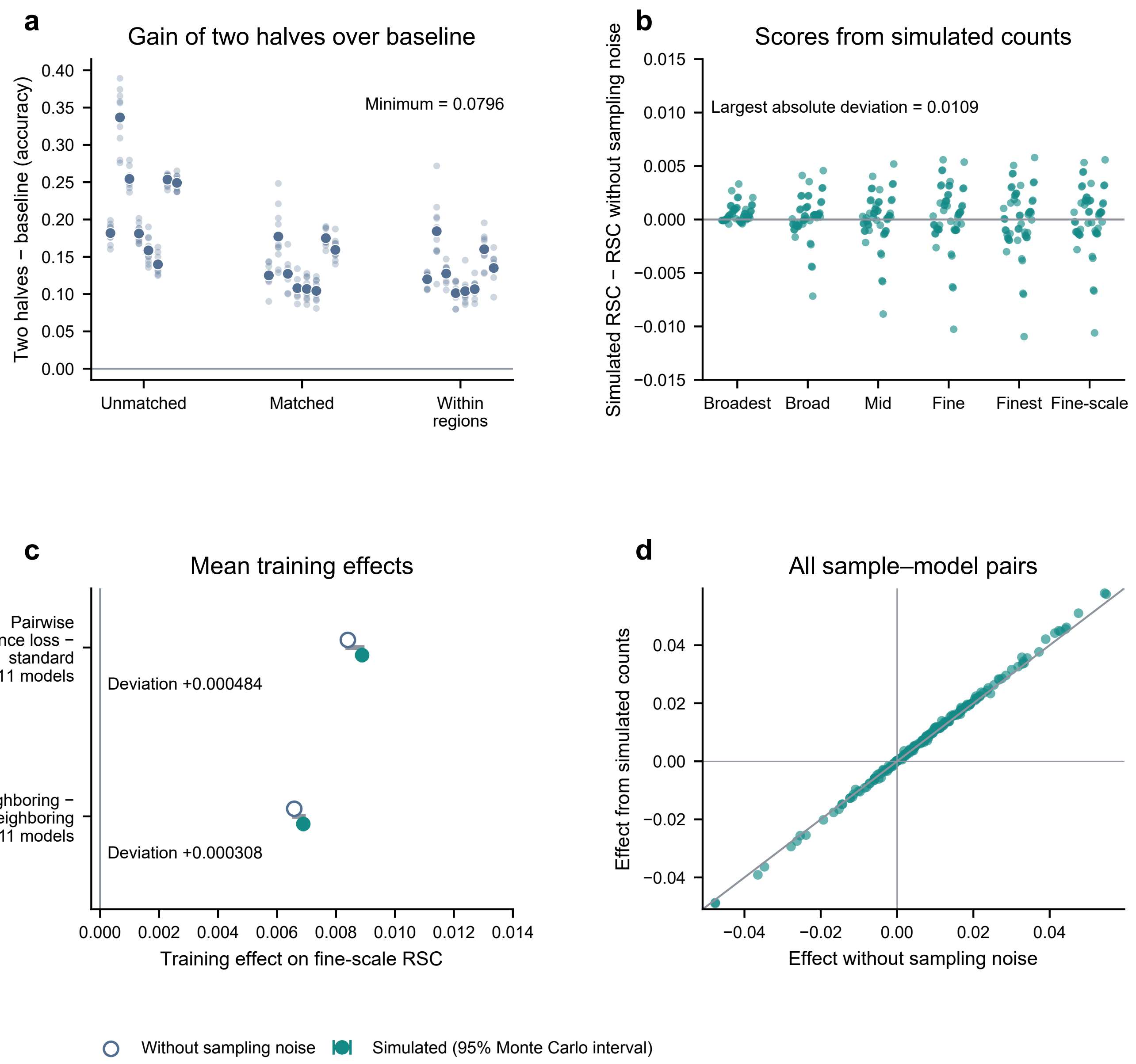


**Supplementary Fig. 3 | Checks of the denominator of the normalized pairwise accuracy and of the reproducible-signal correlation (RSC). a,** Gain of the two halves over the majority-order baseline, the denominator of the normalized pairwise accuracy, for 240 combinations of eight HER2ST patients, three pair sets and ten count splits; larger points, patient means. All values are positive (minimum 0.0796). **b,** Deviation of the reproducible-signal correlation computed from simulated counts from its value without sampling noise, for synthetic predictions of six target correlations in each of nine Xenium samples of breast, pancreatic and lung cancer, at five scales and for the fine-scale RSC (324 conditions). The largest absolute deviation is 0.0109. **c,** Mean training effects of 11 models on the fine-scale RSC when the scores are recomputed from simulated counts, for the pairwise difference loss compared with standard training (top) and neighboring compared with non-neighboring pairs (bottom). Open circles, values without sampling noise; filled circles, simulated estimates with 95% Monte Carlo intervals. Both mean effects are positive, and they deviate from the values without sampling noise by +0.000484 and +0.000308. **d,** Effects from simulated counts versus effects without sampling noise for all 198 pairs of sample and model.

**Supplementary Table 2.** Pairwise accuracy with measured directions from depth-normalized counts.

**a, HER2ST normalized pairwise accuracy and agreement with rankings by whole-section Pearson *r*.**

| Pairs | *G* with raw-count directions (%) | *G* with depth-normalized directions (%) | Raw $\tau_b$ | Normalized $\tau_b$ |
|---|---|---|---|---|
| Unmatched | 28.5433 | 29.0475 | 0.8182 | 0.6727 |
| Matched | 6.6507 | 6.3710 | 0.1273 | 0.0545 |
| Within-region | 6.5966 | 4.6071 | 0.0909 | −0.0545 |

**b, HEST pairwise accuracy on common paired strata.**

| Pairs | Model gain raw (pp) | Model gain normalized (pp) | Gain of the two halves, raw (pp) | Gain of the two halves, normalized (pp) |
|---|---|---|---|---|
| Unmatched | 20.9421 | 18.1072 | 45.5004 | 40.8808 |
| Matched | 3.4105 | 2.8524 | 32.8996 | 30.5130 |

| Matched minus unmatched | Change (pp) | 95% CI | *P* |
|---|---|---|---|
| Raw counts | −17.5315 | [−20.3920, −15.0793] | 0.00390625 |
| Depth-normalized | −15.2548 | [−18.1314, −12.8079] | 0.00390625 |

**c, Pairwise accuracy gains from neighboring pairs.**

| Pairs | Contrast | Raw (pp), 95% CI | Normalized (pp), 95% CI |
|---|---|---|---|
| All neighbors | Neighboring − standard | 0.4199 [0.3611, 0.4796] | 0.3766 [0.3218, 0.4368] |
| All neighbors | Neighboring − non-neighboring | 0.1245 [0.0327, 0.2225] | 0.1009 [0.0205, 0.1912] |
| Depth ratio ≤ 1.25 | Neighboring − standard | 0.2744 [0.2006, 0.3519] | 0.2660 [0.1915, 0.3434] |
| Depth ratio ≤ 1.25 | Neighboring − non-neighboring | 0.1079 [0.0257, 0.1915] | 0.1056 [0.0236, 0.1885] |

The same pairs, predictions and count splits were used for both sets of measured directions. HER2ST values are medians across 11 methods in eight patients. In **b,** HEST values average 11 models and nine samples, with cancer types weighted equally. In **c,** models are weighted equally within samples, and samples are weighted equally. The Kendall $\tau_b$ between the HEST model ranks for unmatched and matched pairs is 0.927 with either set of measured directions.

**Supplementary Table 3.** Broadest-to-finest declines in reproducible-signal correlation (RSC).

| Prediction set | Units | Mean ΔRSC | Interval | Exact *P* | Holm *P* |
|---|---|---|---|---|---|
| Colorectal tissue, fixed 32-$\mu$m grid | 3 patients | +0.5021 | [+0.4739, +0.5202] | 0.25 | 0.375 |
| HEST, Ridge–Phikon-v2 | 9 samples | +0.3651 | [+0.3376, +0.3936] | 0.00390625 | 0.015625 |
| sCellST, malignant cells | 5 patients | +0.0254 | [−0.0003, +0.0505] | 0.1875 | 0.375 |
| DeepSpot2Cell, malignant cells | 5 patients | +0.0753 | [+0.0469, +0.1017] | 0.0625 | 0.1875 |
| GHIST, malignant cells | 5 patients | +0.0033 | [−0.0519, +0.0806] | 1 | — |
| HEST, MLP | 9 samples | +0.3751 | [+0.3410, +0.4103] | 0.00390625 | — |
| HEST, ST-Net | 9 samples | +0.3646 | [+0.3361, +0.3954] | 0.00390625 | — |
| HEST, HisToGene | 9 samples | +0.1674 | [+0.1243, +0.2112] | 0.00390625 | — |
| HEST, Hist2ST | 9 samples | +0.2401 | [+0.1889, +0.2942] | 0.00390625 | — |
| HEST, THItoGene | 9 samples | −0.0057 | [−0.0324, +0.0210] | 0.828125 | — |
| HEST, DeepSpot | 9 samples | +0.2907 | [+0.2513, +0.3310] | 0.00390625 | — |
| HEST, TCGN | 9 samples | +0.2256 | [+0.1949, +0.2566] | 0.00390625 | — |
| HEST, DeepSpaCE | 9 samples | +0.3346 | [+0.3008, +0.3685] | 0.00390625 | — |
| HEST, EGNv1 | 9 samples | +0.4074 | [+0.3698, +0.4436] | 0.00390625 | — |
| HEST, EGNv2 | 9 samples | +0.3540 | [+0.3216, +0.3884] | 0.00390625 | — |

The colorectal contrast is the scale of at least 4,096 $\mu$m minus the scale below 128 $\mu$m on the 32-$\mu$m grid. The HEST and malignant-cell contrasts are Broadest minus Finest; the malignant-cell contrasts use the cells matched across the three methods, as in Fig. 4a. Intervals are 95% percentile intervals from 20,000 bootstrap draws of patients or samples; for the three colorectal patients, this interval is the range of patient values. Exact two-sided sign-flip tests for the first five rows enumerate 8, 512, 32, 32 and 32 patterns, with Holm adjustment over the first four contrasts; the GHIST contrast is unadjusted. The last ten rows are the other HEST models of the pairwise-loss comparison under standard training, shown in Fig. 3d; their contrasts enumerate 512 patterns and are unadjusted.

**Supplementary Table 4.** Normalized pairwise accuracy of Ridge–Phikon-v2 and the cell-count model in 45 HEST-Benchmark patients.

| Pairs | Tied predictions | Ridge–Phikon-v2 | Cell-count model | Difference | Patients with Ridge–Phikon-v2 higher |
|---|---|---|---|---|---|
| Unmatched | Half credit | 49.1 [43.3, 54.7] | 33.9 [28.5, 39.4] | +15.2 [+10.4, +20.1] | 37/45 |
| Matched | Half credit | 19.9 [16.0, 23.9] | 11.3 [8.4, 14.1] | +8.6 [+5.7, +11.7] | 36/45 |
| Matched, expression only | Half credit | 25.3 [21.2, 29.6] | 16.5 [13.3, 19.7] | +8.8 [+5.7, +12.1] | 37/45 |
| Unmatched | Counted as errors | 49.1 | −8.9 [−29.9, +9.5] | +58.1 [+41.5, +77.3] | 43/45 |
| Matched | Counted as errors | 19.9 | −78.6 [−111.4, −49.6] | +98.5 [+69.8, +131.5] | 45/45 |
| Matched, expression only | Counted as errors | 25.3 | −55.4 [−82.0, −32.0] | +80.7 [+57.3, +107.1] | 42/45 |

Values are normalized pairwise accuracies in percent, and differences are in percentage points. Matched pairs were matched for expression profile and total count, as in the main analysis, or for expression profile only. The cell-count model gave tied predictions for 18% of matched pairs. Intervals are 95% percentile intervals from 20,000 bootstrap draws of patients.

**Supplementary Table 5.** Kidney and lung cancer sections: RNA-based checks and agreement of predictions for TLS-associated genes.

**a, Checks of the eight sections.**

| Section | TLS structures | Lymphocyte AUROC (95% interval) | Alignment | Use |
|---|---|---|---|---|
| KC1 | 6 | 0.865 (0.850–0.945) | As recorded | Main analysis |
| KC2 | 5 | 0.763 (0.573–0.852) | Rotated by 180° | Excluded |
| KC3 | 21 | 0.897 (0.835–0.937) | As recorded | Main analysis |
| LC1 | 15 | 0.796 (0.705–0.840) | As recorded | Main analysis |
| LC2 | 11 | 0.638 (0.371–0.845) | As recorded | Sensitivity analysis |
| LC3 | 28 | 0.721 (0.671–0.762) | As recorded | Main analysis |
| LC4 | 16 | 0.537 (0.439–0.649) | As recorded | Excluded |
| LC5 | 6 | 0.325 (0.206–0.389) | As recorded | Excluded |

**b, Mean AUROC of the eight genes for annotated TLS versus other annotated spots.**

| Section | Standard training (95% interval) | Pairwise difference loss | Class-mean baseline | Measured expression |
|---|---|---|---|---|
| KC1 | 0.947 (0.866–0.972) | 0.930 | 0.997 | 0.637 |
| KC3 | 0.925 (0.896–0.945) | 0.924 | 1.000 | 0.709 |
| LC1 | 0.994 (0.984–0.997) | 0.991 | 0.994 | 0.637 |
| LC3 | 0.932 (0.911–0.948) | 0.933 | 1.000 | 0.614 |

**c, Reproducible-signal correlation (RSC) by term.**

| Cancer type | Term | Split-half reliability | RSC, standard training (95% interval) | RSC, pairwise difference loss | RSC, class-mean baseline (95% interval) |
|---|---|---|---|---|---|
| Lung | Whole section | 0.108 | 0.493 (0.401–0.564) | 0.472 | 0.556 (0.496–0.607) |
| | Between regions | 0.959 | 0.793 (0.769–0.808) | 0.754 | 0.827 (0.784–0.862) |
| | Within regions | 0.062 | 0.128 (0.067–0.195) | 0.119 | — |
| | Within TLS | 0.149 | 0.253 (0.146–0.400) | 0.235 | — |
| | Between structures | 0.509 | 0.364 (0.147–0.573) | 0.360 | — |
| | Within structures | 0.102 | 0.183 (0.054–0.315) | 0.159 | — |
| Kidney | Whole section | 0.153 | 0.345 (0.253–0.440) | 0.319 | 0.523 (0.439–0.612) |
| | Between regions | 0.947 | 0.593 (0.520–0.652) | 0.551 | 0.730 (0.685–0.803) |
| | Within regions | 0.144 | 0.191 (0.138–0.250) | 0.168 | — |
| | Within TLS | 0.281 | 0.293 (0.144–0.427) | 0.246 | — |
| | Between structures | 0.610 | 0.331 (0.148–0.522) | 0.287 | — |
| | Within structures | 0.167 | 0.238 (0.014–0.394) | 0.183 | — |

In **a**, the lymphocyte score is the mean of $\log(1+10{,}000N/\sum_g N)$ over *PTPRC*, *CD52*, *CD3E*, *CD2* and *CD79A*, and its AUROC compares annotated TLS with tumor and normal spots, with intervals from 2,000 resamples of TLS structures and the other spots fixed. Alignment is 'as recorded' when none of 21 shifts, rotations and mirror images raised the variance of expression explained by the annotation by more than 0.01 (largest increase, 0.0085 in KC1); in KC2, a rotation by 180° raised it by 0.415. KC2 and LC4 were also excluded in the original study. In **b**, the class-mean baseline is constant within each class and its AUROC therefore has no resampling interval. In **c**, lung is LC1 and LC3 and kidney is KC1 and KC3; the terms within regions of kidney are from KC3. Split-half reliability is computed for each term. Intervals are 95% percentile intervals from 2,000 resamples of TLS structures for the terms within TLS and of blocks of $10 \times 10$ spots for the other terms. The class-mean baseline is constant within each annotated class and has no RSC within regions.

**Supplementary Table 6.** Changes in fine-scale reproducible-signal correlation (RSC) and in proliferation outcomes in five training comparisons of sCellST and DeepSpot2Cell.

| Outcome | Mean Δ | 95% CI | Exact $P$ |
|---|---|---|---|
| *DeepSpot2Cell: neighboring − standard* | | | |
| Fine-scale RSC (all genes) | +0.0027 | [+0.0012, +0.0040] | 0.0625 |
| Reproducible-signal correlation of the proliferation score | −0.0745 | [−0.2116, +0.0307] | 0.4375 |
| Pairwise accuracy of the proliferation score (percentage points) | −0.741 | [−1.475, +0.066] | 0.25 |
| Hotspot enrichment | −0.0713 | [−0.3129, +0.1223] | 0.6875 |
| *DeepSpot2Cell: neighboring − non-neighboring* | | | |
| Fine-scale RSC (all genes) | +0.0025 | [+0.0009, +0.0049] | 0.0625 |
| Reproducible-signal correlation of the proliferation score | −0.0554 | [−0.1941, +0.0504] | 0.6875 |
| Pairwise accuracy of the proliferation score (percentage points) | −0.609 | [−1.318, +0.101] | 0.3125 |
| Hotspot enrichment | −0.0633 | [−0.3371, +0.1553] | 0.75 |
| *sCellST: neighboring − standard* | | | |
| Fine-scale RSC (all genes) | −0.0035 | [−0.0075, +0.0005] | 0.25 |
| Reproducible-signal correlation of the proliferation score | +0.0136 | [−0.0204, +0.0435] | 0.5 |
| Pairwise accuracy of the proliferation score (percentage points) | +0.179 | [−0.354, +0.713] | 0.625 |
| Hotspot enrichment | +0.0100 | [−0.0134, +0.0357] | 0.5625 |
| *sCellST: neighboring − non-neighboring* | | | |
| Fine-scale RSC (all genes) | −0.0053 | [−0.0086, −0.0014] | 0.125 |
| Reproducible-signal correlation of the proliferation score | +0.0001 | [−0.0457, +0.0377] | 1 |
| Pairwise accuracy of the proliferation score (percentage points) | −0.071 | [−0.375, +0.197] | 0.8125 |
| Hotspot enrichment | +0.0215 | [−0.0291, +0.0900] | 0.625 |
| *sCellST: Phikon-v2 − ResNet50* | | | |
| Fine-scale RSC (all genes) | +0.0139 | [−0.0015, +0.0311] | 0.25 |
| Reproducible-signal correlation of the proliferation score | −0.1620 | [−0.4443, +0.0584] | 0.375 |
| Pairwise accuracy of the proliferation score (percentage points) | +0.052 | [−1.690, +2.164] | 0.9375 |
| Hotspot enrichment | −0.1728 | [−0.3394, −0.0237] | 0.125 |

Changes are the first-named condition minus the second, with sections averaged within patients and the five patients weighted equally. Intervals are 95% percentile intervals from 20,000 bootstrap draws of patients. Exact two-sided sign-flip tests enumerate all 32 patient sign patterns; all 20 Holm-adjusted $P$ values are 1, and the smallest attainable unadjusted two-sided $P$ with five patients is 0.0625.

**Supplementary Table 7.** Datasets, sections and patients.

| Dataset | Group | Sections or samples |
|---|---|---|
| HER2ST | Patients A–H | A1, B1, C1, D1, E1, F1, G2, H1 |
| Colorectal cancer | Discovery patients P1–P3<br>Fourth patient P5 | P1CRC, P2CRC, P3NAT<br>P5CRC, P5NAT (analyzed separately and then averaged) |
| HEST | IDC<br>PAAD<br>Lung | TENX99, TENX95, NCBI783, NCBI785<br>TENX116, TENX140, TENX126<br>TENX118, TENX141 |
| Breast cancer Xenium | P1<br>P2<br>P3<br>P4<br>P5<br>Training section for sCellST and DeepSpot2Cell | TENX98, TENX99<br>TENX95, TENX97<br>TENX94, TENX96<br>NCBI784, NCBI785<br>NCBI783<br>TENX39 |
| ccRCC | 24 patients | INT1–INT24; six outer folds of four test patients each |
| HEST-Benchmark | Nine tasks, 45 patients | All 70 samples: IDC 4, PRAD 23, PAAD 3, SKCM 2, COAD 4, READ 4, CCRCC 24 (INT1–INT24), LUNG 2, LYMPH_IDC 4; patient folds of the benchmark |
| Kidney and lung cancer | Kidney<br>Lung | KC1, KC3 (main analysis); KC2 (excluded)<br>LC1, LC3 (main analysis); LC2 (sensitivity analysis); LC4, LC5 (excluded) |

The four HEST IDC samples are also among the breast cancer Xenium sections. NCBI783, NCBI784 and NCBI785 are from Janesick et al.[62]. The other samples and sections are 10x Genomics datasets (`https://www.10xgenomics.com/datasets`), TENX94 and TENX95 from *FFPE Human Breast with Pre-designed Panel*, TENX96 and TENX97 from *FFPE Human Breast with Custom Add-on Panel*, TENX98 and TENX99 from *FFPE Human Breast using the Entire Sample Area*, TENX116 from *Pancreatic Cancer with Xenium Human Multi-Tissue and Cancer Panel*, TENX126 from *FFPE Human Pancreas with Xenium Multimodal Cell Segmentation*, TENX140 from *FFPE Human Pancreatic Ductal Adenocarcinoma Data with Human Immuno-Oncology Profiling Panel*, TENX118 from *Preview Data: FFPE Human Lung Cancer with Xenium Multimodal Cell Segmentation*, TENX141 from *FFPE Human Lung Cancer Data with Human Immuno-Oncology Profiling Panel and Custom Add-on*, and TENX39 from *Human Breast Cancer: Ductal Carcinoma In Situ, Invasive Carcinoma (FFPE)*.
